# A predictive theory of experimental design for inferring neural population geometry in large-scale recordings

**DOI:** 10.64898/2026.07.19.739385

**Authors:** Itamar Daniel Landau, Gabriel C. Mel, Daniel J. O’Shea, Mark J. Schnitzer, Surya Ganguli

## Abstract

Ongoing technological advances will lead to recordings with progressively increasing numbers of neurons, while trial counts may only increase modestly. The analysis of such large-scale data increasingly relies on extracting collective neural population geometry. These combined recording and analysis trends raise the fundamental need for a predictive theory of experimental design that can tell us how accurately we will be able to infer such geometry in future larger scale recordings with more neurons and trials, by extrapolating from past smaller recordings. We derive such a theory for the simplest and most widely used method for extracting population geometry: principal component analysis. Our theory can predict how the dimensionality of neural data will grow with more neurons and trials and how accurate and reliable neuronal correlations and individual neural modes of the population geometry will be. Importantly, we find a blessing of dimensionality in which recording more neurons allows population geometry to be inferred accurately with fewer trials. The need for fewer trials in larger recordings will allow for the design of new experiments with more diverse trial types. Moreover, we derive scaling laws for the performance of neural prediction, setting the stage for the derivation of scaling laws for foundation models in neuroscience. We successfully test our theory across diverse species and recording modalities.

## INTRODUCTION

Over the last two decades, advances in recording technologies have revolutionized neuroscience. Techniques such as multi-beam two-photon calcium imaging Rumyantsev et al. (2020), volumetric calcium imaging Stringer et al. (2019), light-bead imaging Demas et al. (2021); Manley et al. (2024), high-density electrode recordings Jun et al. (2017); Steinmetz et al. (2021); Manley et al. (2024), high-density and high-field fMRI Uğurbil et al. (2013); Allen et al. (2022); Uğurbil (2021) now enable us to simultaneously measure hundreds, thousands, or even tens of thousands of neurons (or voxels) within a single recording session.

These large-scale recordings have motivated systems neuroscientists to move beyond the potentially inscrutable Rigotti et al. (2013) analysis of individual neurons and individual pairwise neural correlations, to the analysis of the coordinated neural population geometry of many neurons Ganguli and Sompolinsky (2012); Gao and Ganguli (2015); Saxena and Cunningham (2019); Chung and Abbott (2021); Jazayeri and Ostojic (2021); Perich et al. (2025). In such analyses, the activity pattern of *N* simultaneously recorded neurons (or voxels) on a single trial is thought of as a point in an *N* dimensional neural activity space. The collection of all such neural activity patterns then constitutes a point cloud in this space. The geometric structure of this point cloud often reveals fundamental insights into the nature of neural representations and computations across diverse subfields, from cognitive neuroscience Ebitz and Hayden (2021) to motor control Gallego et al. (2017) to the study of artificial neural networks Chung and Abbott (2021).

For example, recent work has used population geometry to expose constraints on efficient sensory representation Stringer et al. (2019) and the dimensionality of cortical codes Manley et al. (2024). In motor cortex, geometric analyses have revealed strategies for movement preparation under complex task demands Russo et al. (2018); Zimnik and Churchland (2021); Sun et al. (2022) and low-dimensional structure that is robust across behaviors, time, and animals Gallego et al. (2020); Safaie et al. (2023). Other work has characterized the geometry of abstraction in cortex and hippocampus Bernardi et al. (2020); Courellis et al. (2024). Also the capacity for few-shot learning Sorscher et al. (2022) and semantic cognition Saxe et al. (2019) are each reflected in neural population geometry.

A wide range of methods have been developed to characterize neural population geometry and identify collective population modes in neural data, including for example, principal component analysis, factor analysis Cunningham and Yu (2014), representational similarity analysis Diedrichsen and Kriegeskorte (2017), canonical correlation between brain areas Semedo et al. (2020); Ebrahimi et al. (2022), and various manifold or topological approaches Giusti et al. (2015); Zhang et al. (2023). Despite their diverse formulations, these methods share a common foundation: they all depend only on second-order statistics, describing how pairs of neurons covary across trials, conditions, or time. In fact, many advanced methods for non-linear manifold inference and topological analysis take as input only a matrix of second-order relations between pairs of neurons, be it correlations, or the closely related covariances or distances. The neural correlation matrix thus serves as a foundational object underlying geometric analyses of population activity.

The fundamental importance of neural population geometry and its basis in pairwise neural correlations, raises a foundational question in neuroscience: when, why, and how can we at all reliably estimate such neural population geometry? For example, while we can now record ten thousand single neurons, this number still undersamples the total number of neurons in many brain circuits controlling complex behaviors. For instance, a single hemisphere of macaque primary motor cortex likely contains 1.5-3 million neurons that represent the motion of a single forelimb Young et al. (2013); Higo et al. (2016). Yet many studies of the neural population geometry of motor cortex record only a few hundred neurons. What theoretical principles govern our ability to accurately estimate (if at all) single-trial neural population geometry with 4 orders of magnitude undersampling? How would the geometry change (if at all) were we to record more neurons.

Even more stringently, data is often quite trial limited. In complex behavioral experiments with many trial types, we may only be able to obtain simultaneous neural recordings with tens to hundreds of trials of any given trial type, due to animals’ inability to maintain task engagement within a single session, and due to our potential inability to reliably track neurons across multiple sessions. Thus modern large-scale neuroscience datasets can have many fewer trials than neurons. For example, if we record 100 trials while recording 10,000 neurons, our data is fundamentally a cloud of 100 points in a 10,000 dimensional neural activity space. How can we possibly extract veridical neural population geometry from so few points in such a high-dimensional space? Indeed the number of pairwise neural correlations grows as the square of the number of neurons. So with 10,000 neurons we have about 50 million neural correlations to estimate. How can we reliably estimate so many correlations with only 100 trials? How would errors in our estimate of these correlations propagate to the estimation of neural population geometry, and then to the scientific inferences that many studies make based on this geometry? These issues will only assume greater severity in the future, as recording technologies become better, leading to more simultaneously recorded neurons, but the difficulty of maintaining animals’ task engagement within a single session remains unchanged, leaving us with the same limited number of trials. For example, if we are able to record one to ten million neurons, will we need to dramatically increase the number of trials in order to accurately estimate neural population geometry? This work addresses these questions by offering a predictive theory of experimental design that can tell us how many neurons and trials we will need to record in order to obtain neural population geometry estimates with a given reliability, before we even do the large-scale experiments, by extrapolating from smaller scale experiments. We focus on the simplest but most widely used method for extracting such geometry: principal component analysis (PCA).

We first focus on the starting point of PCA, the matrix of neural correlations. We show that the reliability of our estimate of the entire matrix of neural correlations depends on the ratio of a very specific measure of data dimensionality to the number of trials. We show how to extrapolate the data dimensionality of larger recordings from measurements of data dimensionality of smaller recordings, which enables us to predict the reliability of neural correlations from small pilot recordings. This yields a predictive theory of experimental design for neural correlations that can predict the number of trials and neurons needed in order to achieve a desired level of reliability *before* we do larger scale recordings (assuming the new neurons share patterns of covariation with the old neurons, as explained below). Our results show that data dimensionality grows systematically with the number of recorded neurons, which yields a *curse of dimensionality* for the entire neural correlation matrix: as we record more neurons, we *must* record more trials to preserve its reliability. This curse may be mild if the dimensionality of the entire population is low to begin with, but as has been reported (and as we confirm below) this is not always the case in contemporary experiments Stringer et al. (2019); Manley et al. (2024).

Next we move on to individual neural modes and their mode strengths (i.e. eigenvectors and eigenvalues of the neural correlation matrix). The decomposition of single-trial activity patterns into the strongest modes defines a lower dimensional neural population geometry. We show that in contrast to the *entire* neural correlation matrix, individual modes do *not* suffer from a curse of dimensionality. Instead they enjoy a blessing of dimensionality Donoho (2000) whereby recording *more* neurons actually requires *fewer* trials to accurately estimate *individual modes*. Thus remarkably, we can trade off two very seemingly different experimental resources, neurons and trials, against each other while reliably estimating neural population geometry. This is excellent news for the future of systems neuroscience: as recording technologies become more advanced, and we observe more of a neural population, it will become *easier* to estimate low-dimensional neural population geometry with a smaller number of trials, even as it becomes *harder* to obtain all pairwise neural correlations.

An interesting dividend of our theory is the first step towards a theory of scaling laws for foundation models in neuroscience. A number of recent studies have set out to use self-supervised learning in order to train models to acquire transferable representations of neural activity Wang et al. (2025). In particular, several approaches incorporate the framework of masked autoencoders He et al. (2021), masking a number of neurons and using the rest to predict their activity (see e.g. Le and Shlizerman (2022); Zhang et al. (2024); Willeke et al. (2026)). How the prediction error scales with dataset size (in terms of number of neurons and trials) and model size then become central issues in building and exploiting such foundation models. In the very simple case of linear autoencoders and data from a single session, we analytically derive such scaling laws based on our theory of estimating neural population geometry. These scaling laws exhibit power-law behavior, as observed in more complex settings, and we relate the power laws to power laws in the strength of neural modes.

In the following, we lay out in sequence our predictive theory of experimental design for inferring neural population geometry, from dimensionality, to correlations, to individual neural modes, to scaling laws for neural prediction. At every step of the way we test our theory on diverse neural datasets spanning disparate recording modalities (calcium imaging, neurophysiology, and fMRI), brain regions (sensory,motor and whole brain), and species (mouse, monkey and human). Taken together, our results provide firm theoretical guidance for the design of future experiments aimed to reliably extract neural correlations and population geometry in large-scale neuroscience.

## RESULTS

### Overall framework: neural geometry, modes, and dimensionality

Suppose we record the activity of *N* neurons over *T* trials, yielding an *N* × *T* data matrix *R* (Figure 1A, top). Each column is a single-trial pattern of activity across the recorded population, or equivalently a point in *N* -dimensional neural activity space (Figure 1B, bottom). Essentially all large-scale recording analyses begin from such a matrix.

**Figure 1:**
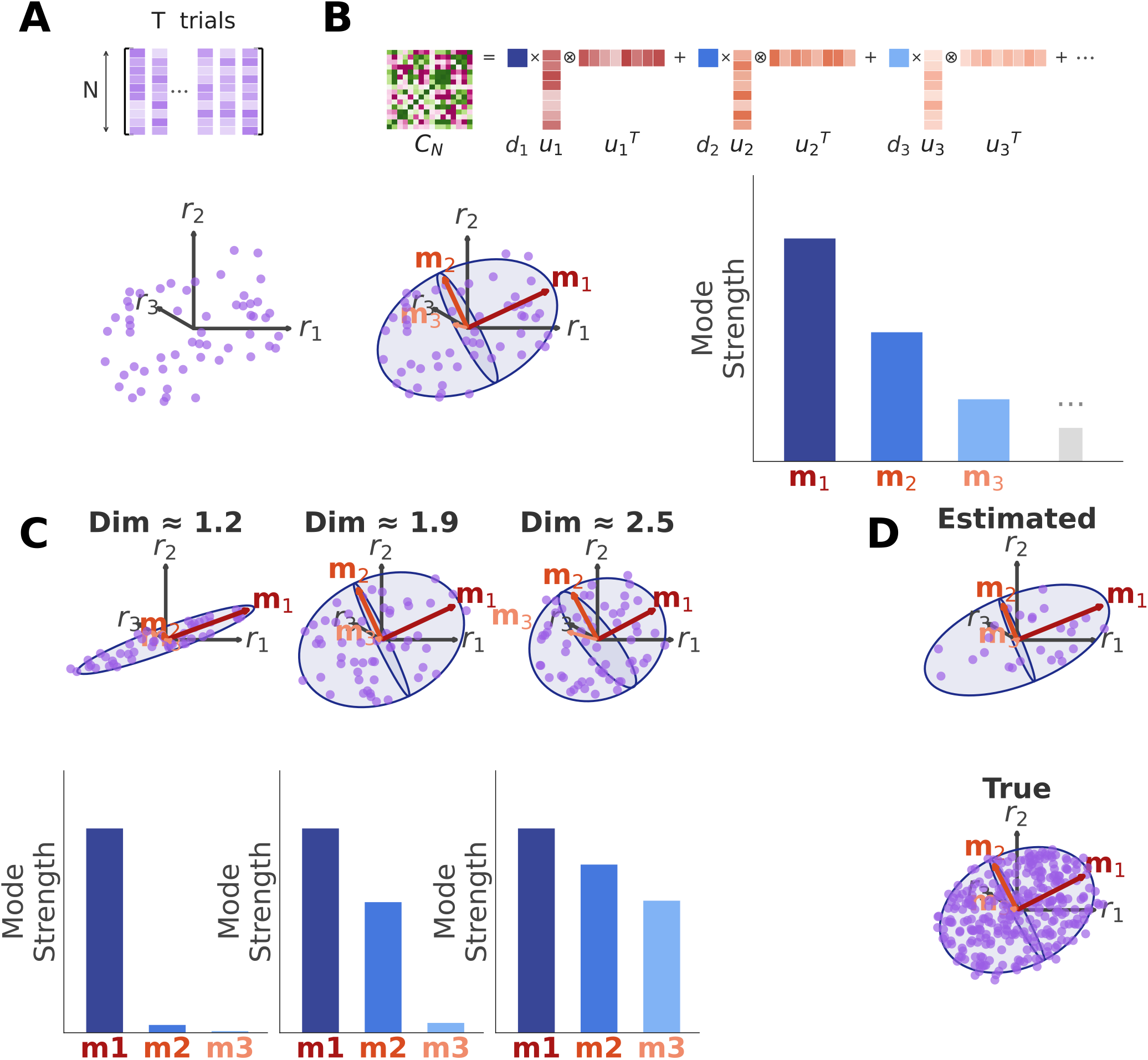
Estimating neural population geometry from finite data through PCA. **A.** A recording of *N* neurons and *T* trials is represented an *N* × *T* matrix (*top*), and as *T* points in *N* -dimensional neural activity space (*bottom*). **B**. *Top:* The *N* by *N* data covariance matrix can be decomposed into *N* data modes (eigenvectors, red) and *N* data mode strengths (eigenvalues, blue). *Bottom:* Geometrically, the data modes are directions in neural activity space (red) and single-trial activity patterns (points) are preferentially spread out along data modes with the largest mode strengths. **C**. Schematic representation of participation-ratio dimensionality of data mode strengths at three different values of dimensionality. **D**. Estimated data modes will generically deviate from the true neural modes at small *T* (*top*), converging to them only as *T* becomes large (*bottom*).

The most common approach to extracting population geometry from *R* is principal component analysis (PCA). PCA starts with the *N* × *N data covariance* matrix, *Ĉ*_*N,T*_, whose (*i, j*) entry quantifies how neurons *i* and *j* covary across the *T* trials (see Methods: Notation). PCA extracts from *Ĉ*_*N,T*_ (Figure 1B, top) a set of *N* estimated *data modes* (eigenvectors of *Ĉ*_*N,T*_). Each data mode can be thought of as a pattern of neural activity across neurons, and has an associated estimated *data mode strength* (the corresponding eigenvalue of *Ĉ*_*N,T*_). This mode strength measures how strongly neuronal activity varies along the corresponding neural mode from trial to trial, through the trial-to-trial variance along the mode. We order the data modes by their data mode strength, from strongest to weakest.

Now each single-trial neural activity pattern can be decomposed into a weighted sum of data modes. If only a small number of modes have appreciable mode strengths, then one can approximate every neural activity pattern in terms of its weights on the strong modes only. For example, if only 2 modes have appreciable strength, each single-trial neural activity pattern can be visualized as a point in a two dimensional space, instead of the original *N* dimensional space (Figure 1B, top). The two coordinates of each trial correspond to the weights of that trial’s neural activity on the first and second data modes. The geometry of this low-dimensional point cloud often reveals insights into the nature of neural dynamics and computation Kaufman et al. (2014); Gallego et al. (2018); Russo et al. (2018); Hénaff et al. (2019); Stringer et al. (2019); Zimnik and Churchland (2021); Sun et al. (2022).

A widespread measure of the complexity of this neural population geometry is its dimensionality. We focus on a specific definition of *data dimensionality* 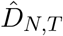, often referred to as the participation ratio of the data mode strengths. The data dimensionality, 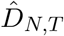, is given by the ratio of the square of the sum to the sum of the squares of data mode strengths (see Methods: Dimensionality). 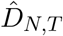 quantifies the effective number of modes over which the total variance of neural activity is dispersed, ranging from 1 (all variance in a single mode) to *N* (variance spread equally across modes). As we will show, this specific definition of dimensionality plays a privileged role in the following theory: it precisely governs the accuracy and reliability of the estimated neural covariance *Ĉ*_*N,T*_ .

With this background, it is clear that accurate estimation of neural population geometry (Figure 1D) relies crucially on accurate estimation of modes and their strengths. However, for large-scale recordings we are often trial limited, with *T* ≪ *N* . This can lead to substantial errors in the estimate of all pairwise neural correlations in the data covariance matrix *Ĉ*_*N,T*_, and therefore errors in the estimated data modes, their strengths, and the extracted neural population geometry. Only in the limit of a very large number of trials (*T* → ∞) does the data covariance *Ĉ*_*N,T*_ converge to the true neural covariance matrix, which we denote by *C*_*N*_ . The true neural covariance matrix will have its own true *neural modes*, true *neural mode strengths*, and true *neural dimensionality*, which we denote by *D*_*N*_ . The true neural modes, their neural mode strengths and the true neural dimensionality will differ from the estimated data modes, their data mode strengths, and the data dimensionality (Fig. 1D top and bottom). We would like to understand the nature of this discrepancy, as finite datasets could lead to a distorted view of the true neural population geometry obtainable only in the infinite data limit.

Furthermore, beyond trial limitations due to finite *T*, we may also be neuron limited, due to a smaller *N* than we would like. Given the precious nature of experimental resources like the number of simultaneously recorded neurons and trials, we would like a *predictive* theory for estimating neural population geometry that can tell us, given the recordings we have made so far, how our ability to accurately estimate neural correlations, neural dimensionality, individual neural modes, their strengths, and the attendant neural population geometry would change if we were to record *more* neurons and/or trials in *future* experiments. Moreover, how do the answers to these questions depend on the complexity and nature of neural activity itself and how should we measure this complexity? In the following, we answer these questions in sequence, first for neural dimensionality, then for the data covariance itself, then for individual neural modes and their strengths. We then show how this theory provides guidance for building simple linear autoencoders of neural activity, and explains the origin of scaling laws in their size and performance. This latter result provides a fundamental first step in analyzing more complex autoencoders in more diverse data, which in turn is a key step in building foundation models in neuroscience.

Throughout, we test the predictions of our theory against the following four datasets: (1) *Mouse V1 multiplane Ca*^*2+*^: 28 recording sessions from mouse primary visual cortex with volumetric, multiplane Ca^2+^-imaging during presentation of natural images Stringer et al. (2019); (2) *Mouse V1 Large FOV Ca*^*2+*^: 11 sessions from mouse primary visual cortex employing Ca^2+^-imaging with a 16-beam, large field-of-view microscope during passive presentation of one of two pairs of oriented visual gratings Rumyantsev et al. (2020); (3) *Monkey dense electrode*: 24 recording sessions from monkey primary motor and dorsal premotor cortices with high-density Neuropixels (phase 3A) probes during performance of center-out reach tasks Sun et al. (2022); O’Shea et al. (2022); Vyas et al. (2020); (4) *Human fMRI*: 8 sessions from humans across visual areas with ultra-high-field fMRI during presentation of natural scenes Allen et al. (2022) (for more information on the datasets and preprocessing see Methods: Datasets). This diversity allows us to validate the predictions of our theory over orders of magnitude variation in neurons and trials, and across recording modalities, brain regions, species, and functionality from sensory to motor.

### A predictive theory of how dimensionality grows with neurons and trials

Suppose one has measured *N* neurons across *T* trials and obtained a data dimensionality 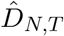. A fundamental question then arises: how would this dimensionality change if we recorded *more* neurons and trials? In particular, if we measured the *entire* neural population for infinitely many trials, what true dimensionality *D*_∞_ would we obtain, and how different would 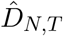 be from it?

To answer these questions we need only two assumptions. First, we assume successive trials are statistically independent and identically distributed such that as *T* → ∞ the data covariance *Ĉ*_*N,T*_ converges to the true neural covariance *C*_*N*_, and the data dimensionality 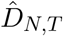 converges to the true neural dimensionality *D*_*N*_ . Second, to predict dimensionality across different numbers of neurons, we assume the neural modes of *C*_*N*_ are distributed indistinguishably across recorded and yet-to-be-recorded neurons (See Supplemental Section S5.2 for the precise mathematical statement of this assumption and Fig S1 for experimental validation).

Under these assumptions, as long as the data dimensionality 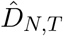 is strictly smaller than both *N* and *T*, we find a very simple and universal formula for the measured data dimensionality (see S5 for derivation):

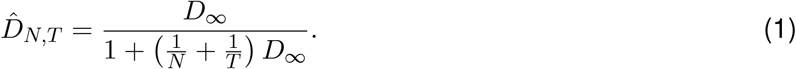

Remarkably, this formula depends on the true neural covariance *C*_*N*_ *only* through a *single* number *D*_∞_, and it reveals that 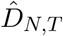 monotonically grows with neurons *N* and trials *T* until it approaches the true dimensionality of the entire population *D*_∞_ (Figure 2A). Notably, two very different experimental resources, neurons *N* and trials *T*, enter completely symmetrically through the sum of their reciprocals. Also, *D*_∞_ sets the scale for both how many neurons and trials are required to accurately estimate dimensionality: if *either N* or *T* are less than or similar to *D*_∞_, then the measured data dimensionality 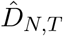 will substantially underestimate the true neural dimensionality *D*_∞_ (e.g. if *N* = *T* = *D*_*∞*_ then 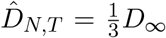). Only when *both N* and *T* are much larger than *D*_∞_ (e.g. by a factor of 10), does 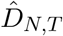 approach *D*_*∞*_ (Figure 2A). Moreover, in actual recordings, where we measure 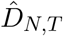, and we know *N* and *T*, we can invert Eq. (1) to obtain *D*_∞_ (Methods Eq.(17)), thereby finding the *true* dimensionality of the *entire* neural population.

**Figure 2:**
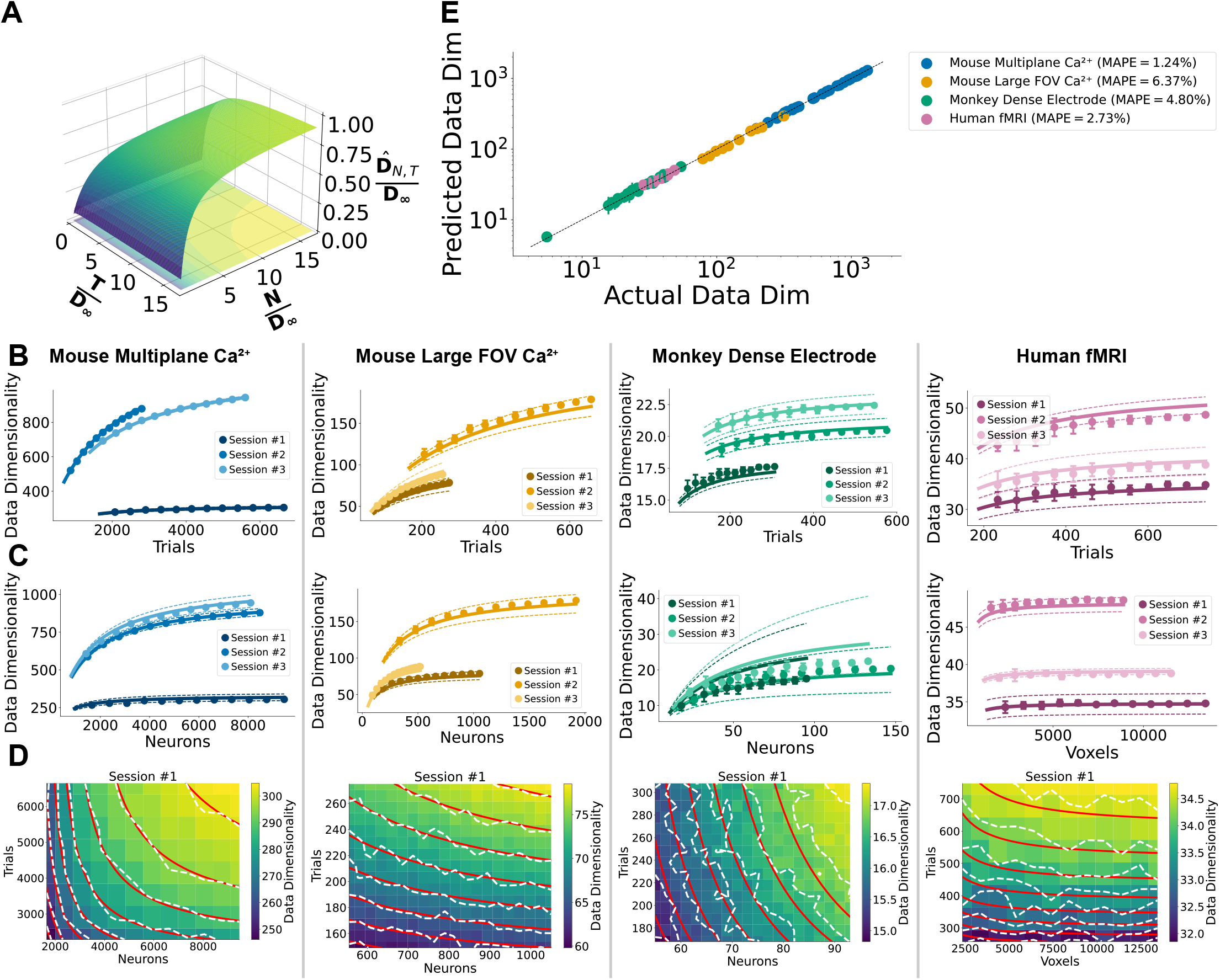
A predictive theory of data dimensionality. **A.** A plot of Eq. (1) showing how the growth of data dimensionality 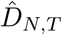 with neurons *N* and trials *T* is governed by a *single* number: the true neural dimensionality *D*_∞_ of the *entire* population. In (B-D) each of the 4 columns displays theory and experiment from one of the 4 datasets, from left to right: Mouse MP, Mouse FOV, Monkey, Human. **B**. Theoretically predicted (lines) and experimentally measured (dots) data dimensionality as a function of trials (Methods Eq. (15)). Predictions are obtained from 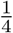 of the trials and all the neurons. Both predictions and measurements are displayed with a mean and standard deviation over repeated subsamples of trials. Numbers of repetitions for each dataset are listed in Methods: Dimensionality. Within each column, each of the 3 shades of color display predictions and measurements from 3 different sessions (or brain regions in the case of fMRI data) of the same dataset. **C**. Theoretically predicted and experimentally measured data dimensionality as a function of neurons (Methods Eq. (16)). Predictions are made from 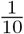 of the neurons and all of the trials. Same format as (B). **D**. Color plot and contours (in dashed white lines) of measured data dimensionality as a function of both trials and neurons, together with theoretically predicted contours in red (Methods Eq. (18)). Predictions are made from subsamples of neurons and trials corresponding to the bottom left point of each grid. **E**. Scatter plot of the theoretically predicted data dimensionality, extrapolated from 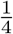 of the neurons and 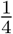 of the trials, versus the measured data dimensionality, 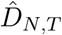, for every single session across all four datasets. Error bars display the standard deviation across subsamples. Mean absolute percentage error (MAPE) in predicted dimensionality is reported for each dataset.

A natural question then arises: how can we experimentally test Eq. (1) if we cannot actually independently experimentally measure *D*_∞_, which would require an inaccessible infinitely large dataset? A key idea is to instead test the predictive capacity of Eq. (1) to extrapolate the dimensionality of *future larger* recordings, from the measured dimensionality of *past smaller* recordings. For example, given 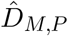 measured from a past small recording of *M < N* neurons and *P < T* trials, we can extract *D*_*∞*_ and then use it to predict 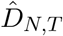 in an arbitrarily larger future recording of *N* neurons and *T* trials (see Methods Eq. (18) for the formula that predicts 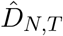 directly from 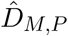).

In this manner, we test this predictive theory of dimensionality across our four datasets, finding close quantitative agreement between theory and experiment. Figure 2B shows successful prediction of dimensionality growth as a function of trials alone (*M* = *N*, *T > P*), and Figure 2C shows the same for neurons alone (*P* = *T*, *N > M*). Figure 2D shows joint prediction across both neurons and trials simultaneously. Finally, Figure 2E summarizes prediction performance across all sessions from all four datasets: we predict the dimensionality of each entire dataset, by extrapolating the dimensionality obtained from just 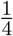 of the neurons and 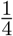 of the trials. The predicted dimensionality matches the measured value with mean absolute percentage error (MAPE) under 7%.

These results establish a predictive theory of data dimensionality that is widely applicable across species, brain regions, recording modalities and brain functions. Remarkably, a single number *D*_∞_, which reflects the true neural dimensionality of the *entire* population, completely governs how measured dimensionality grows with neurons and trials (Eq. (1)), in a manner that allows this dimensionality to be predicted in future experiments that have not even been performed yet.

### A predictive theory for the accuracy and reliability of neural correlations

Next, we ask how well the data covariance *Ĉ*_*N,T*_ estimates the true neural covariance *C*_*N*_, and how reliably it replicates across repeated experiments. Interestingly, we find that accuracy and reliability are each governed by the neural and data dimensionality, respectively.

We quantify estimation accuracy by the fractional squared error between *Ĉ*_*N,T*_ and *C*_*N*_, EstErr(*Ĉ*_*N,T*_) (see Methods: Accuracy), with 0 corresponding to perfect estimation. We derive an exceedingly simple formula for the error (see Supplemental S6.1 for a derivation):

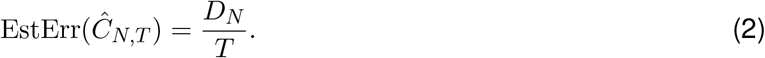

Remarkably, the estimation error depends on the neural covariance matrix *C*_*N*_ *only* through its neural dimensionality *D*_*N*_ . Moreover, our expression reveals that if *D*_*N*_ is much smaller than *N*, we can estimate the entire *N* × *N* matrix of pairwise correlations — *O*(*N* ^2^) numbers — with *T ≪ N* trials.

In real recordings we never have access to *C*_*N*_, so the accuracy cannot be measured directly, and therefore Eq. (2) cannot be tested in any finite neural recording. We therefore turn to the closely related question of *reliability* : if two recording experiments are performed on the same population of *N* neurons, each yielding *T* disjoint trials, how similar are the two resulting data covariances? We define the *replication error*, RepErr(*Ĉ*_*N,T*_), as the fractional squared error of one data covariance relative to the other (Methods: Reliability), with 0 error corresponding to complete reliability. We derive (Supplemental S6.2):

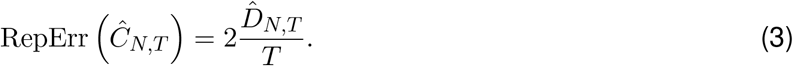

Remarkably, the replication error is simply twice the ratio of data dimensionality to trials. Together, Eqs. (2)–(3) establish a unified picture: accuracy and reliability are each governed by the ratio of a specific dimensionality measure to the number of trials. This highlights a privileged role for the participation ratio measure of dimensionality as opposed to any other definition of dimensionality, as far as the accuracy and reliability of the entire covariance matrix is concerned.

In contrast to accuracy, the replication error *can* be measured directly in neural data, and therefore we can directly test the validity of Eq. (3). We can even *predict* the replication error in *future larger* scale recordings from measurements of the data dimensionality of *past smaller* scale recordings. For example, given 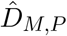 measured from a past small recording of *M < N* neurons and *P < T* trials, we can predict 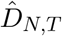 in an arbitrarily larger future recording of *N* neurons and *T* trials (as in Figure 2), and then insert this into Eq. (3) to predict the replication error. This predictive capacity also provides a basis for experimental design by answering the question of how many more trials *T > P* you will need, if you plan to record more *N > M* neurons, to achieve a given desired reliability for all pairwise neural correlations.

We test this predictive theory of reliability across our four datasets (Figure 3). Figure 3A predicts the increase in reliability (decrease in replication error) with additional trials at fixed neurons *N*, starting from 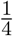 of the recorded trials. Figure 3B predicts the *decrease* in reliability with additional neurons at fixed *T*, starting from 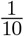 of the recorded neurons. Figure 3C shows joint prediction across both neurons and trials for an example session from each dataset. Figure 3D summarizes performance across all sessions, and demonstrates successful prediction of the achieved reliability on the entire dataset, using only the data dimensionality extracted from 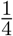 of the neurons and 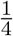 of the trials.

**Figure 3:**
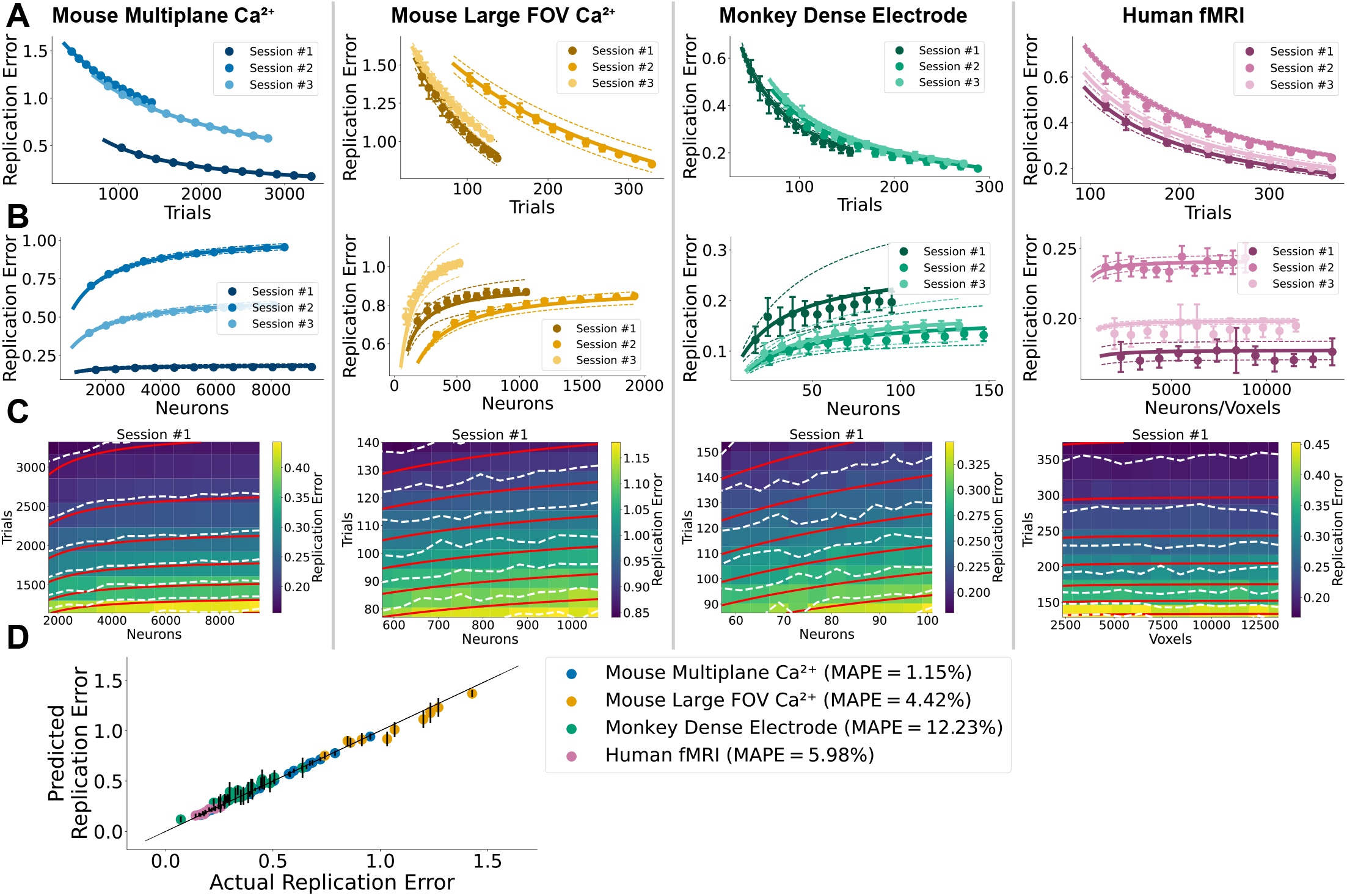
A predictive theory for the reliability of all neural correlations. In (A-C) each column displays theory and experiment from a distinct dataset using the same format as in Figure 2B-D. **A**. Theoretically predicted and experimentally measured replication error (Eq. (3)) as a function of trials. Predictions are obtained from 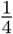 of the trials and all the neurons. Both predictions and measurements are computed from 10 independent subsamples. **B**. Predicted and measured replication error as a function of neurons. Predictions are from 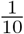 of the neurons and all of the trials. In both (A) and (B) dots and error bars display standard deviation over measurements from subsamples, while lines and dashed lines display mean and standard deviation over predictions. In each column, each of the 3 color shades display predictions and measurements from 3 different sessions of the same dataset. **C**. Color plot and contours (in dashed white) of measured replication error as a function of neurons and trials, together with the theoretically predicted contours in red. Predictions obtained from subsamples of neurons and trials corresponding to bottom left grid point. Both predictions and measurements are averaged over independent subsamples: Mouse MP: 10 repetitions, Mouse FOV: 30 repetitions, Monkey: 50 repetitions, Human: 10 repetitions. **D**. Scatter plot of the predicted replication error, extrapolated from 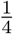 of the neurons and 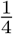 of the trials, vs the measured replication error, for all sessions (or brain regions in fMRI data) across all four datasets. Error bars display the standard deviation of predictions calculated from 10 independent subsamples. The mean absolute percentage error (MAPE) of predicted replication error is displayed for each dataset. Throughout the figure, predictions are obtained by calculating 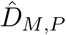, of a subsample (*M < N* and *P < T*), and applying Methods Eq. (23). This is equivalent to extrapolating to 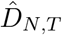 (Methods Eq. (18)), and applying Eq. (3): See Methods: Reliability Procedure) for additional details.

We note Eq. (3) reveals that, at fixed *T*, replication error *increases* with *N*, because 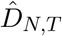 increases with *N* as described in Eq. (1). This is not a small effect: in some of our datasets 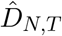 continues to grow with neurons even at the largest recording sizes available, without clear saturation in some of the calcium imaging datasets at ∼ 10,000 neurons (Figure 2C). So in general, recording more neurons *N* without recording more trials *T* unavoidably makes the full pairwise neural correlation matrix *less* accurate and *less* reliable. However, interestingly, in the following we will show that there are specific signals hiding within the neural correlations that become *more* accurate and *more* reliable, even as the *entire* pattern of neural correlations becomes *less* accurate and reliable. These hidden reliable signals correspond to shared patterns of covariation across neurons, which can be uncovered by focusing only on the strongest data modes, as we see next.

### A simple low-dimensional model of shared signal and private noise

In order to uncover hidden, potentially reliable signals within a pattern of otherwise unreliable neural correlations, we define and analyze a simple low-dimensional generative model of neural data consisting of *K* latent variables each with signal strength 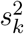 driving *N* recorded neurons through an outgoing signal mode connectivity ***u***_*k*_ (Fig. 4A, top). Additionally, we assume each recorded neuron receives independent private noise of variance *σ*^2^. Importantly, by *signal*, we mean merely any pattern of trial-to-trial variability that is shared across neurons. This could include, for example, sensory inputs, motor commands, cognitive states, or intrinsic network fluctuations. And by *noise*, we mean merely trial-to-trial variability that is private to each neuron and independent across neurons.

**Figure 4:**
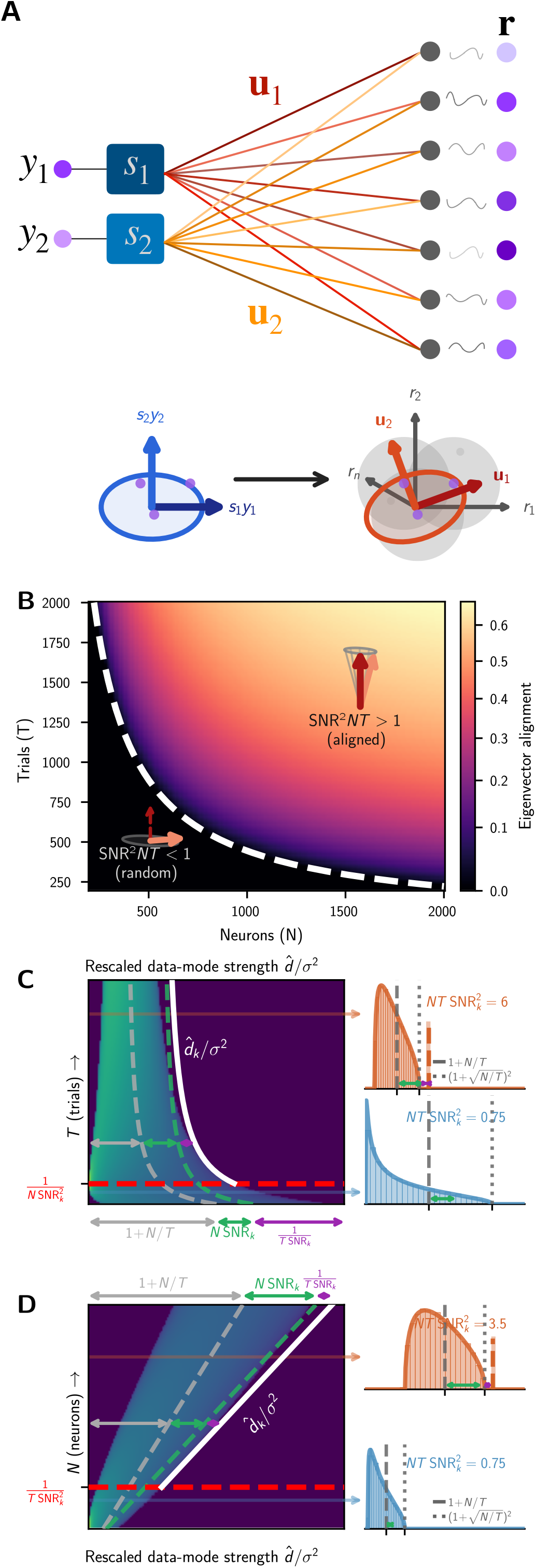
A low-dimensional model of shared signal and private noise. **A.**A schematic representation of neural data generated from 2-dimensional latent activity *y*_1_ and *y*_2_, scaled by the signal mode strengths, *s*_1_ and *s*_2_, and broadcast through the signal mode vectors ***u***_1_ and ***u***_2_ to yield shared clean signal amongst *N* neurons (grey dots). Then private noise is added to each neuron to yield the signal plus noise pattern **r** (purple dots). *Bottom:* Geometrically this corresponds to a mapping of single trials (purple dots) in a 2 dimensional latent space (left) to neural patterns in an *N* dimensional neural activity space with shared signal along the 2 modes (red arrows), corrupted by private noise (grey disks). **B**. A heatmap of the alignment *A*_*k*_ in Eq. (6) between a true signal mode and the data mode estimated from *N* neurons and *T* trials, given a fixed per-neuron SNR for the mode. The detectability phase transition below which the alignment is 0 is shown as the dashed white curve, which obeys *NT* SNR^2^ = 1. **C**. *Left* : A heat map of the Marchenko-Pastur (MP) noise sea of data mode strengths (blue-green horizontal slices) as a function of increasing trials *T*, for a fixed number of neurons *N* . As *T* increases, the noise sea compresses and its right edge or surface falls. When *T* crosses the detectability threshold 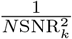 for signal mode *k* (dashed red line), a single outlier data mode strength appears at the location given by Eq. (7) (white line). *Right* : Horizontal slices corresponding to two values of *T*, one below and one above threshold, displaying the theoretical data mode strength distribution, together with a numerical histogram from simulated data. **D**. Analogous to (C) but shows results with increasing neurons *N* at fixed trials *T* . The noise sea expands and its surface rises, but after the detectability threshold at 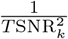 (dashed red line), with increasing *N*, the outlier data mode strength (white line) outpaces the rise of the noise surface, even at fixed trials *T* .

This low-dimensional model yields a neural covariance matrix *C*_*N*_ consisting of *K* signal modes ***u***_1_,. .., ***u***_*K*_ each with neural mode strength 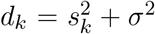, coming from the sum of signal and noise, and *N K* noise modes ***u***_*K*+1_,. .., ***u***_*N*_ each with mode strength *σ*^2^ coming from noise alone (see App. S8 for a mathematical description and analysis of this model). In terms of neural population geometry, the *K* dimensional latent state drives a *K* dimensional signal subspace of shared trial-to-trial neural variability across *N* neurons, spanning the *K* signal mode directions, while additional private trial-to-trial noise potentially obscures the shared signal modes (Fig. 4A, bottom).

While it is reasonable to assume that the private noise strength *σ*^2^ for each neuron is independent of the number of recorded neurons *N*, it is less clear how the signal mode strength 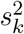 should scale with *N* . To make this crucial modeling choice, we first note that each signal mode ***u***_*k*_ is a unit vector spread across *N* neurons, so its squared components are *O*(1*/N*). Then for the variances and pairwise correlations of *C*_*N*_ (i.e., its individual matrix entries) to be *O*(1) quantities that do not change as we record more neurons, the signal-mode strengths 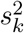 must grow linearly with *N* (see App. S8 for a full mathematical description of this argument). We make this scaling explicit by defining the relation

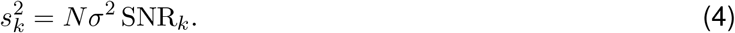

Here SNR_*k*_ can be thought of as a signal-to-noise ratio per-neuron of the signal mode ***u***_*k*_. Indeed SNR_*k*_ is simply the signal-to-noise ratio 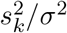 of the mode divided by the number of recorded neurons *N* .

This scaling assumption (Eq. (4)) is a fundamental contribution of our work. It is what allows us to derive a predictive theory of how accurate and reliable estimates of neural population geometry would become if we were to record *more* neurons. The reason why is that this scaling assumption views the per-neuron signal-to-noise ratio as an *intrinsic* property of a neural population (when viewed through a specific recording modality) sharing a common mode of variation, that does *not* change as we record more neurons. Thus, as we will see below, we can infer *SNR*_*k*_ using a small number of neurons and trials, and then predict how accurately we can estimate neural modes and their strengths with more neurons and trials. Also, an immediate consequence of the constancy of the per-neuron *SNR*_*k*_ is that the mode’s overall signal strength 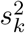 *increases* linearly with the number of neurons. As we will see below, this increased signal in any particular mode, as one observes more of it by recording more neurons alone, has a dramatic and beneficial impact on our ability to better estimate *individual* neural modes of *C*_*N*_, even if the estimate of the *entire C*_*N*_ is degraded as in Eqs. (2) and (3).

### A theory of how shared signal modes emerge from finite and noisy data

In summary, our simple low-dimensional model of neural activity of the previous section posits a true neural covariance matrix *C*_*N*_ with *K* signal modes ***u***_*k*_, for *k* = 1,. .., *K*, each with neural mode strength *d*_*k*_ = *σ*^2^(*N* SNR_*k*_ + 1), and with all remaining *N* − *K* noise modes each having neural mode strength *σ*^2^. However, in any experiment with a finite number of trials *T*, we do not have access to *C*_*N*_ ; we only have access to the finite data covariance *Ĉ*_*N,T*_, which has its own *N* data modes ***û***_*k*_, for *k* = 1,. .., *N* and corresponding data mode strengths 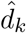. This raises a fundamental question: what is the relationship between individual data modes ***û***_*k*_ and strengths 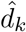, and the true neural modes ***u***_***k***_ and strengths *d*_*k*_? In particular, as a joint function of neurons *N*, trials *T*, and SNR_*k*_, when and how well can a neural mode be estimated? To answer this question, we exploit random matrix theory Benaych-Georges and Nadakuditi (2012), but with our non-standard scaling in Eq. (4) that is a central aspect of this work (see Method Sections: Low-Rank and Supplemental Section S8 for a full derivation of all the following statements).

We find that as one increases the number of recorded neurons *N* and trials *T*, for each signal mode *k*, one passes a detectability phase transition boundary given by

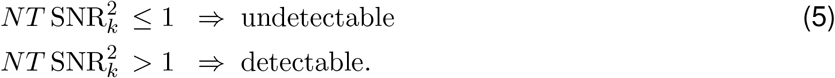

In the undetectable phase, there is no trace of the signal mode in the data, whereas in the detectable phase there is. This detectability phase transition manifests itself most simply in the alignment, or squared cosine angle, between the data mode ***û***_*k*_ and signal mode ***u***_*k*_ given by *A*_*k*_ = | ***û***_*k*_ · ***u***_*k*_ | ^2^ for *k* = 1,. .., *K*. We find this alignment obeys (see Fig.4B for a plot)

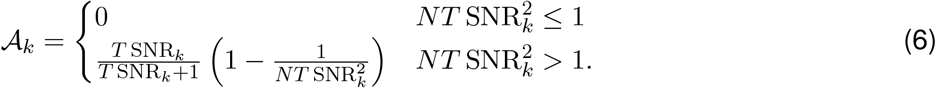

Thus in the undetectable phase, there is no alignment between the data mode and the neural signal mode. However, as one moves deeper into the detectable phase, the alignment increases.

Intriguingly, the detectability phase transition boundary 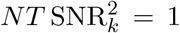 depends on neurons *N* and trials *T only* through their product *NT* . This means there is a hyperbolic tradeoff (e.g. hyperbolic phase transition boundary in Fig.4B) between two very *different* resources, neurons and trials. Recording *more* neurons requires *fewer* trials, and recording more trials requires fewer neurons, to cross the detectability threshold for any given signal mode. Moreover, increased mode SNR_*k*_ lowers the detectability phase transition boundary in the *N* by *T* plane. However, once in the detectability phase, neurons *N* and trials *T* start to play more asymmetric roles. As *T* → ∞ for fixed *N*, *A*_*k*_ → 1 and the data mode aligns perfectly with the signal mode regardless of how few neurons are recorded. However, as *N* → ∞ at fixed *T*, the alignment saturates at *T* SNR_*k*_*/*(*T* SNR_*k*_ + 1) *<* 1. Adding neurons is helpful but cannot fully compensate for limited trials. Still, alignment increases monotonically with *both N* and *T* in the detection phase.

This detectability phase transition boundary also manifests itself in the distribution of data mode strengths 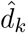. If *NT* is so small that every single one of the *K* modes is in the undetectable phase in Eq. (5), then the distribution of 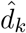 behaves as if the neural covariance *C*_*N*_ were simply pure noise, e.g. *C*_*N*_ = *σ*^2^*I*, with no detectable signal modes. Such a neural covariance would have all mode strengths *d*_*k*_ = *σ*^2^. However, due to finite trials *T*, the corresponding data mode strengths 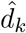 instead spread out to form a Marchenko-Pastur (MP) noise sea Marčenko and Pastur (1967), that ranges from a left edge, or depth of 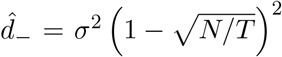 to a right edge, or surface of 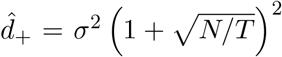 with a midpoint at 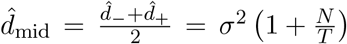 (Fig.4CD). This MP noise sea behaves very differently as a function of neurons *N* and trials *T* . For fixed *N*, as *T* increases, the noise sea compresses its range and increasingly moves its midpoint towards the true noise level *σ*^2^ from above (Fig.4C). In contrast, for fixed *T* as *N* increases, the noise sea expands its range and both its midpoint and its surface rise away from the true noise *σ*^2^ (Fig.4D).

Now as the product *NT* increases to cross the detectability threshold in Eq. (5), the noise sea persists as described above, but a single outlier data mode strength rises above the surface of the noise sea at the location

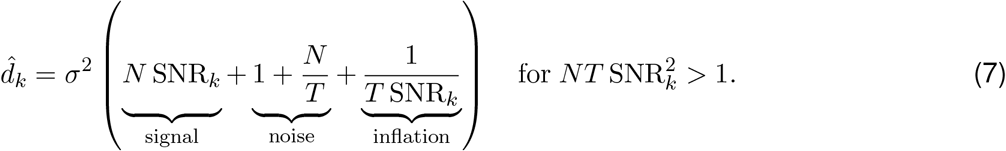

At the same time, the associated data mode ***û***_*k*_ acquires the positive alignment *A*_*k*_ as described above in Eq. (6). The data mode strength decomposes into a sum of three contributions: the signal-mode strength itself, 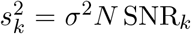, the midpoint of the noise sea, 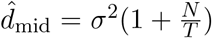, and an SNR-dependent inflation term, 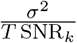, which is diminished with more trials. This expression passes two consistency checks. First, as *T* increases at any fixed *N*, the data mode strength 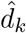 in Eq. (7) decreases to approach true neural mode strength 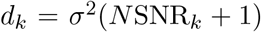. Second, at the transition boundary 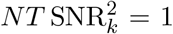, the data mode strength 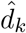 in Eq. (7) rests just at the surface of the noise sea at 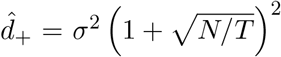 (e.g. substitute 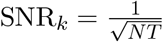 into Eq. (7)).

The outlier data mode strength rises above the surface of the noise sea in two very different ways, when one increases trials *T* at fixed neurons *N* (Fig.4C), versus when one increases neurons *N* at fixed trials *T* (Fig.4D). As one moves across the detectability transition in the former case, increased trials compresses the noise sea, the surface falls, leaving an outlier data mode strength, and then both the sea surface and outlier fall to their true values (Fig.4C). In the latter case, increased neurons expands the noise sea, and its surface *rises*. This may seem detrimental, and it *is* for the purposes of estimating the *entire* pattern of neural correlations as seen in Eqs. (2) and (3). However, for a *single* signal mode, the signal strength 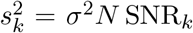 *also* increases, which allows the outlier data mode strength in Eq. (7) to eventually outrace the rising sea and pop out of it as *N* crosses the detectability threshold. Thereafter, *both* the surface of the noise sea *and* the outlier data mode strength rise, though the sea surface cannot catch the outlier (Fig.4D).

In summary, our results show that as the *product NT* of neurons and trials increases, one will see a series of detectability transitions in which each successive signal mode of progressively smaller per-neuron SNR_*k*_ is detected. Detection corresponds to an outlier data mode strength 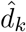 rising out of the surface of the noise sea, and the corresponding data mode ***û***_*k*_ aligning to the neural signal mode ***u***_*k*_. We next test our theory in detail in the next three subsections. First, we show how to fit our theory to data to estimate its critical parameters *σ*^2^ and SNR_*k*_. Second, we test Eq. (7) and our central assumption of the constancy of the per-neuron SNR_*k*_ in Eq. (4), by showing that we can *predict* individual data mode strengths in *future larger* recordings with more trials and neurons, using only estimates of *σ*^2^ and SNR_*k*_ in *past smaller* recordings. Third, we test the theory of the alignment of data modes and signal modes in Eq. (6).

### Fitting a low-dimensional model to data through the shape of the noise sea

To fit our theory to data, we first form an estimate 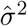 of the noise variance by matching the median data mode strength to the median of the MP distribution (Methods: Model-fit, Figure 5A), following Gavish and Donoho (2014). Then we estimate the surface of the noise sea as 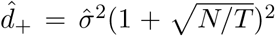. Each data mode with strength 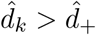 then becomes an inferred signal mode, and we form an estimate 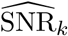 of its per-neuron signal-to-noise ratio by inverting Eq. (7) (see Methods: Model-fit, Eq. (33)). In the following sections we will use this estimated SNR to test our predictions about neural population geometry.

**Figure 5:**
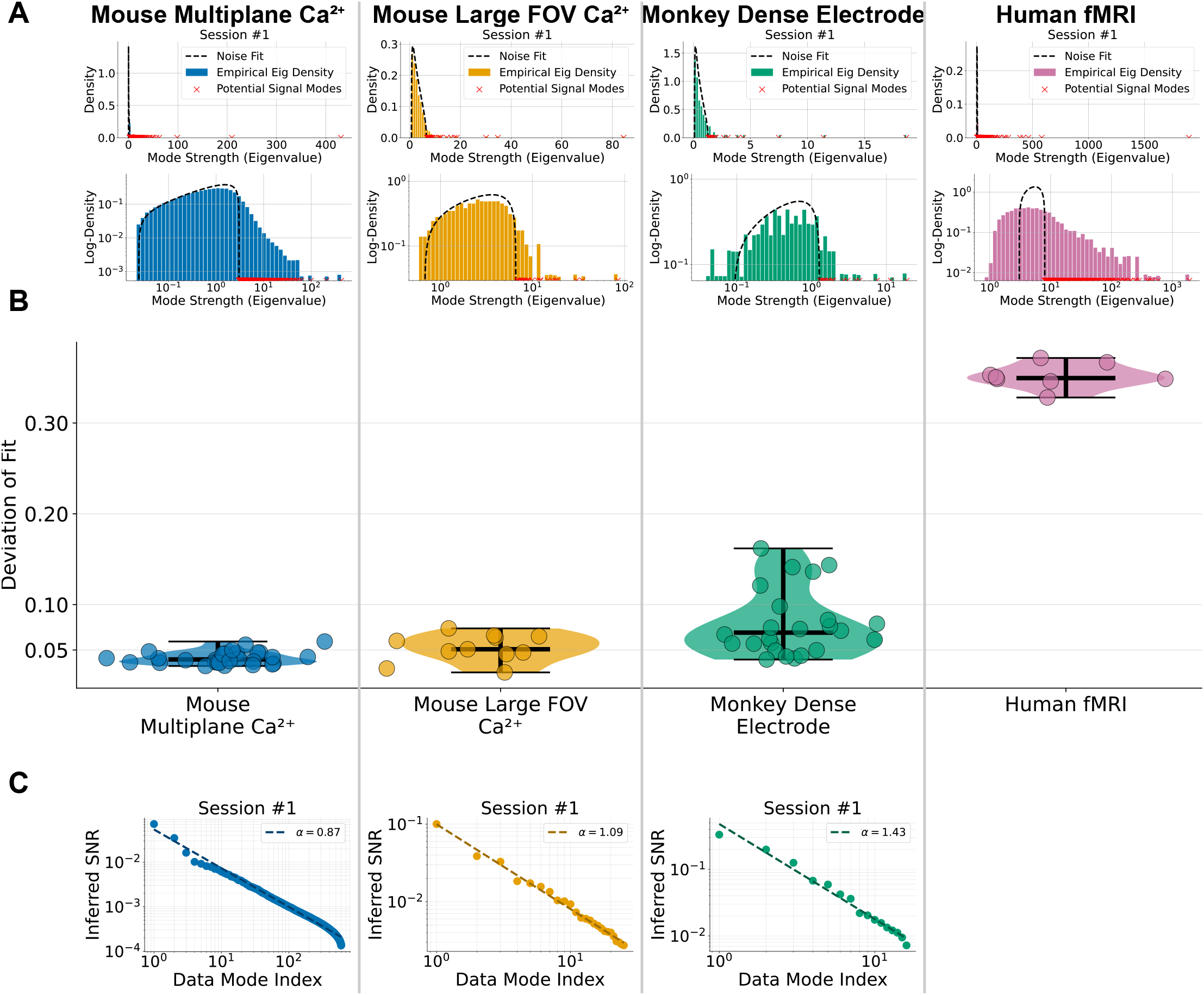
Assessing the goodness of fit of the low-dimensional model. **A.** *Top:* Measured histograms of data mode strengths (colored bars) from 1 session from each of the 4 datasets, together with theoretical fits of the MP noise sea (dashed black curves). The estimated private noise variance 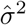 is set to match the median measured data mode strength. Then mode strengths above the estimated sea surface 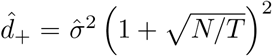, shown by red x’s, correspond to potential signal modes (Methods: Model-fit). *Bottom:* Same theory and measurements as above but shown on log-log axes. **B**. The deviation of the empirically measured data mode strength histogram from its MP noise sea theoretical fit, as quantified by the Wasserstein-1 distance, normalized by the extent of the MP noise sea (Methods Eq. (30)). Each circle denotes an individual session from each dataset. Though measured histograms of data mode strengths do not exhibit a clear separation between the noise sea and outlier mode strengths, the theoretical model nevertheless fits the measured noise sea well for Mouse MP, Mouse FOV, and Monkey, but not for *Human fMRI*. **C**. For each of the potential signal modes (red x’s in (A)), we can estimate their per-neuron SNRs. These rank-ordered inferred SNRs exhibit clear power-law behavior, as shown in Log-log plots of the inferred SNR_*k*_ as a function of index *k* (dots). A linear fit of log SNR_*k*_ vs log *k* (dashed black line) yields an estimate of the power-law exponent, *ε*. See Figure S6 for more recording sessions.

But before testing those predictions, we first assess the goodness of fit of our low-dimensional model by comparing the shape of the empirically observed noise sea to that of the theoretically predicted MP noise sea. In particular, we measure the normalized 1-Wasserstein distance between the two distributions 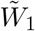 (Methods: Model-fit, Figure 5B). We find the theoretical MP distribution matches well the empirical noise sea in three of four datasets — Mouse MP, Mouse FOV, and Monkey all show 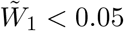. Our low-dimensional model of shared signal and private noise is thus consistent with neural recordings of widely varying complexity, with data dimensionalities 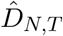 ranging from ∼ 15 to over 1000.

On the other hand, the model does not fit the Human data well 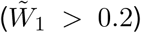. The deviations are consistent with two non-exclusive explanations: correlated noise across trials, and a sufficiently long tail of neural mode strengths that the assumption of a small number of shared signal modes may not apply. We discuss both possibilities, and what kind of theoretical extensions could fit the fMRI dataset in the discussion.

Given the fundamental importance of the per-neuron signal-to-noise ratios SNR_*k*_ in the theory of estimating neural population geometry, we examined the inferred SNR’s across the datasets. Interestingly, for the 3 datasets in which the low-dimensional model fit well, we found that the rank ordered inferred SNR’s exhibited power-law distributions (Figure 5C), and are well fit by a power law,

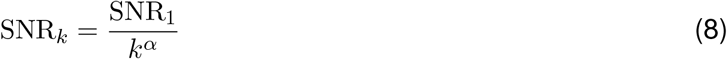

with exponents *ε* varying between 0.6 and 1.5 across sessions and datasets (see Figure S6 for more recording sessions).

We will return to the consequences of this power law for learning autoencoder models of neural activity below. But first, in the next two sections, we test the predictions of our theory in the three datasets where our low-dimensional model fits well.

### Testing the predictive theory of data mode strengths

As described above, a fundamental contribution of our work is the proposition that the per-neuron SNR_*k*_ of a neural mode, defined in Eq. (4), is an intrinsic property of a neural assembly (viewed through a recording modality) sharing common signals, independent of how many neurons or trials are recorded. Here we test this proposition by inferring signal modes and forming estimates 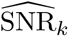 of their per-neuron signal-to-noise ratio using *one* value of neurons and trials, and then *predicting* the data mode strength 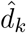 at *many other values* of neurons and trials. We do this simply by inserting 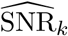 into our prediction for 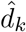 in Eq. (7), using *other* values of *N* and *T different from* those used to form the estimate 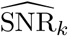.

In particular, in Fig. 6A we estimate 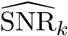 using all neurons and trials, and then compare our theoretically predicted data mode strengths 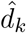 in Eq. (7), for fewer neurons and the same number of trials, to actual measured data mode strengths, finding excellent agreement. Indeed, we can accurately predict more than 100 data mode strengths in the largest datasets. These curves show that our theory can quantitatively predict how data mode strengths become smaller as one records fewer neurons. We next show that we can accurately predict data mode strengths if we were to record *more* neurons and trials. In Fig. 6B we estimate 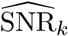 using only 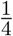 of the neurons and 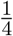 of the trials, and we then successfully predict the data mode strengths 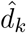 obtained using all the neurons and all the trials.

**Figure 6:**
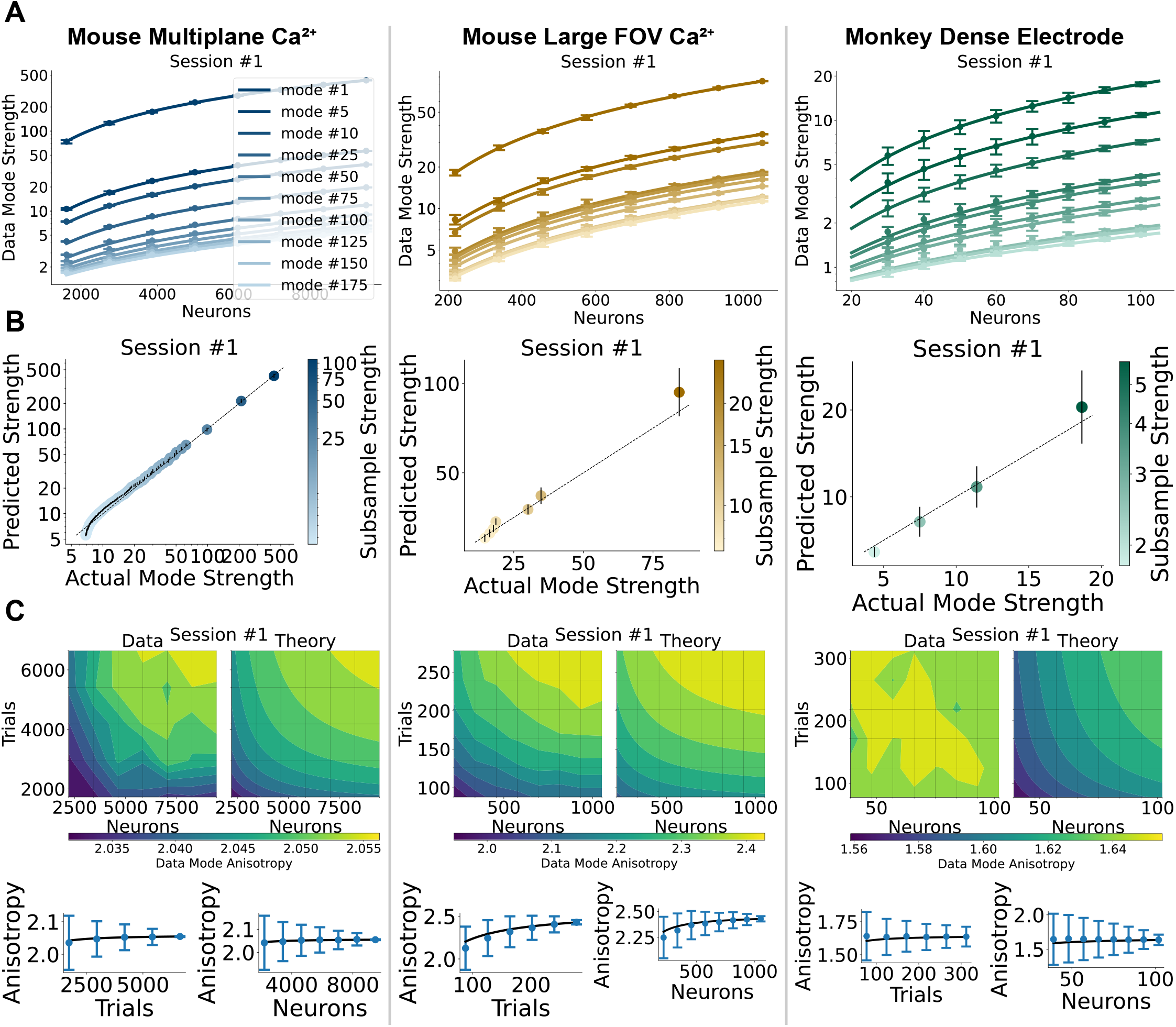
A predictive theory of data mode strengths. The three columns display results from the three datasets that are well fit by the low-dimensional model. **A**. Theoretical predictions (lines) and direct experimental measurements (dots) of data mode strengths as a function of number of neurons. Predictions are derived from estimating the per-mode SNR using all neurons and trials, then using Eq. (7) to predict the data mode strength at fewer neurons. Error bars on measurements reflect standard deviation over repeated subsamples of neurons (Mouse MP: 5 repetitions, Mouse FOV: 10 repetitions, Monkey: 50 repetitions). The top 10 modes are shown unless otherwise indicated. **B**. Predicting data mode strengths in larger recordings: scatter plot of theoretically predicted data mode strengths using per-neuron mode SNRs inferred from 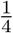 of the neurons and 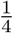 of the trials vs experimentally measured data mode strengths in the full data set using all neurons and trials. Each point is a detected mode in the subsampled dataset and its color denotes its subsampled data mode strength. The mean and standard deviation of theoretical predictions are obtained from repetitions over 10 subsamples. Black line indicates the unity line. **C**. *Top:* Contour plot of aspect ratio between top two data mode strengths, 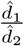, data and theory predicted from full data. *Bottom:* one-dimensional slices showing aspect ratio vs trials and neurons respectively, with solid black line showing theory prediction.

According to Eq. (7) signal modes undergo an SNR-dependent inflation: lower-SNR modes are inflated more than higher-SNR ones — the geometry of the observed signal-mode subspace is distorted. The anisotropy, i.e. the ratio between data mode strengths, eg 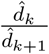, is decreased systematically relative to the anisotropy of the true signal strengths — the observed signal subspace appears *less elongated* than the true signal subspace, with variance distributed more equally across modes than it actually is (Methods: Signal Geometry, Eqs. (34) and (35)). This is the opposite of what we found for the *full* set of data modes, whose data dimensionality is *lower* than the true neural dimensionality — i.e., as a whole, the data modes appear more elongated, not less. The anisotropy predictions as a function of neurons and trials (Figure 6C) match the data well within the boundaries of error.

On the one hand, finite sampling stretches the full set of data modes, inflating high-variance signal modes but also generating under-sampled, artificially low-variance noise modes. On the other hand, when restricted to the signal subspace, finite sampling drives the signal modes toward isotropy.

### Testing the predictive theory of data mode alignment and reliability

We next seek to test our theory of alignment *A*_*k*_ between a data mode ***û***_*k*_ and the true signal mode ***u***_*k*_ in Eq. (6). At first sight, this theory seems untestable because we do not have access to the true signal mode ***u***_*k*_ in any dataset with a finite number of trials. This problem is directly analogous to our inability to directly test our theory for the estimation error of the covariance matrix in Eq. (2) because we similarly do not have access to the true covariance matrix *C*_*N*_ . There we passed from a theory of error in Eq. (2) to a theory of reliability in Eq. (3), which was then eminently testable in finite datasets. Here we follow the same strategy for data modes by passing from alignment to reliability.

Consider a dataset of *N* neurons and *T* trials. Then consider a split of this dataset into two disjoint subsets of the same *N* neurons but only *T/*2 trials. We measure two data modes 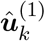 and 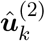, one for each of the two disjoint data subsets. We define the reliability of data mode *k* to be 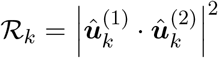. The reliability is simply the squared cosine angle between the two modes computed on the disjoint data subsets, and ranges from 0 (most unreliable and not reproducible) to 1 (completely reliable and reproducible). We derive an exceedingly simple theoretical relationship between reliability and alignment: 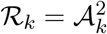 (see Supplemental Section S8.3.3). Since ℛ_*k*_ is completely measurable in data, this relationship makes our theory of mode alignment testable.

In the limit of large *N* and *T*, our theory predicts that the *k*^th^ data mode in the second data subset, if reliable, will align only with the *k*^th^ mode in the first data subset, and no other data mode (Fig. 7A left subplots). However, in finite datasets, overlaps with a given second-subset data mode can “leak” into a range of first-subset data modes with similar data mode strength (Fig. 7A middle subplots), reflecting the fact that individual eigenvectors in a subspace of nearly degenerate eigenvectors are poorly determined — small perturbations rotate the eigenvectors within the subspace — even when the subspace is stable. In practice, this can cause the empirical reliability ℛ_*k*_ to deviate from the theoretical prediction 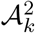 when either *N* or *T* is not sufficiently large, or when signal mode strengths are not well-separated. To mitigate this issue, we consider a better behaved measure, the *subspace* reliability 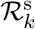, computed by summing the overlap of the *k*^*th*^ mode 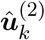 over *all* potential signal modes of the first data subset (see Methods Eq. (39) for precise definition). In effect, 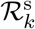 recovers alignment that has been lost to nearby data modes, and indeed we find that 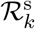 closely matches 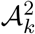 in experimental data (Fig. 7A; left subplots show 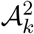, while right subplots show 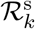 along the diagonal). See Section S8.3.4 for a brief discussion related to subspace overlaps.

**Figure 7:**
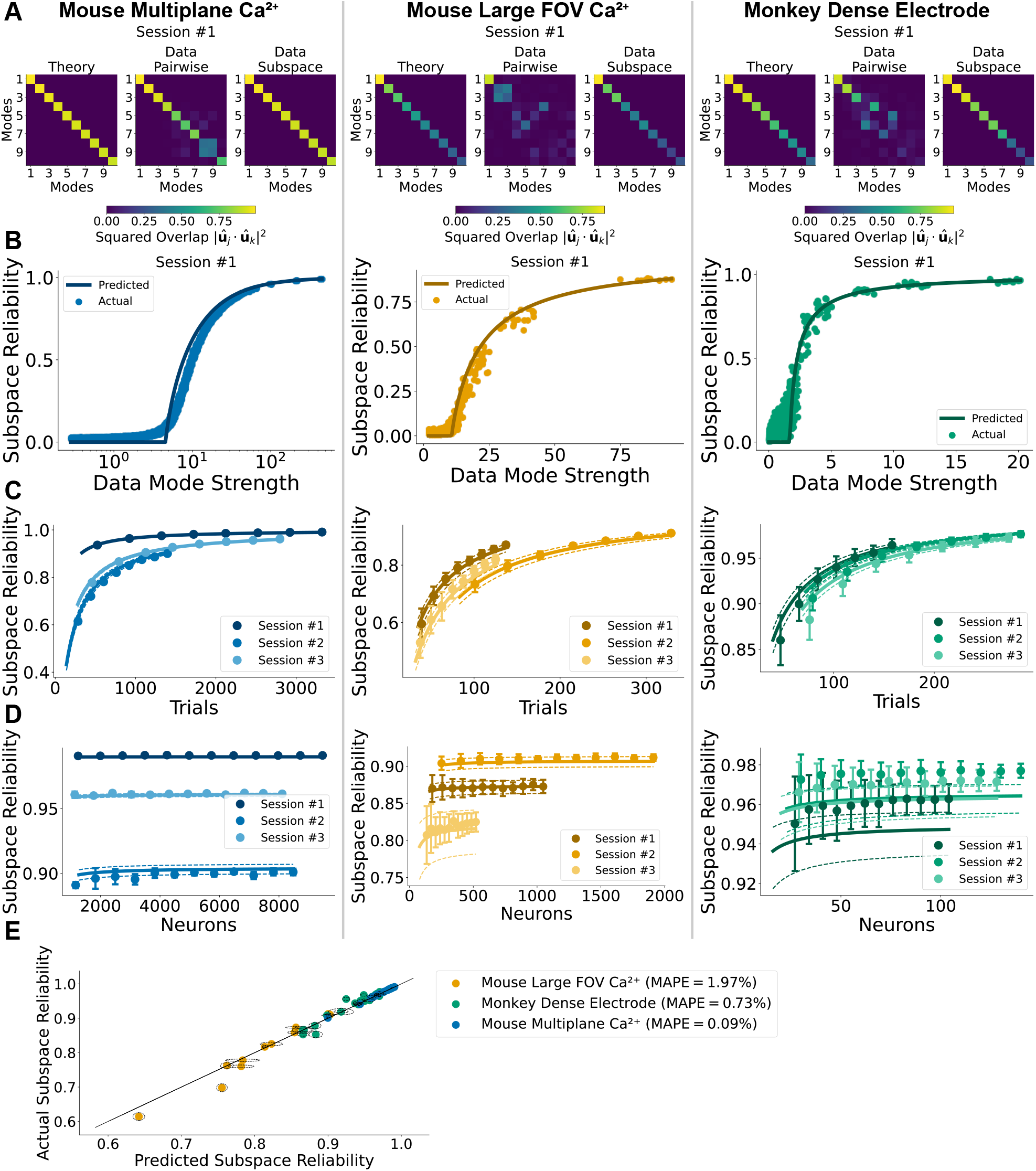
A theory of reliability of data modes via alignment between data modes and signal modes. In panels A-D, the three columns show the three datasets that are well fit by our low-dimensional theory (as in Figure 6). E aggregates across all datasets. **A**. Alignment between data mode *k* from one data subset and data mode *l* from a disjoint subset. *Left:* Theory predicts mode *k* only aligns with itself, with magnitude 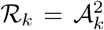 with *A*_*k*_ given by Eq. (6). *Middle:* Measured alignments instead spread across modes. *Right:* Subspace reliability, 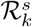, obtained by summing mode *k*’s alignment over the full signal subspace of the other data subset, is well predicted by 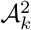(left). **B**. Subspace reliability vs. data mode strength, for a single session; Solid line, theory; dots, every mode from 10 random half-trial-count splits. **C**. Subspace reliability of the strongest data mode vs. trials. Theoretical predictions are derived by inferring the strongest data mode’s SNR from 1*/*4 of trials (1*/*10 for Mouse MP) and then extrapolating to more trials. **D**. Subspace reliability of the strongest data mode vs. neurons. Increasing neurons increases the reliability of individual signal modes. Theory extrapolated from 15 percent of neurons. **E**. Subspace reliability of the strongest data mode (mean over 5 random splits) vs. theory extrapolated from 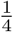 of neurons and 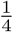 of trials, across all sessions. In (C) and (D), dots/solid lines are mean of experiment/theory, respectively. Error bars and dashed lines (dashed ellipses in E) give measurement SD and prediction deviation, the latter obtained from SNR *±* one SD across subsamples. (C) and (D) use 5, 50, and 100 subsample repetitions for Mouse MP, Mouse FOV, and Monkey, respectively.

Across all three datasets that are well fit by our low-dimensional theory, we find that our theoretical prediction 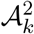 matches well the empirically measured subspace reliability 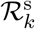 in a variety of scenarios (Fig. 7BCDE). In Fig. 7B for session #1 of each dataset, we split the dataset into two disjoint subsets of *N* neurons and *T/*2 trials and report the measured subspace reliability *R*^s^ for every single mode *k* as a function of the mode strength 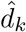 in the second data subset (colored dots). For our theoretical prediction (black lines), we first infer 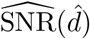 from the first data subset *as a function* of data mode strength 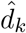 (Methods Eq. (33)). We then insert it into Eq. (6) to obtain the alignment 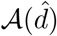 and finally predict the reliability as a function of the subset’s data mode strength as 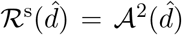. Our theory matches experiments well, and reveals that only modes with mode strengths above the inferred surface of the noise sea are reliable, with the reliability increasing with mode strength height above the noise sea surface.

In Fig. 7CDE, we demonstrate the predictive utility of our theory to design future experiments, by extrapolating reliability in larger recordings from smaller recordings. In Fig. 7C we predict reliability as a function of trials from a smaller subset of trials. Focusing on the strongest data mode, we first randomly subsample a set of trials (0.25*T* for Mouse FOV and Monkey, and 0.1*T* for Mouse MP) in order to infer the top-mode 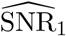, and use that to predict the reliability of disjoint data subsets as a function of the number of trials (up to *T/*2). Our theory 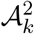 again matches experiment (displayed for 3 different sessions per dataset) and shows how reliability 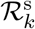 grows with number of trials. In Fig. 7D, we predict the reliability as a function of neurons from a smaller subsample of neurons. We infer the strongest data mode 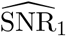 from only a fraction of the neurons (0.15*N* for Mouse FOV and Monkey, 0.1*N* for Mouse MP), and then compare theory and experiment for the subspace reliability of the strongest data mode as a function of the number of neurons *M* included in the analysis (up to *N/*2).

Finally, in Fig. 7E we successfully compare theory and experiment for the subspace reliability of the strongest mode across all sessions and datasets, using data subsets with all *N* neurons and *T/*2 trials. Importantly, we extrapolate the theoretical prediction by inferring the strongest data-mode SNR using just 1*/*4 of the neurons and 1*/*4 of the trials.

A striking property of *A*_*k*_ in Eq. (6) is that it *increases* with increasing neurons *N* at fixed trials *T* (Fig. 4B). This implies that the reliability 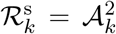 with which any mode *k* is estimated also increases with *N* at fixed *T*, as demonstrated in Fig. 7D, though the increase is mild if *N* is already very large. This increase in reliability may seem surprising because the signal mode ***u***_*k*_ has *N* elements, so one must estimate *more* quantities with the *same* number of trials if one only increases the number of neurons *N* . This improvement stands in direct contrast to the *decrease* in reliability of the *entire* covariance matrix as one increases neurons *N* at fixed trials *T*, as shown in Eq. (3).

A key reason why the reliability of individual signal modes increases, but the reliability of the entire pattern of pairwise correlations decreases, originates from our central assumption that the per-neuron signal-to-noise ratio SNR_*k*_ is constant, leading to a total signal 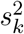 for the mode in Eq. (4) that grows with *N* . This allows the data mode strength 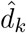 to grow with *N*, outracing the surface of the noise sea (Fig. 4D), and makes the estimate of the signal mode more reliable. In this fashion, large-scale recordings come with a blessing of dimensionality: as one observes more and more neurons, estimates of low-dimensional population geometry, as measured through signal modes, become *more* reliable (because more of the same modes are observed), even as estimates of all pairwise neural correlations become *less* reliable in aggregate.

### A theory of predicting single-trial activity via masked auto-encoding

Inspired by the framework of masked autoencoders He et al. (2021), several approaches to building foundation models in neuroscience partially involve predicting masked neurons from unmasked neurons and trying to minimize the prediction error (see e.g. Le and Shlizerman (2022); Zhang et al. (2024); Willeke et al. (2026)). How this prediction error scales with dataset size and model size then becomes a central issue in building such foundation models.

To begin to address this question in a simple setting, and to illustrate another application of our theory of estimating neural population geometry, we obtain an analytic theory for the prediction error for a simple masked linear autoencoder (Fig. 8A). This autoencoder uses the top 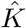 data modes ***û***_*k*_ for 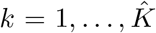 to project neural activity onto a hidden space describing the neural population geometry. Then it uses the same set of modes to expand back to the original neural activity space to fill in predicted neural activity for any neurons that were masked, or zeroed out, at the input. In a situation where the neural activity is generated by our low-dimensional model in Fig. 4, we develop a theory for how the prediction error depends on dataset size (number of neurons *N* and trials *T*), model size (number of hidden neurons 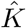), and dataset properties (number of true modes *K* and their true per-neuron SNR_*k*_ values for 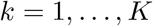). If we let 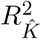 be the fraction of variance explained in predicting a small number of masked neurons on a held-out test trial, when keeping 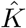 modes, we find (Methods: Autoencoder):

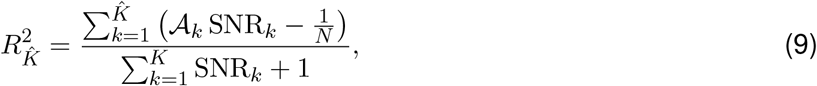

where *A*_*k*_ is the data and signal mode alignment given in Eq. (6) (see Supplement for a refinement of the derivation when data modes mix with multiple signal modes, and see Figure S2 for a validation of this refinement).

**Figure 8:**
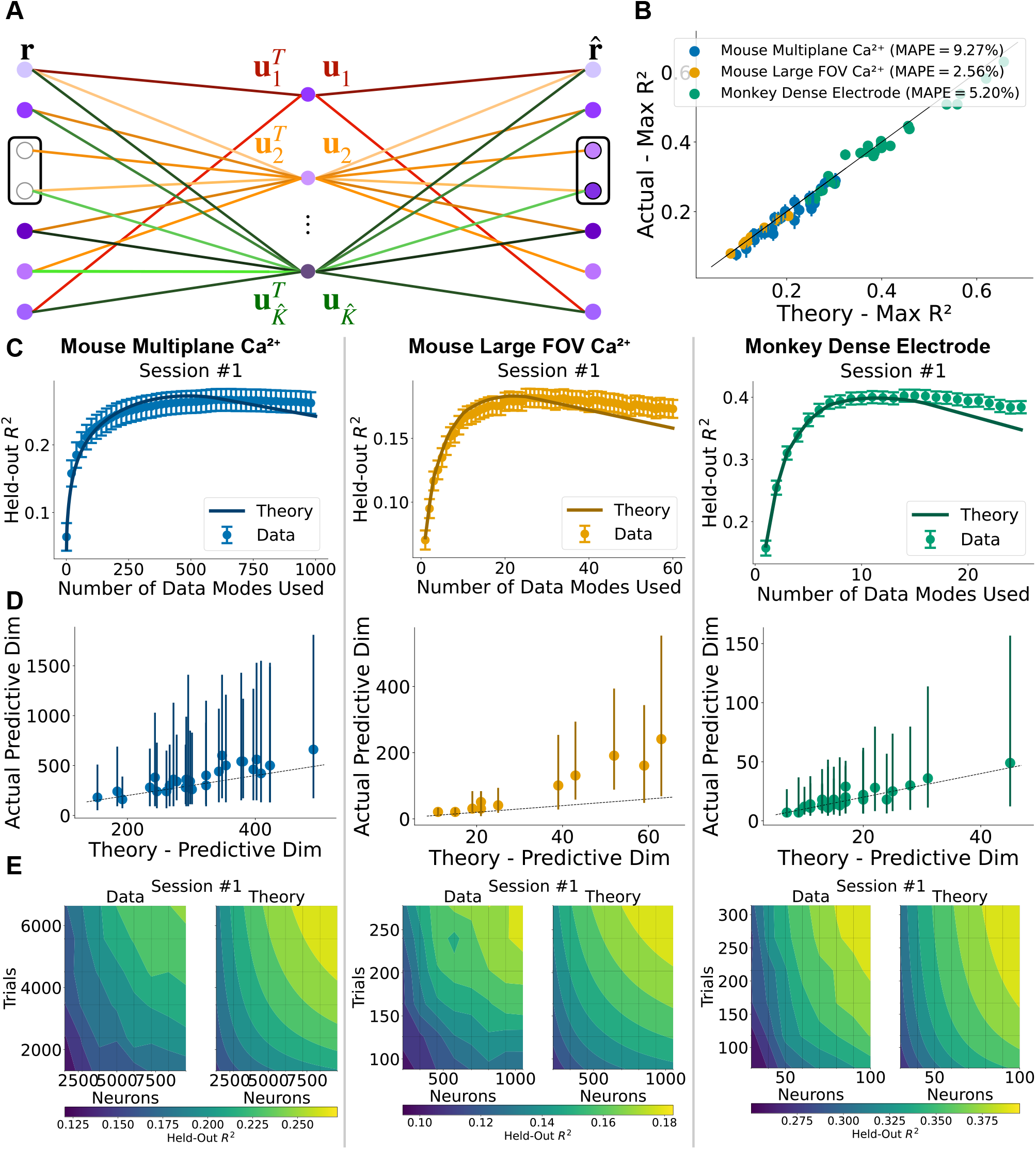
A theory of predicting masked neurons on single trials. **A.**On a new held-out test trial, 2 neurons are masked in the input and predicted in the output, using an autoencoder that employs the top 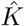 data modes obtained from a training set of separate trials. **B**. Scatter plot of the measured maximal variance explained (R^2^) for predicting single neurons obtained by scanning 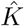 and measuring the best autoencoder performance, versus theoretical predictions obtained via Eq. (9) with 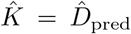 using the threshold in Eq. (10). Results are for all recording sessions, with MAPE shown for each dataset. Vertical black lines display SEM over choice of masked neurons. In (C,D,E) the three columns are for the three well-fit datasets. **C**. Variance explained (*R*^2^) as a function of the number of modes 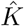 retained with dots showing experiments and solid line showing theory from Eq. (9). **D**. Scatter plot of all sessions showing measured predictive dimensionality 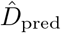, i.e. the optimal number of retained modes prediction, vs that predicted from theory (number of modes with SNR above Eq. (10)). Vertical black lines mark the range of dimensionality that obtains an *R*^2^ within 10% of the maximum. **E**. Contour plots of optimal heldout *R*^2^ vs neurons and trials, *left:* experiment, *right:* theory. Iso-prediction curves are roughly hyperbolas in the space of neurons and trials. Throughout (B-E) we hold out one test trial and predict one masked neuron at a time. For total numbers of masked neurons and test trials for each dataset across panels see Methods: Autoencoder.

This fraction of variance explained has a simple interpretation. The denominator is the total per-neuron variance in units of the noise variance *σ*^2^. It receives a contribution of SNR_k_ from each of the true *K* signal modes plus a unity contribution from the noise. The numerator receives a contribution from each of the 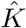 modes kept, and each contribution contains two terms: a positive signal contribution *A*_*k*_ SNR_*k*_ which is weakened by the misalignment between the data and signal mode reflected by *A*_*k*_ *<* 1, and a negative term 1*/N* originating from test noise that propagates through each data mode to the prediction in the autoencoder in Fig. 8A. Intriguingly Eq. (9) shows that it is not enough for an inferred signal mode to be reliable (i.e. have *A*_*k*_ *>* 0) for it to be useful for prediction. Predictive utility demands a higher bar: a mode *k* must have 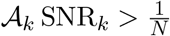 to help improve prediction performance; otherwise it will hurt performance.

This theory provides practical guidance for choosing the number of hidden units 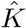 of the autoencoder as follows. In particular, when we view *A* in Eq. (6) as a function of *N*, *T*, and SNR, we can solve the equation 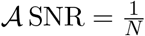 to obtain a threshold SNR_th_ given by (see Methods: Autoencoder),

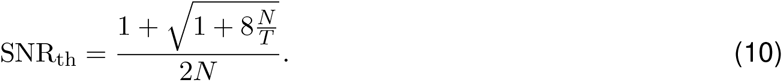

If a mode has SNR_*k*_ *>* SNR_th_, then and only then will it have predictive utility. As a consistency check, as the number of trials *T* → ∞, *A* → 1 in Eq. (6) and 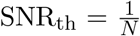 as expected. Interestingly, this means that regardless of the number of trials, for a fixed number of neurons, a signal mode with 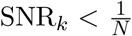 will not be predictive even when its corresponding data mode is perfectly reliable. Quite simply, such a signal mode does not have sufficient SNR to overcome the test noise that it propagates.

We can use the theoretically predicted threshold SNR_th_ to decide whether to retain any mode *k* for held-out neuron prediction in any finite dataset. To do so, we take the data mode strength 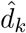, and from this estimate 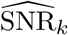 (Methods Eq. (33)), then retain it if it exceeds SNR_th_. Our theory concludes that the optimal number of hidden units in the autoencoder in Figure 8A is the number of data modes satisfying 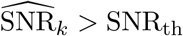. We call this number of modes the *predictive dimensionality*, 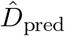. This is the number of dimensions of neural population geometry that are useful for predicting held-out neurons. By substituting SNR_th_ in Eq. (7) for the data mode strengths, we also obtain a closed-form expression for the optimal threshold applied directly to the data mode strengths 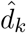, depending only on *N*, *T*, and the inferred noise level, 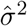 (Methods Eq. (44)).

We directly test our theory in Fig. 8BCDE. In Fig. 8B we compare our theoretically predicted maximal fraction of variance explained for single neuron prediction in Eq. (9) when 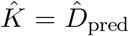, against our empirically obtained maximal variance explained obtained by scanning all possible values of 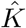 and choosing the highest performance. We see an excellent match between theory and experiment across all sessions of all 3 datasets well fit by our low-dimensional model. In Fig. 8C, focusing on the first session of each dataset, we show a good match between theory and experiment for the fraction of variance explained as a function of 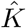. The theory curve is obtained by plotting Eq. (9) as a function of 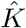 with the estimated 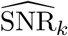 values. Fig. 8D plots the theoretically obtained predictive dimensionality 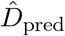 against the optimal 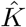 obtained by measuring autoencoder prediction performance, again showing a good theory-experiment match across almost all sessions of all 3 datasets.

Finally, Fig. 8E compares theory and experiment for the optimal fraction of variance explained as a joint function of neurons *N* and trials *T*, for the first sessions of the 3 datasets, again finding a good match. Both in theory and experiment, prediction performance as a joint function of neurons and trials again reveals a striking hyperbolic tradeoff between these two very different experimental resources. If one records *more* neurons, one can get away with *fewer* training trials while still achieving the same predictive performance. Similarly, if one records more trials, one can get away with recording fewer neurons without hurting performance. This tradeoff is again a fundamental consequence of the constancy of the per-neuron SNR_*k*_ of the signal modes in Eq. (4). This leads to the hyperbolic detectability phase transition in Eq. (5) and Fig. 4B. Given this transition, as either neurons or trials are increased, more and more signal modes are detected, and then become useful for prediction, leading to improved performance via Eq. (9).

The scaling laws governing how prediction performance improves jointly with both neurons and trials are of fundamental interest in the new field of foundation models in neuroscience, albeit in much more complex scenarios. Even in our simple theory of such scaling laws for a linear autoencoder model for a single session, the scaling law is complex as it depends on the entire set of per-neuron signal-to-noise ratios SNR_*k*_ for *k* = 1,. .., *K*. For each *N* and *T*, one must compute 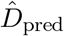 and insert this into Eq. (9) to obtain a theoretically predicted scaling law. However, for some simple and reasonable assumptions about the set of SNR_*k*_, one can derive simple families of scaling laws.

For example, for the three datasets that are well fit by our low-dimensional model, we found that the SNRs follow an approximate power-law SNR_*k*_ ≈ SNR_1_*k*^− *α*^ (Figure 5C and Figure S6). In this case we find (Supplemental Section S9.3) the predictive dimensionality is given by

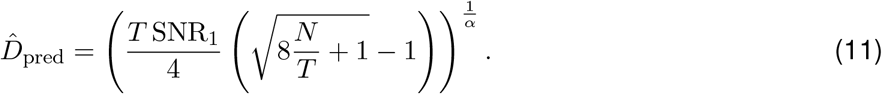

We verify this formula predicts the optimal number of modes to retain almost as well as the full theory, specified by the entire set of inferred SNRs (Figure S6B). In addition, the power-law expression for the SNRs predicts performance across recording sessions comparably to using the precise set of 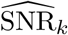 (Figure S6C).

From this formula we derive (see Supplemental Section S9.3) scaling laws for the predictive dimensionality, or equivalently the optimal autoencoder size. We find that in the limit of large numbers of trials, the predictive dimensionality asymptotes to leading order to 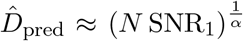. Thus the optimal model autoencoder size will hit a bottleneck proportional to 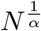 in the limit of infinite trials, as optimal performance saturates due to test noise. On the other hand, in the limit of many neurons, we find that 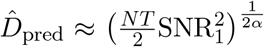. In contrast to the large-trial limit then, the autoencoder’s optimal model size continues to grow in the limit of large *N* .

## DISCUSSION

In summary, we have obtained a predictive theory of experimental design that enables us to determine the accuracy and reliability of various aspects of neural population geometry, as viewed through the simplest and most widespread method of extracting such geometry, PCA. Importantly, our theory provides guidance to neuroscientists to *plan future* experiments, by precisely delineating analytic formulas for how neural dimensionality, accuracy and reliability predictably change as we record more neurons and trials. These analytic formulas reveal exactly what we should measure in past smaller recordings to make such extrapolative predictions to future larger ones. And furthermore, we successfully test these predictions across diverse recording modalities, species, tasks, and brain regions. Our theory reveals several lessons about the experimental design and analysis in the era of large-scale neural recordings.

### The fundamental role of participation-ratio based dimensionality

First, the dimensionality of neural activity has drawn great interest in systems neuroscience (Gao et al. (2017); Altan et al. (2021); Sadtler et al. (2014); Gallego et al. (2017); Stringer et al. (2019); Manley et al. (2024); Rigotti et al. (2013); Fusi et al. (2016)). However, there is no consensus on how dimensionality should be defined. The choice of definition necessarily relies on the question the dimensionality measure is designed to answer. In the case where the question is how reliable is the *entire pattern* of neural correlations across a large population, our results show that the choice of dimensionality measure is unambiguous: it is the participation ratio based measure of data dimensionality 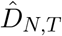. Specifically, we derived a remarkably simple formula: the total replication error of pairwise correlations across two equal-trial recordings is twice the ratio of 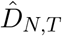 to the number of trials (Eq. (3)). This reveals a very practical experimental insight that only when the measured data dimensionality 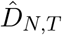 is much less than the number of trials *T*, will the measured neuronal correlation matrix *Ĉ*_*N,T*_ replicate across repeated experiments. This provides a useful diagnostic for experimentalists to ascertain the reliability of their measured neural correlations from a *single* experiment, *without* having to conduct any new repeated set of experiments.

### The predictable growth of dimensionality with recording size

Second, given the relationship between dimensionality and reliability, it is extremely useful to be able to *predict* how measured data dimensionality increases (if at all) with more recorded neurons and trials collected in *future* experiments, *without* having to do those experiments. We answer this question in Eq. (1). This shows, remarkably, that measured data dimensionality grows with neurons and trials in a lawful manner, governed by a *single* number *D*_*∞*_, the true dimensionality obtained by recording the entire relevant neural population for infinitely many trials. This provides several useful guidelines for experimentalists. First, one can extract *D*_*∞*_ from a measured 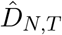 by inverting Eq. (1) to obtain the true dimensionality of the *entire* neural population (see Methods Eq.(17) for the explicit inversion formula). Second, one can *predict* the data dimensionality 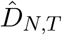 in *future larger* scale recordings with *N > M* neurons and *T > P* trials from the measured data dimensionality 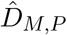 in *past smaller* scale recordings with only *M* neurons and *P* trials (see Methods Eq.(18) for the explicit prediction formula).

We note that Hu and Sompolinsky (2022) employed free probability cumulant relations to derive two formulas equivalent to Methods Eqs. (15) and (16), for participation-ratio dimensionality as a function of subsampling trials and neurons, respectively. Concurrently with our work, Chun et al. (2026) derive a similar scaling law for the data dimensionality via direct moment computation and a second-order Taylor expansion of the estimator ratio. In contrast, our derivation proceeds by evaluating fourth-order mixed traces exactly using free-probability identities for freely independent matrices.

Throughout this work, as an extended preprocessing step beyond z-scoring each neuron’s activity over trials, we also normalize the neural population activity vector on each trial to have unit norm, to constrain the overall squared neural activity (i.e. power). In Supplemental Section S7, we show that trial-to-trial variability in the overall power of neural activity reduces the reliability of the sample covariance predictably. We derive a novel, but more complex theory that goes beyond Hu and Sompolinsky (2022) and Chun et al. (2026), to incorporate trial-to-trial variability in overall power. Our extended theory successfully predicts data dimensionality (Figure S4) and covariance matrix reliability (Figure S5) without any normalization of overall population power.

### The curse of dimensionality for the entire neural population

The combined lawful growth of data dimensionality with neurons and trials (Eq. (1)), and its fundamental role in governing the reliability of neural correlations (Eq. (3)), together reveal a curse of dimensionality for the *entire* pattern of neural correlations. In particular, if one records *more* neurons *N* without a concomitant increase in the number of trials *T*, then measured neural correlations will become *less* reliable because the data dimensionality numerator 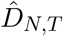 in Eq. (3) will increase. Thus more neurons requires more trials to maintain overall reliability.

However, our theory can provide practical guidance for the design of future experiments in order to achieve a criterion level of reliability. For example, given a measured data dimensionality 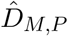 in a small recording of only *M* neurons and *P* trials, suppose we wish to obtain a much larger recording of *N > M* neurons from the same neural population. How many trials *T* would we then need to achieve a given replication error RepErr (*Ĉ*_*N,T*_) for the entire neural covariance matrix *Ĉ*_*N,T*_ ? To answer this, we can use our dimensionality extrapolation formula Methods Eq. (18) to predict 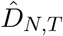 from 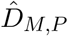, insert this prediction into Eq. (3) and solve for *T* for any desired replication error.

We note also that the curse of dimensionality for the entire neural correlation matrix can be mitigated by the fact that the growth of measured data dimensionality 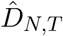 with neurons *N* and trials *T* saturates to a fixed value *D*_∞_ as soon as both *N* and *T* are substantially larger than *D*_∞_ (Eq. (1) and Fig. 2A). For example, suppose we could record from *all N* ≈ 1.5 − 3 million (Young et al. (2013); Higo et al. (2016)) neurons in a macaque motor cortical area controlling arm movements. Assuming the unrecorded neurons share common modes with the recorded ones (see Assumptions and Limitations below), our theory reveals that we won’t need as many as 3 million trials to reliably estimate the *entire* neural covariance matrix. Instead we only need about 10 × *D*_∞_ trials. Indeed in macaque motor cortex, our measurements reveal that *D*_∞_ is less than 50 (Fig. 2BCD, third column), which substantially mitigates the curse of dimensionality.

### The blessing of dimensionality for neural population geometry

While the entire neural correlation matrix suffers from a curse of dimensionality (recording more neurons requires more trials to maintain reliability), interestingly, the low-dimensional geometry enjoys a *blessing* Donoho (2000) of dimensionality: recording *more* neurons actually requires *fewer* trials to maintain reliability. In particular, we find that the (subspace) reliability ℛ_*k*_ of a single data mode corresponding to a shared signal across the neural population, is well captured by its squared alignment 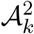, with *A*_*k*_ given by Eq. (6). This formula clearly exhibits a striking tradeoff between two very different experimental resources: neurons and trials. As one records either more neurons or more trials, one can detect successively weaker signal modes with smaller per-neuron signal-to-noise ratios (as defined in Eq. (4)). Moreover, for any given signal mode, recording more neurons *N* requires recording *fewer* trials *T* to estimate it with the same reliability.

Intuitively, the fundamental reason that one can estimate an individual signal mode *k*, with *fewer* trials given *more* neurons, rests on the crucial assumption that the mode’s per-neuron signal-to-noise ratio SNR_*k*_ is an intrinsic constant inherent to the neural population and recording modality. This then implies that the total signal in the mode *grows* with the number of recorded neurons, as reflected in our crucial modeling assumption in Eq. (4). Thus as one observes more of the mode, by recording more neurons, its overall signal increases, and therefore fewer trials are required to reliably estimate it.

Our theory again provides practical guidance for designing future experiments to estimate neural signal modes. First, to estimate a signal mode *k* with a given SNR_*k*_, Eq. (4) tells us how many neurons and trials we will need to achieve a given reliability. Second, from a given recording, we can form an estimate 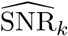 of the per-neuron signal-to-noise ratio of any potential signal mode by measuring its data mode strength 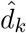 and then inverting Eq. (7) (see Methods Eq. (33) for the explicit inversion procedure). Third, with the estimate 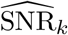 in hand, and assuming its constancy, we can *predict* the reliability of the mode in *future* recordings with *more* neurons and trials using the square of Eq. (6).

Taken together, the blessing of dimensionality for neural population geometry provides excellent news for the future of systems neuroscience. With improved technology to record more neurons in the future, we will be able to use fewer trials of any given trial type to reliably estimate neural population geometry, paving the way to new forms of experiments with many more diverse trial types.

### The rudiments of a theory of foundation model scaling in neuroscience

Inspired by recent trends in artificial intelligence, a modern development in systems neuroscience is the creation of foundation models that are capable of modeling neural activity across multiple sessions, tasks, and subjects (see e.g. Le and Shlizerman (2022); Zhang et al. (2024); Willeke et al. (2026)). Often, the error in predicting the activity of held-out single neurons decreases as a power law with both neurons and training trials, while the optimal model size increases as a power law (Willeke et al. (2026)). Such scaling laws Kaplan et al. (2020) have driven significant technological investment in artificial intelligence, notably in large language models, where we have begun to develop a theory of their origin Cagnetta et al. (2026). However, there exists as of yet no mathematical theory that explains the empirically observed scaling laws for foundation models in neuroscience. Given prediction performance improves with model size and recording size, it is important to understand why.

We have developed the beginnings of such a theory limited to a single session, and a very simple linear masked autoencoder model for predicting held-out neurons. Our theory analytically predicts the prediction performance (Eq. (9)) and optimal model size (via Eq. (10)) as a joint function of the number of neurons and trials used in training. Moreover, it reveals the origin of power-law scaling in optimal model size with neurons and trials to be a natural consequence of power-law distributions of per-neuron signal-to-noise ratios of neural modes in neural data (Figure 5C and Figure S6A). Such a power law in data leads to a power law in the optimal model size (Eq. (11)). The basic intuition is that as one increases neurons and trials, one can detect more and more lower SNR signal modes in the data, and use these detected signal modes to build a bigger and better performing autoencoder.

Indeed, the number of hidden neurons in the optimally sized autoencoder constitutes another notion of dimensionality, which we call the predictive dimensionality. It is basically the number of signal modes that are useful for predicting the activity of held-out neurons on held-out trials, which is a key part of the training objective of foundation models in neuroscience. Importantly, we do not find a consistent relationship between the participation-ratio based dimensionality and the predictive dimensionality (Figure S3). Thus these two notions of dimensionality reflect distinct properties of neural data, which is natural, as each arises as a principled answer to a distinct question: the reliability of the entire neural correlation matrix versus the predictive utility of individual modes of neural population geometry.

### Assumptions and Limitations

It is important to note that we make several assumptions about the structure of neural data to derive our theory, assumptions that necessarily limit the applicability of our theory. First, we assume that all trials are independent of each other. This precludes the possibility of quantitatively analyzing dynamical processes that exhibit strong correlations over time. However, there does exist random matrix theory that deals with the analog of correlated trials (Yao (2012)), and such work may potentially be used to extend our theory. However, typical dynamical systems exhibit strong temporal correlations within a certain auto-correlation time, while neural activity at time separations larger than this time are approximately independent. In this sense our theory may be approximately applied with the number of effective trials equal to the total recording time divided by the auto-correlation time, with an appropriately increased variance reflecting temporal correlations within the auto-correlation time.

Second, to make predictions about the results of analyses on neurons that have not yet been recorded, we necessarily must make an assumption about the relationship between the unrecorded neurons and the recorded ones. Our key assumption is that the neural modes across the entire population are distributed indistinguishably across recorded and yet-to-be-recorded neurons (See Supplemental Section S5.2 for the precise mathematical statement of this assumption and Fig S1 for experimental validation). This assumption could be violated, for example, if the unrecorded neurons have distinct functional properties from the recorded neurons (e.g. as different cell-types or from different brain regions), and therefore have their own neural modes, distinct from the modes of the recorded neurons. In essence, if the unrecorded neurons bear no relation to the recorded ones, one can say nothing about them. Nevertheless, the isolation of our simple assumption of indistinguishably shared modes across recorded and unrecorded neurons powerfully enables us to obtain a predictive theory of experimental design for neural population geometry, as demonstrated across all our theory-experiment comparisons.

The above two assumptions are all that is required to obtain a predictive theory of experimental design for data dimensionality and the accuracy and reliability of all neural correlations, as demonstrated in Figures 2 and 3. However, to obtain a predictive theory of the accuracy and reliability of individual neural modes, we make a third assumption about how these modes are generated, through a model of shared variability driven by a small number of latent modes, and additive private noise of common variance (Fig. 4). There are several ways this model could be violated. First, the number of latent modes *K* may be quite large, as large as the number of neurons *N* and trials *T* . Second, the private noise may not be additive, or homogeneous across neurons. Nevertheless, our theory provides a first step to test whether a dataset satisfies our low-dimensional model assumptions: namely check if the noise sea has a Marchenko-Pastur shape (Fig. 5AB). Thus, like any good theory, it provides a self-contained test of its own regime of validity. Interestingly, the human fMRI dataset does not pass this test, despite the fact that it has relatively low data dimensionality 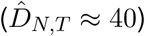, suggesting assumptions of homogeneous additive private noise or uncorrelated trials are violated in these data. However, in the other 3 datasets that passed the validity check, we were able to successfully confirm all predictions of our theory (Figures 6,7, and 8). In future work there may be ways to go beyond our assumption of low dimension, using more sophisticated random matrix theory that assumes the number of latent modes *K* is proportional to the number of neurons or trials (Bun et al., 2018; Landau et al., 2023), or using finite dataset size corrections (Bai and feng Yao, 2008).

Fourth, our theory of scaling laws for held-out neural prediction is only performed in a limited setting: a simple linear masked autoencoder on a single session. However, while simple, our theory fulfills an important prerequisite to deriving theories of scaling laws for more complex architectures on more diverse data spanning multiple sessions and subjects. Also, already in our simple setting, we observe power laws in model size and performance, and we relate them to power laws present in the neural population geometry itself. An intriguing future direction lies in answering what shared aspects of neural population geometry across subjects allow foundation models in neuroscience to benefit from integrating across subjects.

In summary, as progress in recording technologies continues to thrust systems neuroscience squarely into a new era of many neurons and potentially few trials, we will need new predictive theories that can help us design experiments *before* the data is even collected. Our work provides such a theory for the extraction of neural population geometry using PCA, and reveals a remarkable blessing of dimensionality that will enable us to obtain veridical results in *higher* dimensional recordings with *more* neurons, using *fewer* trials.

## METHODS

### Key resources table

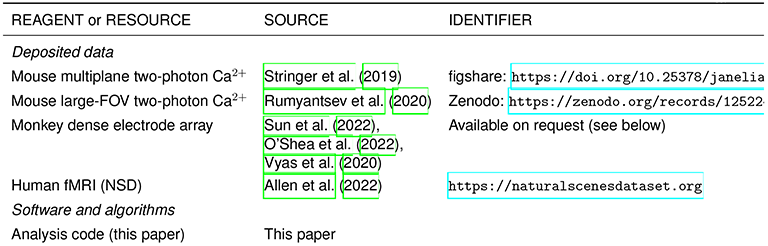

### Resource availability

#### Lead contact

Further information and requests should be directed to and will be fulfilled by the lead contact, Itamar D. Landau.

#### Materials availability

This study did not generate new materials.

#### Data and code availability

- This study analyzes existing, publicly available datasets except where noted. The mouse multiplane two-photon calcium imaging data Stringer et al. (2019) are available at figshare (DOI: 10.25378/janelia. The mouse large-FOV data Rumyantsev et al. (2020) are at Zenodo (https://zenodo.org/records/12522419). The human fMRI data Allen et al. (2022) are available through the Natural Scenes Dataset (https://naturalscenesdataset.org). The monkey dense-electrode data Sun et al. (2022); O’Shea et al. (2022); Vyas et al. (2020) are not publicly archived and are available from the corresponding authors of that study upon reasonable request.
- Any additional information required to reanalyze the data reported in this paper is available from the lead contact upon request.

### Method details

#### Datasets and Preprocessing

We evaluated our theory across four datasets spanning species, brain regions, and recording modalities. From each recording session we extract an *N* × *T* matrix of trial-averaged activity, where *N* is the number of neurons (or voxels) simultaneously recorded and *T* is the number of distinct trials in the session (whether the stimuli or behavioral conditions are repeated or unique). Within-trial averaging windows are specified per dataset below. Z-scoring and trial-normalization are described below; all other prior preprocessing follows the original publications. Per-session values of *N*, *T*, as well as a number of key experimentally inferred values are tabulated in Supplemental Section S2 (Tables 1–4).

1. **Mouse multiplane two-photon calcium imaging** (mouse V1; Stringer et al. (2019)). Seven mice, 28 recording sessions of natural-image presentation, with ∼ 10,000 neurons per session. Most sessions presented each of 2800 images with two randomly ordered repetitions, for 0.5 s each; activity was averaged over the first two frames after stimulus onset. See Table 1 (label *Mouse V1 multiplane Ca*^*2+*^) for stimulus descriptions.
2. **Mouse large-FOV multi-beam two-photon calcium imaging** (mouse V1; Rumyantsev et al. (2020)). Eleven mice each with a single session viewing one of two oriented gratings per trial (*±*6° or *±*30° separation), with ∼500–2000 neurons and ∼200–300 trials per session. Stimuli were presented for 2 s; activity was averaged over the final 1.5 s. *T* pools both grating orientations. See Table 2 (label *Mouse V1 Large FOV Ca*^*2+*^).
3. **Monkey dense electrode-array recordings** (monkey M1 and PMd; Sun et al. (2022)). Two rhesus macaques (P and V) performing a center-out reach task with ≥8 movement directions, 24 sessions, ∼100–300 neurons and ∼200–500 trials per session. *T* pools all reach directions. See Table 3 (label *Monkey dense electrode*).
4. **Human fMRI** (visual cortex; Natural Scenes Dataset, Allen et al. (2022)). Eight subjects, left hemisphere, voxelwise responses to 750 natural images, pooled across visual ROIs (V1v, V1d, V2v, V2d, V3v, V3d, hV4). This dataset is found to be a failure case for the low-rank plus noise model and is treated as such throughout (see Results and Discussion). See Table 4 (label *Human fMRI*).

We applied two preprocessing steps to every session, in the following order.

##### Per-neuron z-scoring

Each neuron’s response was z-scored across the *T* trials, so that every row of the activity matrix has zero mean and unit variance. Neurons whose across-trial standard deviation fell below a small threshold (effectively silent or non-responsive units) were excluded prior to z-scoring.

##### Per-trial normalization

Each trial’s population activity was normalized as follows: For each trial *t* we computed the across-neuron mean square activity 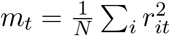, normalized these to unit average, 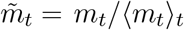, to obtain a per-trial scale factor, 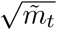, and divided each trial by its scale factor, 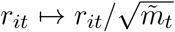. This leaves the average per-trial power (mean square activity) unchanged while equalizing trial-to-trial scale.

In Supplemental Section S7 we derive theory for the data dimensionality, and for the accuracy and reliability of the data covariance that incorporates trial-to-trial scale fluctuations. Figures S4 and S5 compare theory to data without trial normalization.

#### Notation and setup

Throughout, *R* ∈ ℝ^*N*×*T*^ denotes the preprocessed activity matrix of *N* neurons (or voxels) across *T* trials, and the *data covariance* is

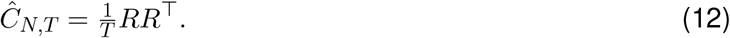

We model the trials as independent draws from a fixed *neural covariance C*_*N*_, writing 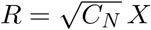, where *X* has i.i.d. zero-mean, unit-variance entries. The data covariance is then 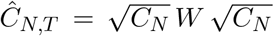 with 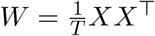 is a pure noise covariance. We write the neuron-to-trial aspect ratio as

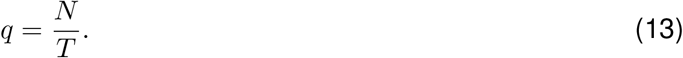

We refer to the eigenvalue–eigenvector pairs (*d*_*k*_, ***u***_*k*_) of *C*_*N*_ as *neural modes* and to the pairs 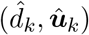 of *Ĉ*_*N,T*_ as *data modes*, so that 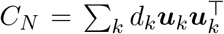 and 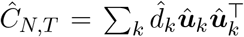. When the neural covariance carries low-rank signal structure (introduced in Section Low-rank below), we distinguish *signal* modes from the flat *noise* floor, and the data spectrum into *outlier* data modes that rise above the noise floor and the *bulk*, or the “noise sea”, that does not.

We write *τ* for the dimension-normalized trace, 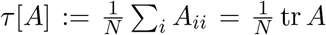, and use it throughout in place of unnormalized sums so that quantities have well-defined limits as *N, T* → ∞ at fixed *q*. The free-probability machinery and trace-moment identities used in the derivations are collected in Supplemental Section S4.

We quantify the population geometry of a covariance by its participation-ratio dimensionality. Applied to the neural covariance, we have the *neural dimensionality* :

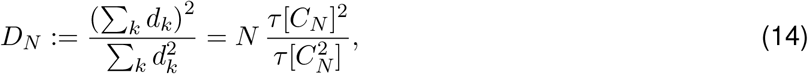

and the *data dimensionality* 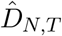 defined similarly in terms of the eigenvalues of *Ĉ*_*N,T*_ . *D*_*N*_ counts the effective number of neural modes; it is model-free and does not separate signal from noise (pure i.i.d. noise attains the maximum *D*_*N*_ = *N*).

#### Dimensionality scaling and extrapolation

Finite sampling distorts the data dimensionality relative to the neural dimensionality. We show (Supplemental Section S5) that the data dimensionality and neural dimensionality obey a reciprocal-sum law with the number of trials:

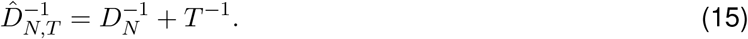

Under the assumption that neural modes are distributed identically across recorded and as-yet unrecorded neurons, we show that dimensionalities obey a similar reciprocal-sum law with regard to numbers of neurons:

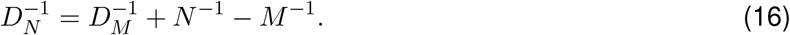

Taking *M* → ∞ and substituting into Eq. (15) yields Eq. (1) in the main text.

Solving instead for *D*_∞_ yields

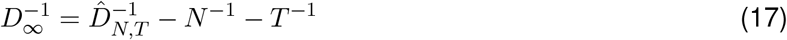

Here we state the operational form that can be applied directly to obtain the data dimensionality at (*N, T*) predicted from a subsample of *M* ≤ *N* neurons and *P* ≤ *T* trials:

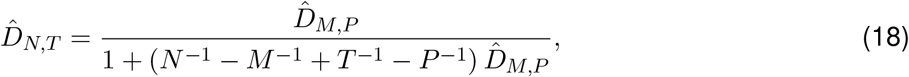

which is equivalent to the reciprocal-sum rule: 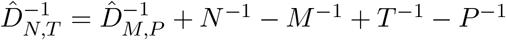.

##### Subsampling and extrapolation procedure

Each panel of Figure 2 tests the extrapolation formula, which predicts the data dimensionality 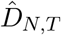 at a *target* size (*N, T*) from a measurement 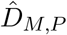 taken on a smaller *anchor* subsample of *M* = *f*_*N*_ *N* neurons with *f*_*N*_ ≤ 1 and *P* = *f*_*T*_ *T* trials with *f*_*T*_ ≤ 1. The target is swept along the trial axis (panel B), the neuron axis (panel C), or both (panel D). In panel E the single target is the full session itself, (*N, T*)= (*N*_full_, *T*_full_).

At each target we compare the *predicted* dimensionality against the *observed* dimensionality measured directly at that target. The observed value is averaged over independent random draws of the target subsample, with error bars giving the standard deviation across draws (vanishing in panel E, where the target is the full session and only one draw exists). The predicted value is obtained by extrapolating from the anchor: we evaluate the extrapolation formula at the anchor mean and at the anchor mean *±* one standard deviation, propagating the anchor’s across-draw uncertainty through the formula rather than computing the prediction distribution directly. Because the anchor coefficient of variation does not exceed ∼ 0.1 in any panel (and is most often far smaller) and the extrapolation maps are near-affine over this range, the difference between the two procedures is negligible. Panel D is a colormap, on which this variability is computed but not displayed.

| Figure 2 Panel | Anchor $(f_N, f_T)$ | Repetitions | | | |
| --- | --- | --- | --- | --- | --- |
|  |  | Mouse MP | Mouse FOV | Monkey | Human |
| B (sweep $T$) | $(1, \frac{1}{4})$ | 10 | 20 | 20 | 20 |
| C (sweep $N$) | $(\frac{1}{10}, 1)$ | 10 | 20 | 20 | 10 |
| D (sweep $N, T$) | grid corner | 50 | 50 | 200 | 100 |
| E ($N_{\text{full}}, T_{\text{full}}$) | $(\frac{1}{4}, \frac{1}{4})$ | 10 | 10 | 10 | 10 |

Anchor sizes are fractions of the full session, (*M, P*)= (*f*_*N*_ *N*_full_, *f*_*T*_ *T*_full_); for panel D the anchor is the smallest (*M, P*) on the displayed grid (corresponding to the bottom-left corner of the panel). Repetitions count both the number of independent anchor draws used to compute the prediction band and the number of repeated target draws to compute the observation band.

#### Accuracy and reliability of the data covariance

We assess the data covariance *Ĉ*_*N,T*_ by two criteria: its *accuracy* relative to the neural covariance *C*_*N*_, and its *reliability* across disjoint subsets of trials. Both reduce to a dimensionality-over-trials ratio; derivations are given in Supplemental Sections S6.1–S6.2.

##### Accuracy

The *estimation error* is defined as the normalized squared Frobenius distance between the data and neural covariances,

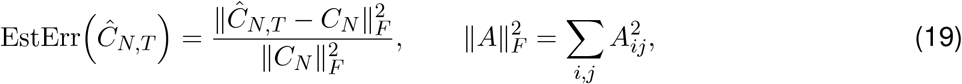

with 0 denoting perfect estimation.

We show that its expected value is the ratio of the neural dimensionality to the number of trials,

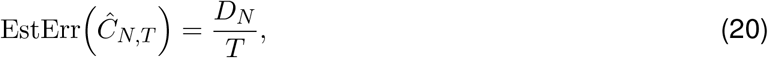

up to an 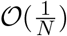 relative correction.

##### Reliability

Because *C*_*N*_ is unobservable, accuracy cannot be measured directly. We therefore define the *replication error* between two data covariances 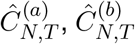 computed from disjoint sets of *T* trials,

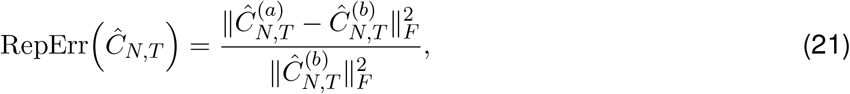

with 0 denoting perfect replication.

We show that its expected value is twice the ratio of the data dimensionality to the number of trials (see Supplemental Section S6.2 for derivation),

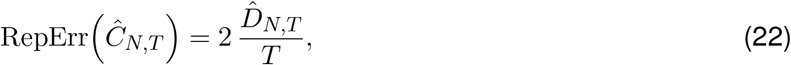

up to a relative correction of 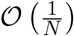.

Unlike accuracy, replication error is directly measurable from disjoint data splits.

Substituting the dimensionality extrapolation formula (18) into Equation (22) yields a prediction of the full-data replication error from a subsample (derivations in Supplemental Section S6.2). From *P* ≤ *T* trials and *M* ≤ *N* neurons,

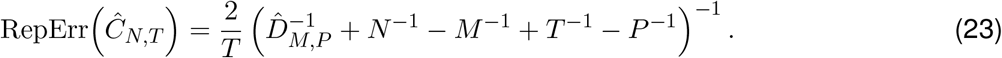

##### Subsampling and extrapolation procedure

Each panel of Figure 3 tests the extrapolation formula that predicts the replication error RepErr(*Ĉ*_*N,T*_) at a *target* size (*N, T*) from a measurement of 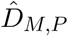 on a smaller *anchor* subsample of *M* = *f*_*N*_ *N* ≤ *N* neurons and *P* = *f*_*T*_ *T* ≤ *T* trials, as in the dimensionality procedure above (Section Dim Procedure). The target (*N, T*) is swept along the trial axis (panel A), the neuron axis (panel B), or both (panel C). Panel D uses the full session split as the target, 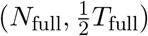. The predicted error-band is again obtained (in panels A,B,D) by propagating the anchor’s across-draw mean and *±* one standard deviation through this map; since the replication error is a *linear* readout of the extrapolated dimensionality, the approximate agreement of the two bounds established for the dimensionality extrapolation carries over directly.

The observed replication error at each target is computed from two disjoint, equal-size sets of *T* trials drawn from the session; the displayed trial count *T* in the figure axes is the number of trials *in each split*, so the full-session measurement uses all *N*_full_ neurons and *T* = *T*_full_*/*2 trials per split. It is averaged over independent random draws of the split, with error bars giving the standard deviation across draws. Unlike the dimensionality case, this remains stochastic even at the full session, because the two trial sets are a random split; panel D therefore retains observed error bars. Both predictions and observations are averaged over the same number of disjoint-split repetitions listed below. Panel C is a colormap, on which the across-draw variability is computed but not displayed.

| Figure 3 |  | Repetitions |  |  |  |
| --- | --- | --- | --- | --- | --- |
| Panel | Anchor $(f_N, f_T)$ | Mouse MP | Mouse FOV | Monkey | Human |
| A (sweep $T$) | $(1, \frac{1}{4})$ | 10 | 20 | 20 | 10 |
| B (sweep $N$) | $(\frac{1}{10}, 1)$ | 10 | 20 | 20 | 10 |
| C (sweep $N, T$) | grid corner | 10 | 30 | 50 | 10 |
| D $(N_{\text{full}}, \frac{1}{2}T_{\text{full}})$ | $(\frac{1}{4}, \frac{1}{4})$ | 10 | 10 | 10 | 10 |

Anchor sizes are fractions of the full session, (*M, P*)= (*f*_*N*_ *N*_full_, *f*_*T*_ *T*_full_); for panel C the anchor is the smallest (*M, P*) on the displayed grid (corresponding to the bottom-left corner of the plot). Repetitions give the number of independent anchor draws and target draws.

#### Low-rank plus noise model

The low-rank plus noise model assumes *K* ≪ *N* shared signal modes on top of independent identically distributed noise with variance *σ*^2^, giving a neural covariance

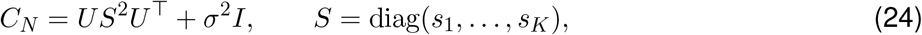

where the columns of *U ∈* ℝ ^*N* × *K*^ are orthonormal *signal mode vectors* and 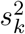 is the strength of signal mode *k*. We parameterize each signal mode by a *per-neuron* signal-to-noise ratio SNR_*k*_, defined so that

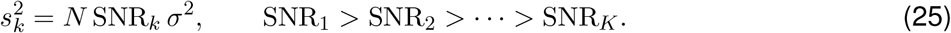

The neural mode strengths (eigenvalues of *C*_*N*_) are 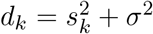 for *k* ≤ *K* and *d*_*k*_ = *σ*^2^ for the rest, and we take *K* = *O*(1) as *N* → ∞. We hold SNR_*k*_ fixed as *N* grows, so signal strength scales linearly with *N* (Eq. (25)).

#### Finite-sampling distortion of the spectrum

Sampling *Ĉ*_*N,T*_ from *T* trials inflates each signal mode and spreads the noise into a Marchenko–Pastur (MP) noise sea. For every mode above the critical ratio 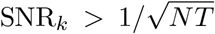 (Eq. (5) in the main), the corresponding data mode rises above the noise sea with expected strength

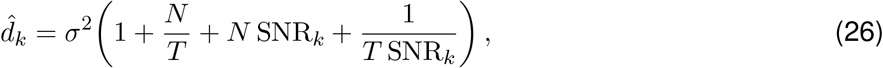

while modes below the critical ratio are absorbed into the noise bulk and are unrecoverable. See derivation in Supplemental Section S10.

#### Fitting the low-rank model to data

We fit the model in two stages — first we fit the noise variance and report a goodness-of-fit statistic, then we infer the per-mode SNRs for each potential signal mode — and fit an SNR power-law exponent.

##### Noise variance

We estimate the noise variance by matching the median data mode. That is, we choose the noise variance, 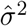, under which i.i.d. noise would yield the same median Gavish and Donoho (2014):

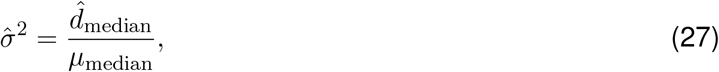

where 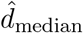 is the median data mode strength and *µ*_median_ is the median of the standard MP distribution of aspect ratio 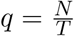, obtained numerically from

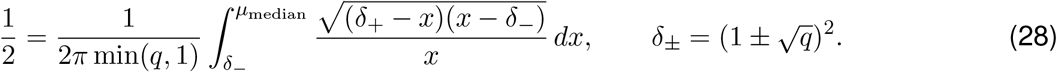

The fitted MP noise bulk is displayed for session #1 for each dataset in Figure 5A.

Note that for the case *N > T*, we do not consider the *N* − *T* zero eigenvalues due to rank deficiency to be among the data modes.

##### Goodness of fit

To assess how well each dataset conforms to the model we compare the empirical distribution of bulk data modes to the MP distribution of aspect ratio 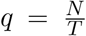 using the 1-Wasserstein distance between their cumulative distributions *F*_emp_ and *F*_MP_ (Figure 5B).

The Wasserstein-1 distance,

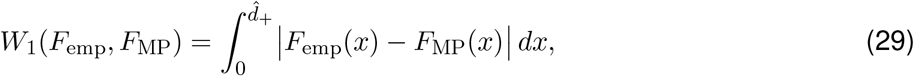

measures the area between the curves of the CDFs. This distance scales with the overall scale of the eigenvalues, and because the medians are matched, it is bounded by 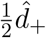, where 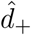 is the upper boundary of the noise sea (see following section; Equation (31)). In order to obtain a unit-free measurement that is comparable across datasets, we normalize the Wasserstein distance:

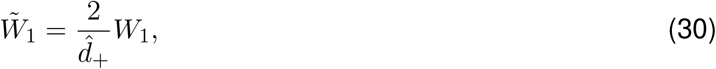

so 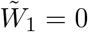 is a perfect match and 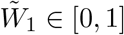.

The 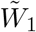 score for every session of each dataset is displayed in Figure 5B.

##### Signal modes and inferred SNR

Every data mode whose strength exceeds the noise sea is treated as a potential signal mode. The upper boundary of the noise sea, i.e. the upper MP bulk edge, is given by

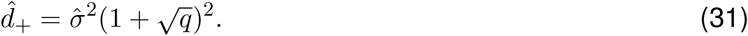

For each mode *d*_*k*_ below 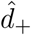 we invert Eq. (26) to find the inferred 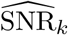. For notational clarity, we do so in two stages. First, subtracting the midpoint of the noise sea 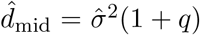 and rescaling by 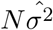, yields

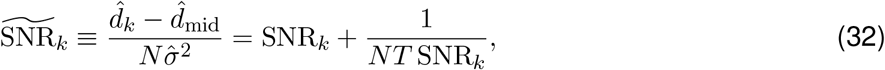

and finally, inverting this formula yields the inferred SNR as the positive quadratic root,

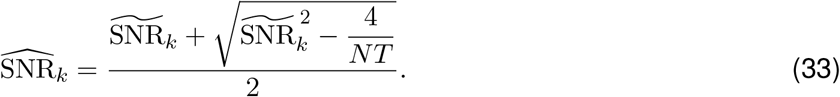

At the ceiling 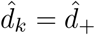 this gives 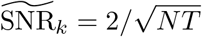, so the radicand is non-negative for all retained modes. Note also that substituting this value into the previous equation gives the critical signal-to-noise ratio 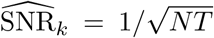. Substituting 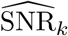 back into Eq. (26) at a different (*N, T*) predicts the data mode strength at that data size.

##### SNR power-law exponent

Across datasets we observe the inferred SNRs fall off approximately as a power law in mode index. We fit 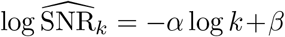 by ordinary least squares over the retained signal modes (Figure 5C); the slope magnitude *α* is the SNR power-law exponent and SNR_1_ ≡ e^*β*^ is the fitted top SNR used in the predictive-dimensionality scaling (Held-out neuron prediction). The linear fit in log-log scale is displayed for Session #1 for each of the *Mouse V1 multiplane Ca*^*2+*^, *Mouse V1 Large FOV Ca*^*2+*^, and *Monkey dense electrode* datasets in Figure 5C.

#### Signal geometry under finite sampling

Under the low-rank plus noise model, finite sampling distorts the inferred signal geometry in two ways: it inflates the signal mode eigenvalues, deforming the signal subspace, and it misaligns the signal mode vectors. We state the operational predictions here and derive them in Supplemental Section S8.

##### Signal mode prediction and geometry

Given an estimate of a signal mode 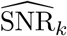 from any dataset size, (*M, P*), we obtain a prediction for the corresponding data mode eigenvalue at arbitrary *N* and *T* . For 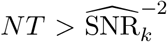, we use Equation (26), and for 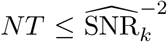, the prediction is simply the upper boundary of the noise: 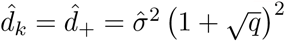.

Figure 6A displays data mode strengths as a function of *N* . Predictions are obtained by inferring the 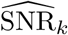 from the full data. Observations are obtained by repeatedly computing the eigendecomposition of 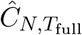 from random samples of *N < N*_full_ neurons. Figure 6B displays extrapolated data modes, predicted by inferring 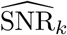 from repeated random subsamples of 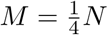 and 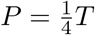.

We assess the shape of the signal subspace with the anisotropy, measured as the ratio of successive signal eigenvalues. Taking the ratio of Eq. (26) for consecutive modes,

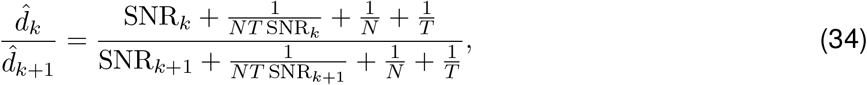

which approaches the true ratio SNR_*k*_*/*SNR_*k*+1_ as *N, T* grow. Setting Eq. (34) to a fixed value *R*_*k*_ gives the iso-ratio curve in the neuron–trial plane,

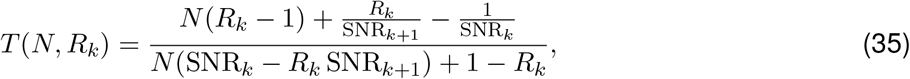

along which the data mode anisotropy is held constant.

Figure 6C displays contours of equal anisotropy as a function of neurons and trials together with those predicted by inferring the 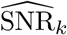 from the full data.

##### Signal mode-vector deformation

The low-rank plus noise model also predicts that finite sampling rotates each data mode vector ***û***_*k*_ away from its corresponding signal mode vector ***u***_*k*_. We show that their alignment, *A*_*k*_, as measured by squared overlap is predicted to be

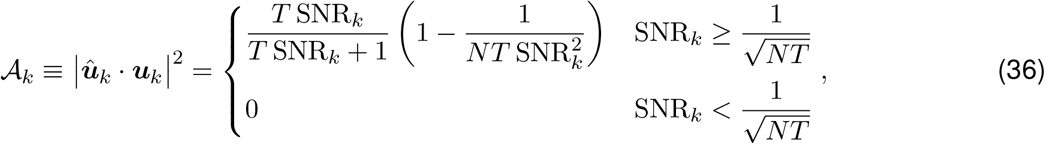

and the cross-mode overlap |***û***_*k*_ · ***u***_*j*_|^2^ for *j* ≠ *k* vanishes in the large-*N* limit. See derivation in Supplemental Section S10.

##### Data mode reliability

The alignment *A*_*k*_ (Eq. (36)) depends on the neural mode vector ***u***_*k*_, which is unobservable. We therefore define the measurable *data mode reliability* ℛ_*k*_ as the squared overlap between the *k*th data mode vectors recovered from two independent equal-trial data samples,

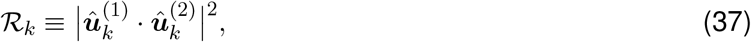

where 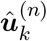 is the *k*th data mode vector from data sample *n* (each of size *N* × *T*).

We show in Supplemental SectionS8.3.3 that reliability is the square of the alignment:

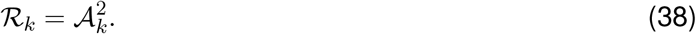

##### Data mode subspace reliability

Because data obtained from finite *N* and *T* may exhibit cross-overlaps, where a single data mode has significant overlap with multiple signal modes, we also define the *subspace reliability* 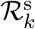, the squared norm of the projection of the *k*th data mode from one data sample onto the full signal subspace of an independent equal-size sample,

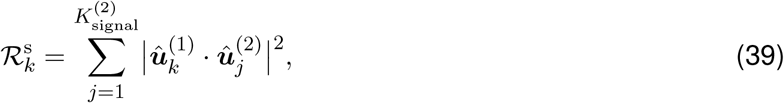

Where 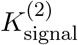 is the number of super-threshold data modes 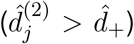 in the second data sample. For a finite number *K* of signal modes, random matrix theory predicts vanishing cross-mode overlap (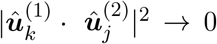 for *j* ≠ *k*) as *N, T* ⟶ ∞, so at leading order the theoretical subspace reliability prediction reduces to the mode reliability in the large data-size limit,

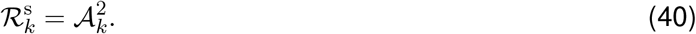

Figure 7A displays the matrix of pairwise squared overlaps, 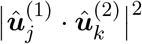 (middle panel), and the subspace reliability 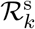 (right panel, diagonal), from a single equal-trial split of the data, as well as the theoretical prediction which is the same for both quantities: 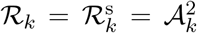 with Equation (36) along the diagonal, and zero elsewhere. Figure 7B shows subspace reliability as a function of data mode strength, calculated for equal-trial splits of the full data. Figure 7C shows the subspace reliability of the top mode from equal-trial splits of size (*N*_full_,*T*) as a function of *T* . Figure 7D shows the subspace reliability of the top mode from equal-trial splits of size 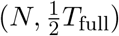 as a function of *N* . Figure 7E shows the top-mode sub-space reliability of all sessions for splits of 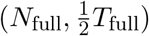. In all panels, the predicted subspace reliability is based on the set of 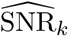 inferred from the full data, (*N*_full_, *T*_full_).

#### Masked Autoencoder prediction via low-rank structure

We assess the predictive capacity of data modes by constructing a masked linear autoencoder (Figure 8A). On a train-set of trials we use the top 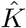 data mode vectors, collected as the columns of 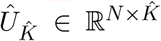, to project to the 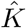 hidden units of the auto-encoder. On a test trial we mask a small number of neurons, project the remaining neural activity onto the hidden units, and then use the rows of 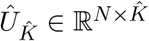 to project back to neural space.

For simplicity, we write the procedure for a single masked neuron, with test-trial activity *r*_0_. Let ***r***_−0_ ∈ ℝ^*N*×1^ be the activity of the remaining population, 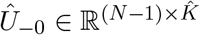 the retained modes with neuron 0’s row masked, and 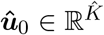 that masked row (ie. neuron 0’s loadings on the retained modes).

The autoencoder’s prediction is therefore

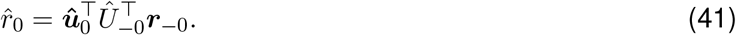

##### Prediction accuracy

We define the predictive accuracy by the fraction of variance explained (coefficient of determination), averaged over held-out test trials and masked neurons:

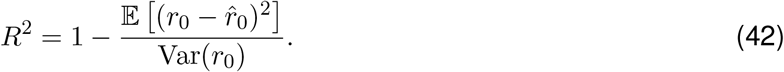

##### Optimal predictive dimensionality

Under the low-rank plus noise model we derive a closed-form expression for the expected *R*^2^ (Supplemental Section S9):

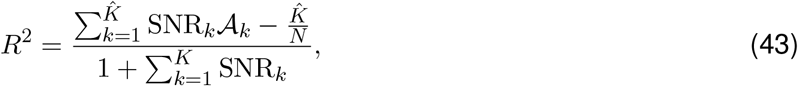

where *A*_*k*_ is the data mode alignment of the *k*th data mode with the corresponding signal mode (Eq. (36)), and *K* is the number of true signal modes.

We show that maximizing *R*^2^ over the number of retained modes, 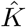, yields an optimal threshold that predicts the predictive dimensionality, 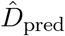: modes should be retained if and only if they have inferred SNR that satisfies 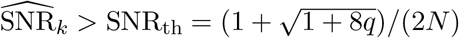, as reported in Equation (10) in the main text.

Here we report an equivalent formulation of this threshold, in terms of the covariance data modes (eigenvalues) themselves. After estimating 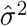 with Equation (27), the number of modes retained is the number of data modes with 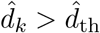, where the threshold is:

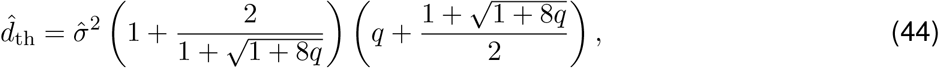

where again, 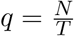.

##### Empirical procedure

On each repetition, we hold out one trial and from the remaining train set we compute *Û* and the inferred 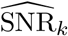 (Section Low-rank Fit). We mask a single neuron at a time, predict each via Eq. (41), and average the resulting *R*^2^ (Eq. (43)) over all masked neurons and test trials; error bars in Figure 8B give the SEM over the product of masked neurons times test trials. Figure 8B compares the maximal *R*^2^ for all sessions from a) the data, scanning the number of retained modes 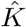 and measuring *R*^2^ for each value, or b) the theory of Eq. (43) evaluated with all the inferred 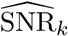 and with 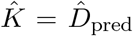 where 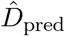 is the number of data modes with 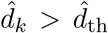 as given in Equation (44); panel C shows *R*^2^ versus 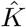 for session #1 for each dataset; panel D compares the empirical optimal number of retained modes to the predicted 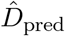. For panel E we recompute the maximal *R*^2^ over a grid of subsampled neuron and trial counts, repeating the entire procedure for each subsample size. Theoretical predictions are obtained from the full set of 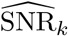 obtained from the full data. Held-out neuron and test-trial counts are listed below. Only the three datasets that admit a low-rank fit are included throughout.

| Dataset | Figure 8B–D |  | Figure 8E |  |
| --- | --- | --- | --- | --- |
|  | Masked neurons | Test trials | Masked neurons | Test trials |
| Mouse MP | 200 | 500 | 96 | 1000 |
| Mouse FOV | 300 | 1000 | 72 | 200 |
| Monkey | 200 | 300 | 48 | 200 |

## S1 Supplemental Figures

**Figure S1:**
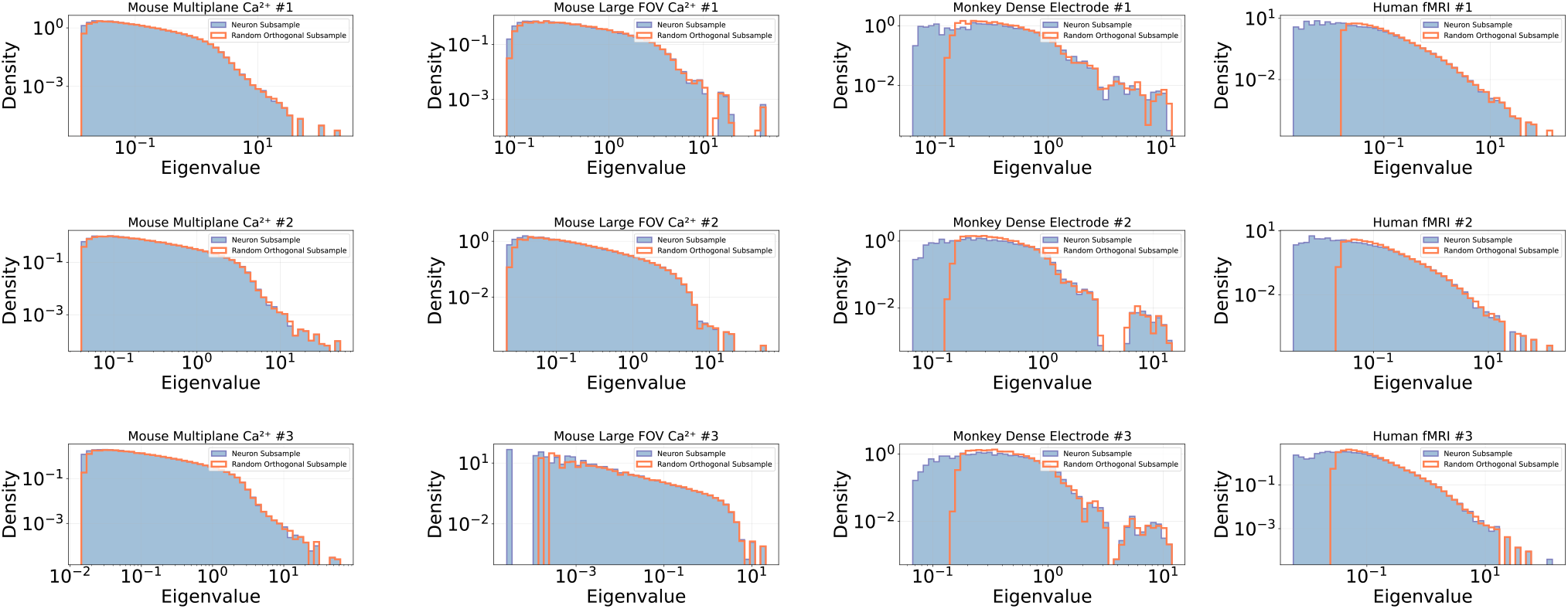
Comparing Eigenvalue Spectra of Neuron-Subsampled Data to Random Projection. Log-log eigenvalue histograms after the full data is reduced to 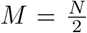 dimensions, either by direct subsampling of *M* neurons (blue histograms), or by random orthogonal projection to *M* dimensions (orange lines). Histograms are mostly indistinguishable except for Monkey and Human data (columns 3 and 4) in which neuron-subsampled data have a noticeable overabundance of small eigenvalues compared to randomly projected data. (Referenced in Section Dimensionality)

**Figure S2:**
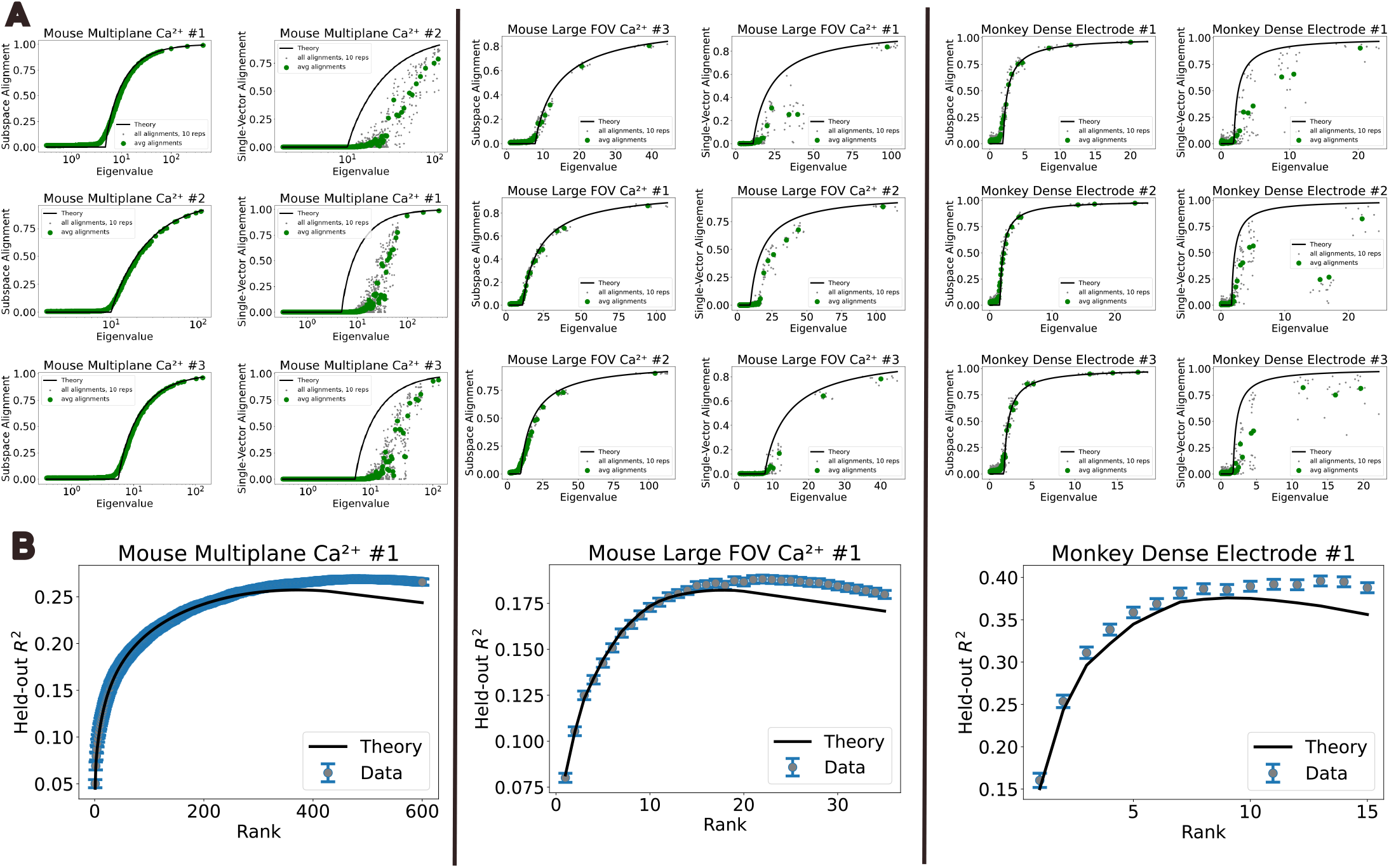
Eigenvector Alignment and Held-Out Neuron Prediction for Low-Rank Toy Model Fit to Data. We generate low-rank toy models from data by inferring SNR for each data mode via Equation (33), keeping those above the critical SNR threshold. We then construct a ground-truth-signal-plus-identity covariance with matching rank and SNRs and a random orthonormal set of eigenvectors, and generate independent Gaussian trial vectors with this covariance. **A**. Subspace alignment (left) and single-vector alignment (right) as a function of eigenvalue for low-rank toy model data. Subspace alignment matches low-rank theory, similarly to real data (Figure 7E). **B**. *R*^2^ in held-out neuron prediction on low-rank toy model data as a function of train rank. Results match low-rank theory similarly to real data (Figure 8C). (Referenced in Discussion)

**Figure S3:**
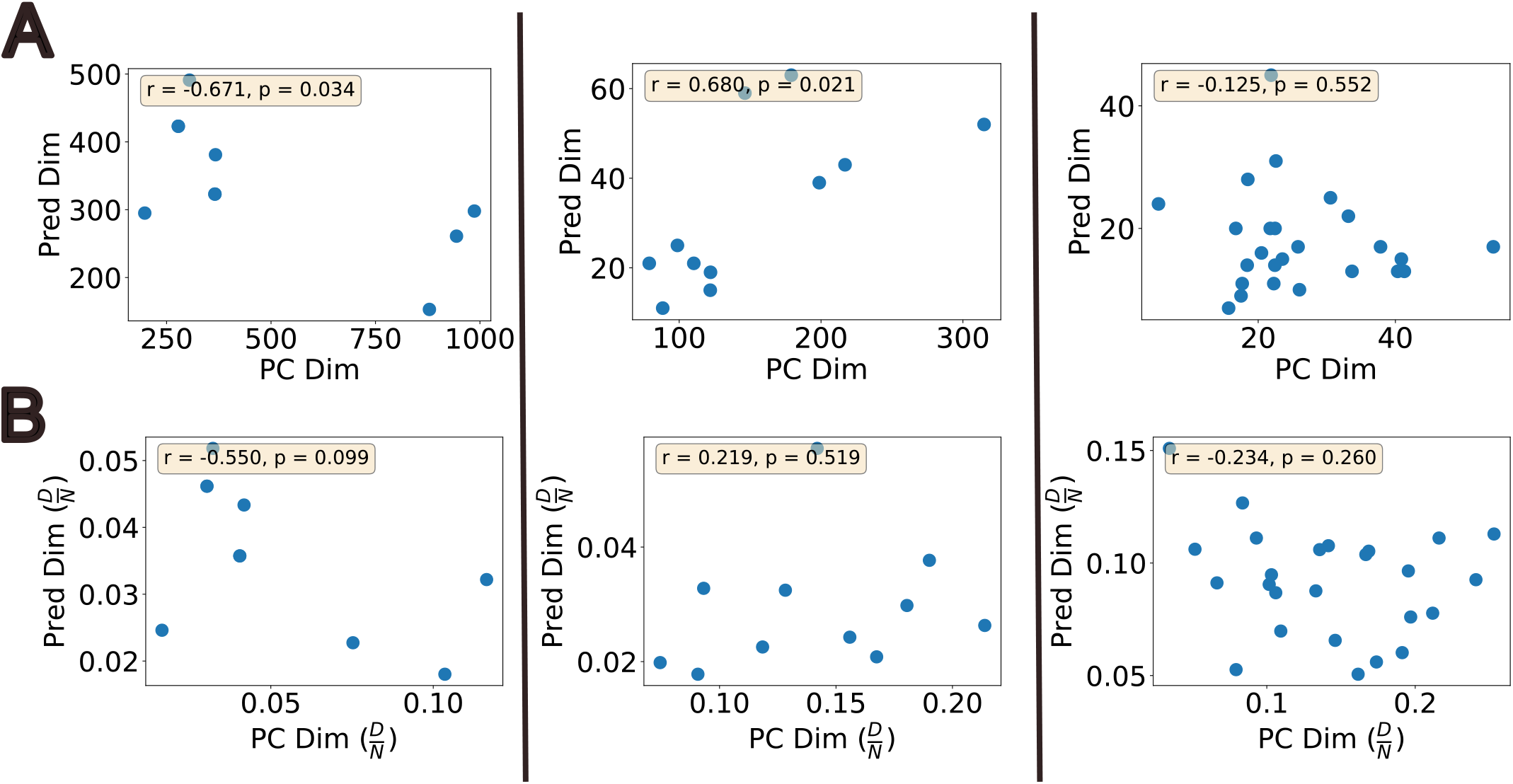
Data Dimensionality Is Not Indicative of Predictive Dimensionality. Scatter plots of predictive dimensionality (optimal rank for masked linear autoencoder) vs data dimensionality. **A**. Unnormalized dimensionalities of all datasets for each data source. (Left) *Mouse V1 multiplane Ca*^*2+*^: predictive and data dimensionalities display a mild negative correlation, (Middle) *Mouse V1 Large FOV Ca*^*2+*^: predictive and data dimensionalities display a positive correlation, (Right) *Monkey dense electrode*: predictive and data dimensionalities are uncorrelated. **B**. The predictive and data dimensionalities normalized by number of neurons, *N*, are uncorrelated across the three datasets. (Referenced in Section Autoencoder)

**Figure S4:**
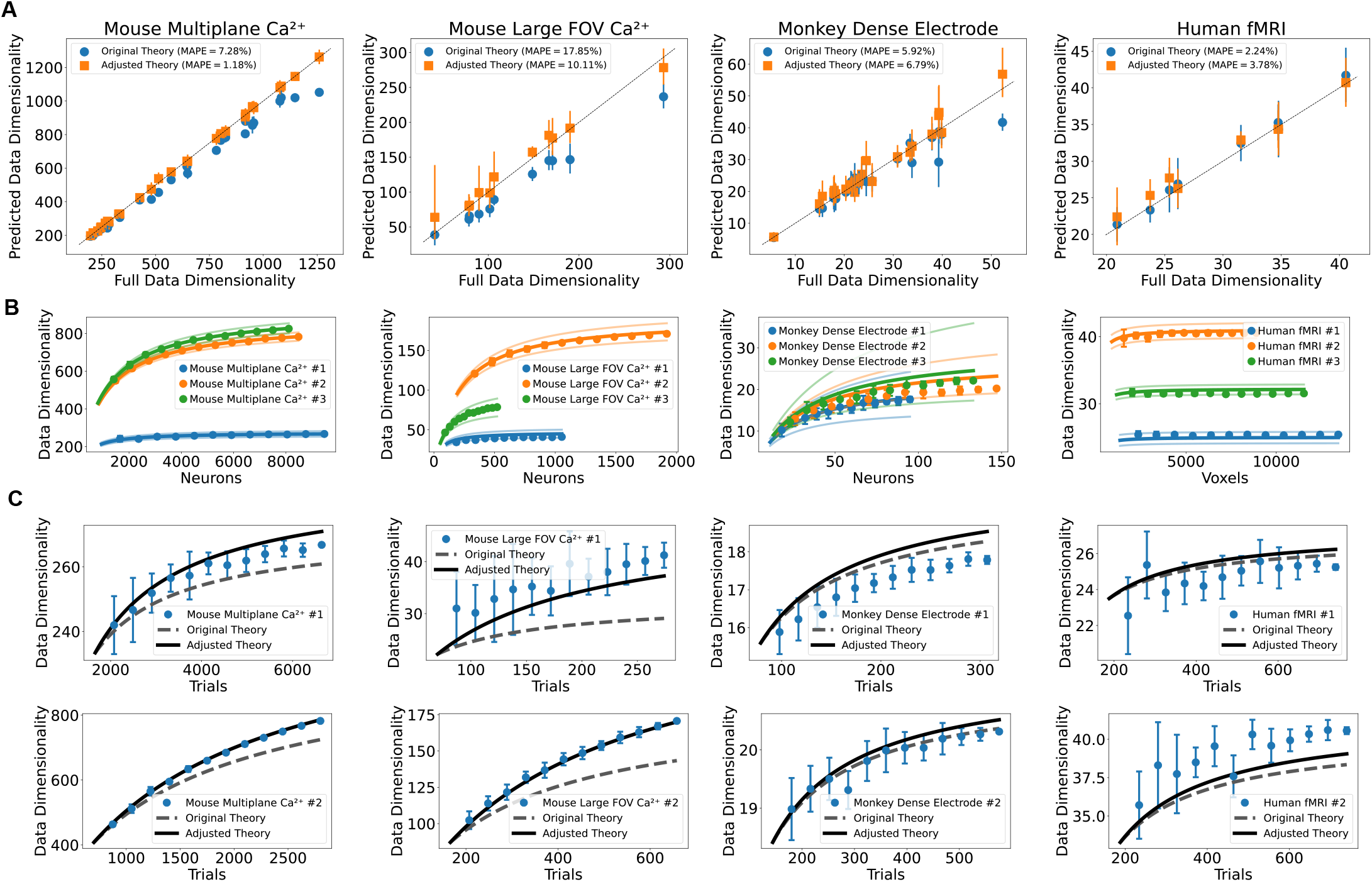
Predicting Data Dimensionality with Trial-Power Adjusted Theory. The theory accounting for variable trial-power effectively extrapolates data dimensionality across datasets without trial-normalizing during preprocessing. See Section S7. **A**. Predicted data dim (extrapolated from 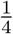 of neurons and trials) vs actual for all datasets of each data source, without trial normalization (Eq. (S7.13)). Blue circles display original theory. Orange squares display adjusted theory. MAPE: Mean absolute percentage error. Standard deviation was computed over 10 repetitions of subsample for computing prediction. **B**. Data dim as a function of neurons/voxels, extrapolated from 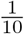 of neurons/voxels. Adjusted and original theory agree as a function of neurons. **C**. Data dim as a function of trials, extrapolated from 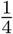 of trials.

**Figure S5:**
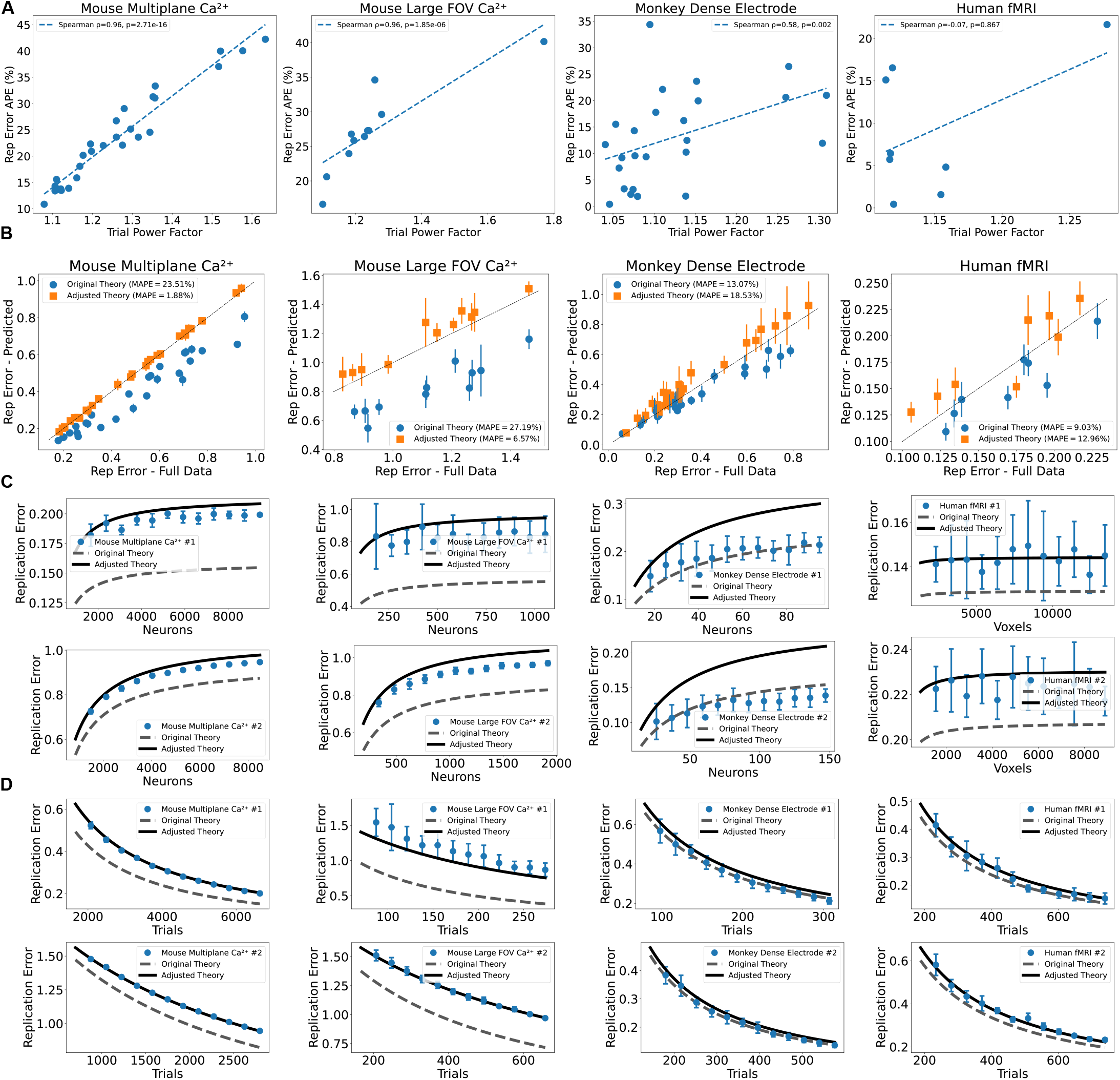
Predicting Replication Error with Trial-Power Adjusted Theory. The theory accounting for variable trial-power effectively predicts replication error across datasets without trial-normalizing during preprocessing. See Section S7. **A**. Absolute percentage error in predicting replication error using the non-adjusted theory vs meansquare trial-power factor, for all datasets of each data source, *without trial normalization*. Datasets with larger trial-to-trial power variability are less well predicted by the original replication error theory (Eq. (S6.19)). **B**. Replication error theory (extrapolated from 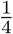 of neurons and trials) vs actual for all datasets of each data source, without trial normalization (Eq. (S7.16)). Blue circles display original theory. Orange squares display adjusted theory. MAPE: Mean absolute percentage error. Standard deviation was computed over 10 repetitions of subsample for computing prediction. **C**. Replication error as a function of neurons/voxels, extrapolated from 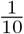 of neurons/voxels. **D**. Replication error as a function of trials, extrapolated from 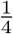 of trials.

**Figure S6:**
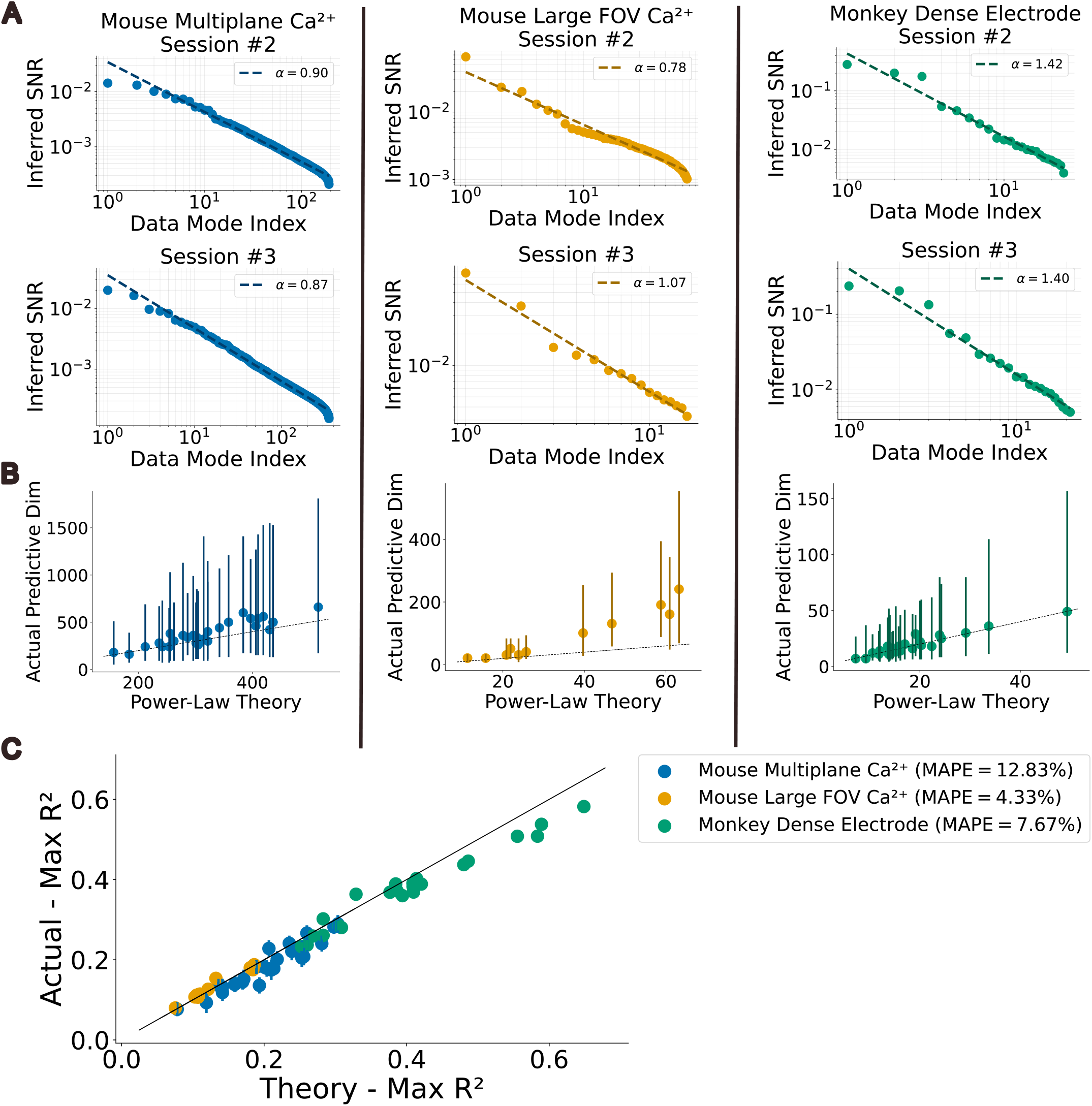
SNR Power-Laws and Predictive Dimensionality. **A**. Log-log scatter of inferred SNRs for all potential signal modes together with linear fit for power-law exponent 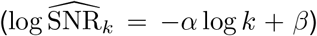, for sessions #2 and #3 of each of the well-fit datasets. See Figure 5C for session #1. **B**. Optimal number of modes, 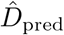, retained for masked autoencoder vs power-law prediction from theory. Prediction is obtained by inserting *α* and SNR_1_ = e^*β*^ From the above fit into Equation (11). **C**. Maximal explained variance obtained with the autoencoder vs power-law theory obtained by estimating SNR_*k*_ = e^*β*^*k*^−*α*^ from the above fit and inserting in Equation (9). See Figure 8 for procedure and to compare theory obtained using individually inferred SNRs.

