## Supplemental Information for "A predictive theory of experimental design for inferring neural population geometry in large-scale recordings"

#### Supplemental Contents

**S1 Supplemental Figures** 2

**S2 Dataset Tables** 8

**S3 Mathematical Supplement – Intro** 10

**S4 Mathematical Preliminaries – Moments of Sample and Population Covariances** 11

**S5 Neural Dimensionality and Data Dimensionality** 12

#### S1 Supplemental Figures

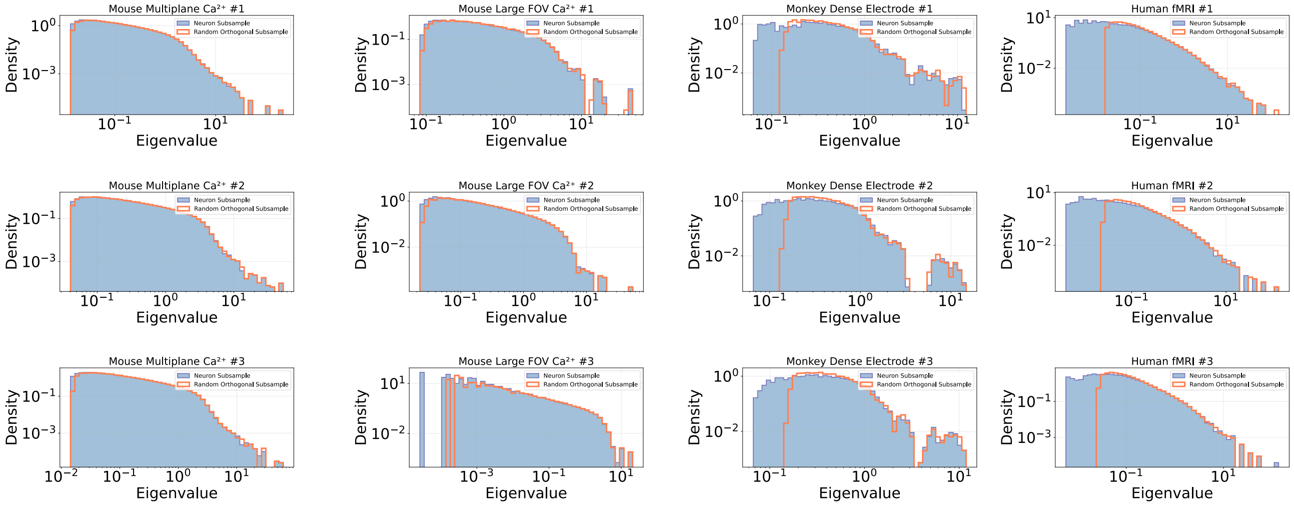

Figure S1: **Comparing Eigenvalue Spectra of Neuron-Subsampled Data to Random Projection.** Log-log eigenvalue histograms after the full data is reduced to $M = \frac{N}{2}$ dimensions, either by direct subsampling of $M$ neurons (blue histograms), or by random orthogonal projection to $M$ dimensions (orange lines). Histograms are mostly indistinguishable except for Monkey and Human data (columns 3 and 4) in which neuron-subsampled data have a noticeable overabundance of small eigenvalues compared to randomly projected data. (Referenced in Section Dimensionality)

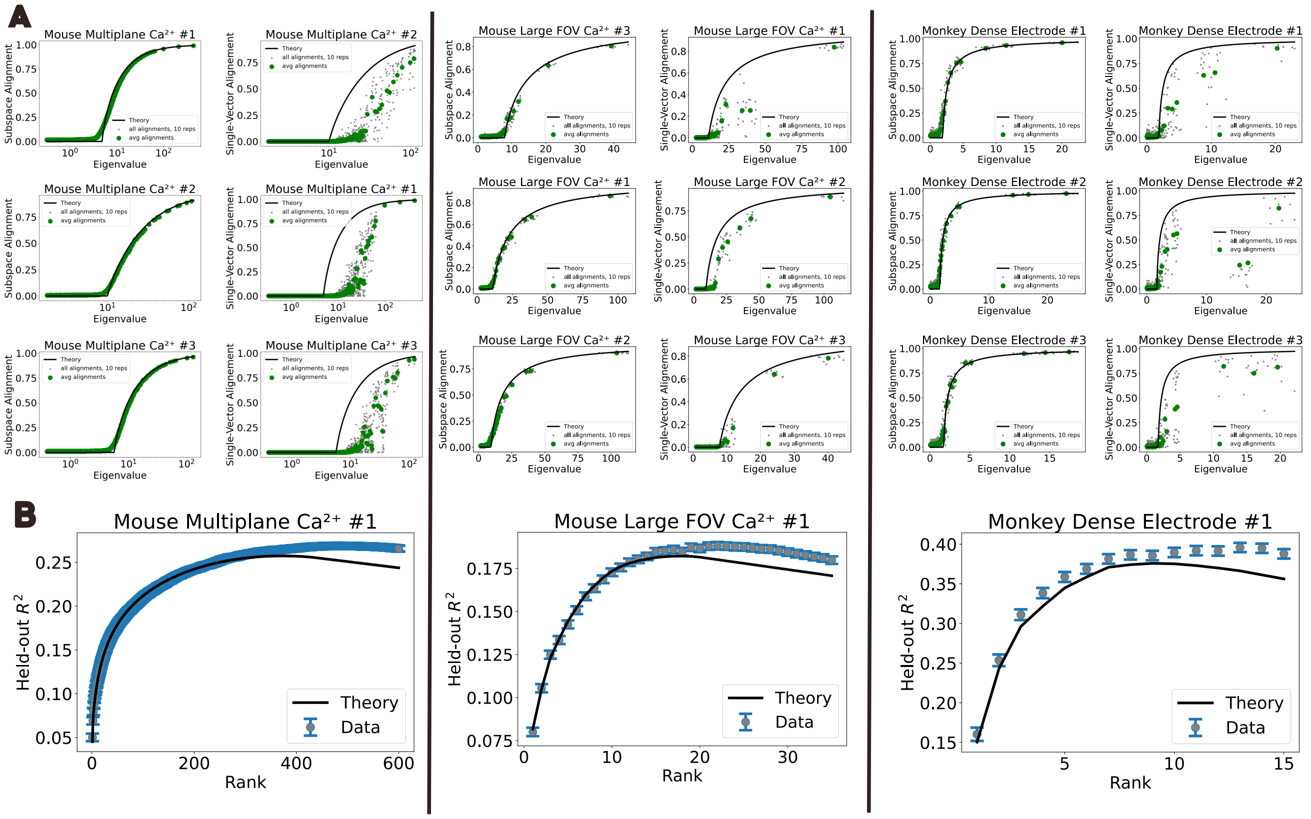

Figure S2: **Eigenvector Alignment and Held-Out Neuron Prediction for Low-Rank Toy Model Fit to Data.** We generate low-rank toy models from data by inferring SNR for each data mode via Equation (33), keeping those above the critical SNR threshold. We then construct a ground-truth-signal-plus-identity covariance with matching rank and SNRs and a random orthonormal set of eigenvectors, and generate independent Gaussian trial vectors with this covariance. **A.** Subspace alignment (left) and single-vector alignment (right) as a function of eigenvalue for low-rank toy model data. Subspace alignment matches low-rank theory, similarly to real data (Figure 7E). **B.** $R^2$ in held-out neuron prediction on low-rank toy model data as a function of train rank. Results match low-rank theory similarly to real data (Figure 8C). (Referenced in Discussion)

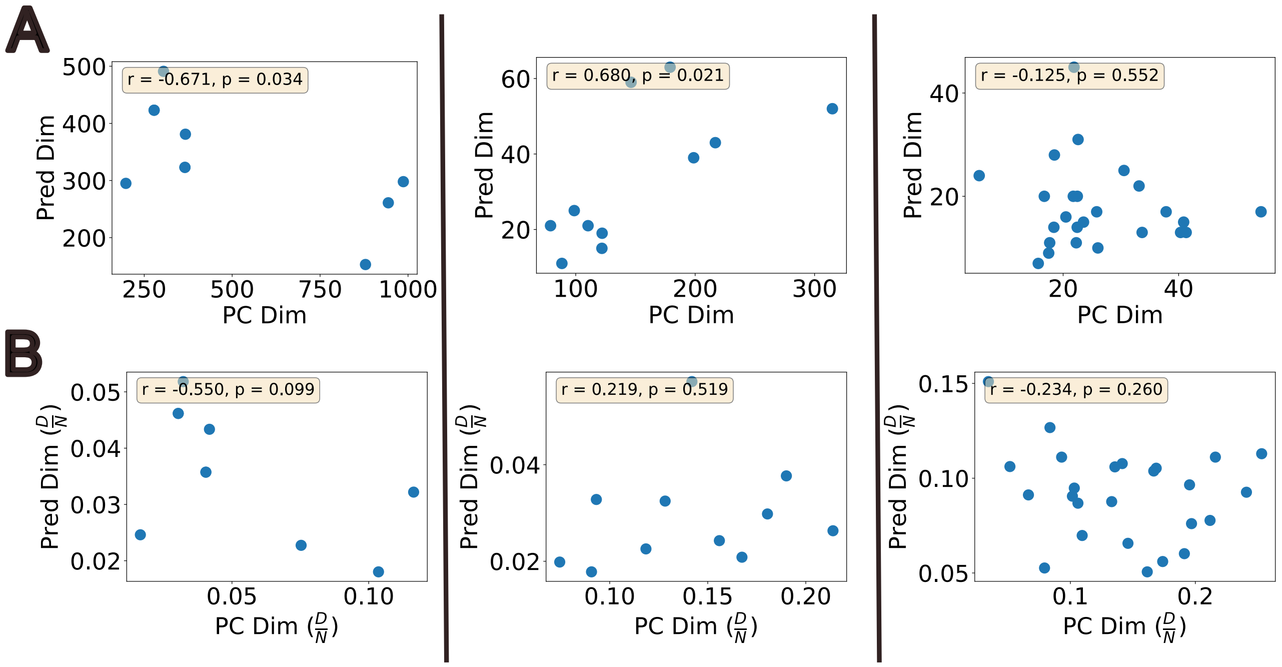

Figure S3: **Data Dimensionality Is Not Indicative of Predictive Dimensionality.** Scatter plots of predictive dimensionality (optimal rank for masked linear autoencoder) vs data dimensionality. **A.** Unnormalized dimensionalities of all datasets for each data source. (Left) *Mouse V1 multiplane* $Ca^{2+}$: predictive and data dimensionalities display a mild negative correlation, (Middle) *Mouse V1 Large FOV* $Ca^{2+}$: predictive and data dimensionalities display a positive correlation, (Right) *Monkey dense electrode*: predictive and data dimensionalities are uncorrelated. **B.** The predictive and data dimensionalities normalized by number of neurons, $N$, are uncorrelated across the three datasets. (Referenced in Section Autoencoder)

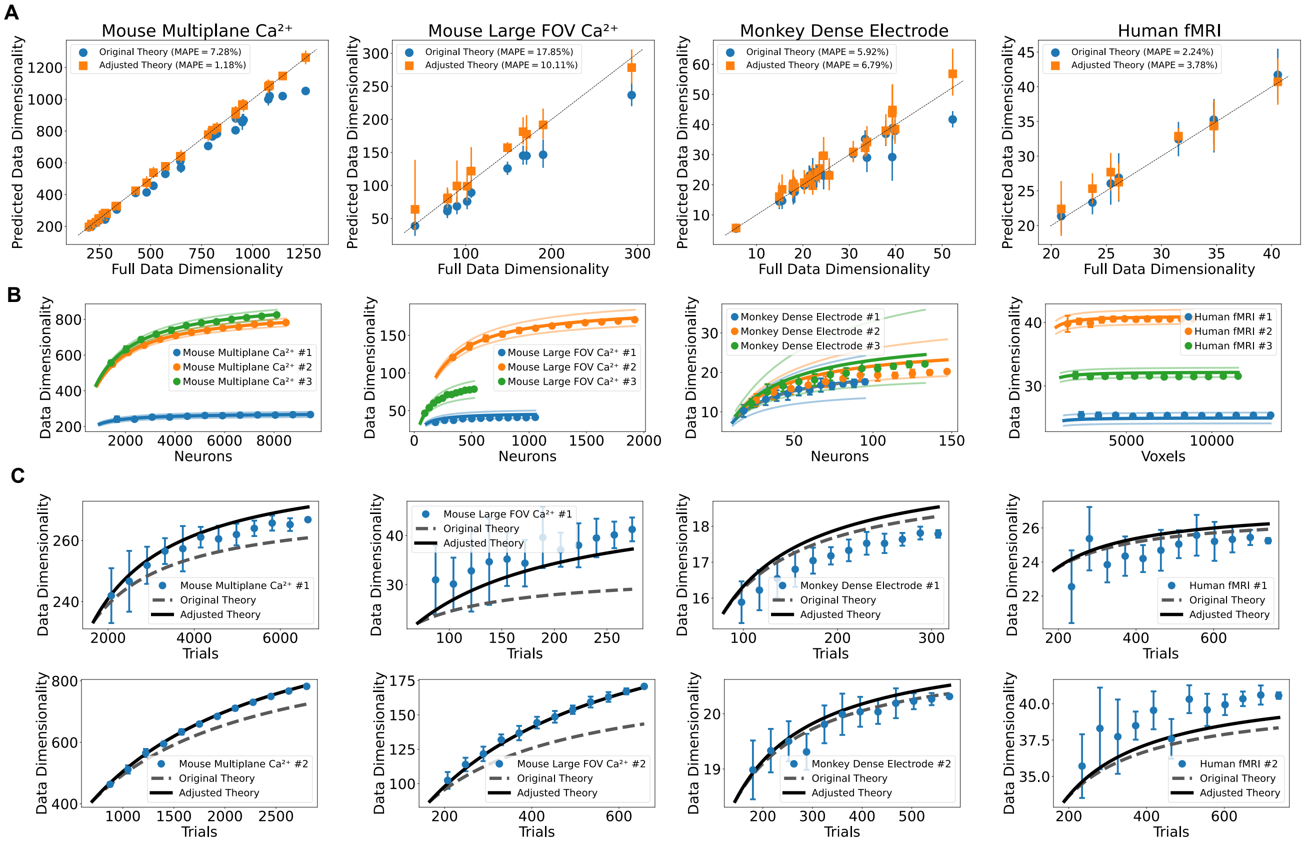

Figure S4: **Predicting Data Dimensionality with Trial-Power Adjusted Theory.** The theory accounting for variable trial-power effectively extrapolates data dimensionality across datasets without trial-normalizing during preprocessing. See Section S7. **A.** Predicted data dim (extrapolated from $\frac{1}{4}$ of neurons and trials) vs actual for all datasets of each data source, without trial normalization (Eq. (S7.13)). Blue circles display original theory. Orange squares display adjusted theory. MAPE: Mean absolute percentage error. Standard deviation was computed over 10 repetitions of subsample for computing prediction. **B.** Data dim as a function of neurons/voxels, extrapolated from $\frac{1}{10}$ of neurons/voxels. Adjusted and original theory agree as a function of neurons. **C.** Data dim as a function of trials, extrapolated from $\frac{1}{4}$ of trials.

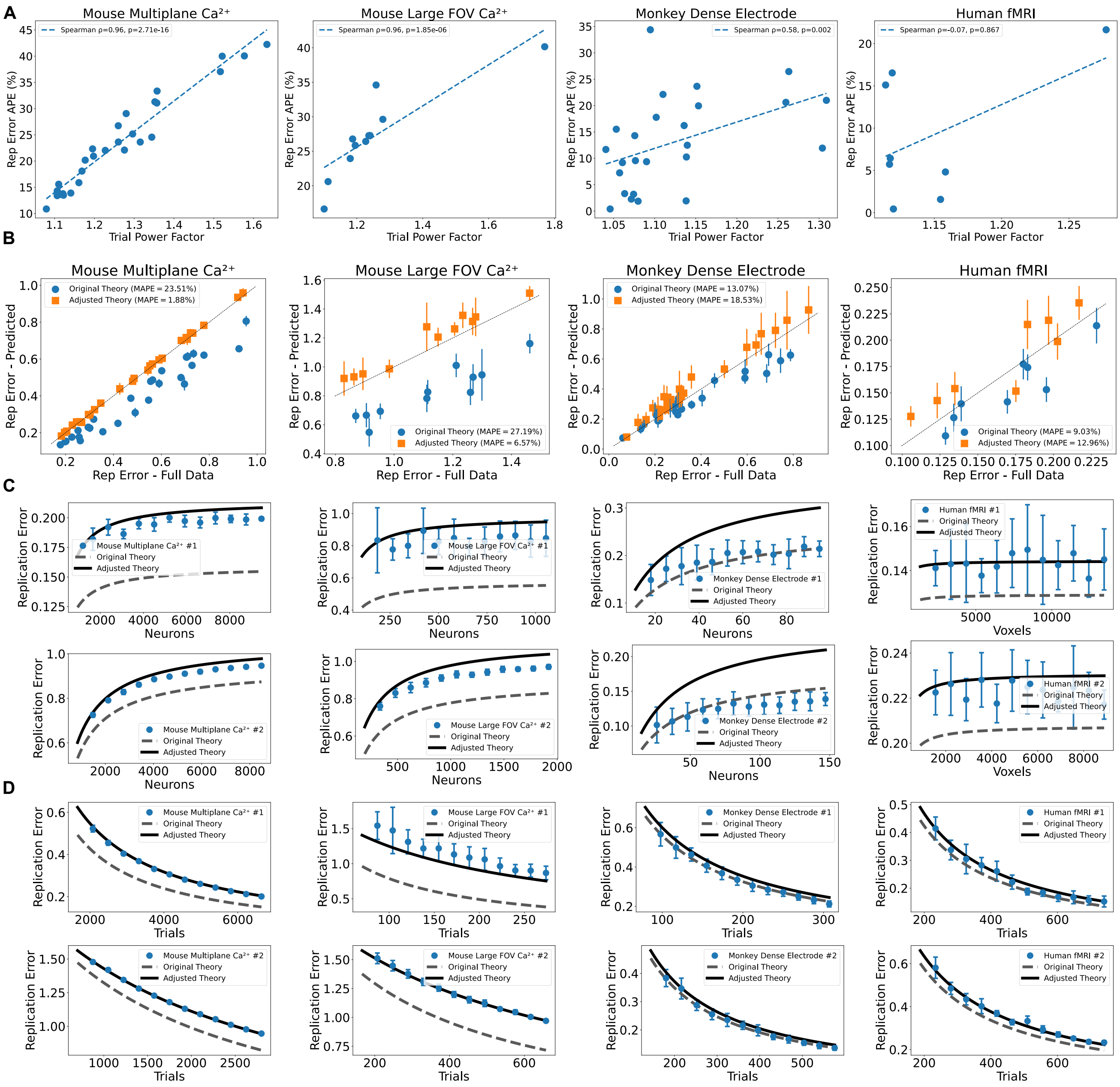

Figure S5: **Predicting Replication Error with Trial-Power Adjusted Theory.** The theory accounting for variable trial-power effectively predicts replication error across datasets without trial-normalizing during preprocessing. See Section S7. **A.** Absolute percentage error in predicting replication error using the non-adjusted theory vs mean-square trial-power factor, for all datasets of each data source, *without trial normalization*. Datasets with larger trial-to-trial power variability are less well predicted by the original replication error theory (Eq. (S6.19)). **B.** Replication error theory (extrapolated from $\frac{1}{4}$ of neurons and trials) vs actual for all datasets of each data source, without trial normalization (Eq. (S7.16)). Blue circles display original theory. Orange squares display adjusted theory. MAPE: Mean absolute percentage error. Standard deviation was computed over 10 repetitions of subsample for computing prediction. **C.** Replication error as a function of neurons/voxels, extrapolated from $\frac{1}{10}$ of neurons/voxels. **D.** Replication error as a function of trials, extrapolated from $\frac{1}{4}$ of trials.

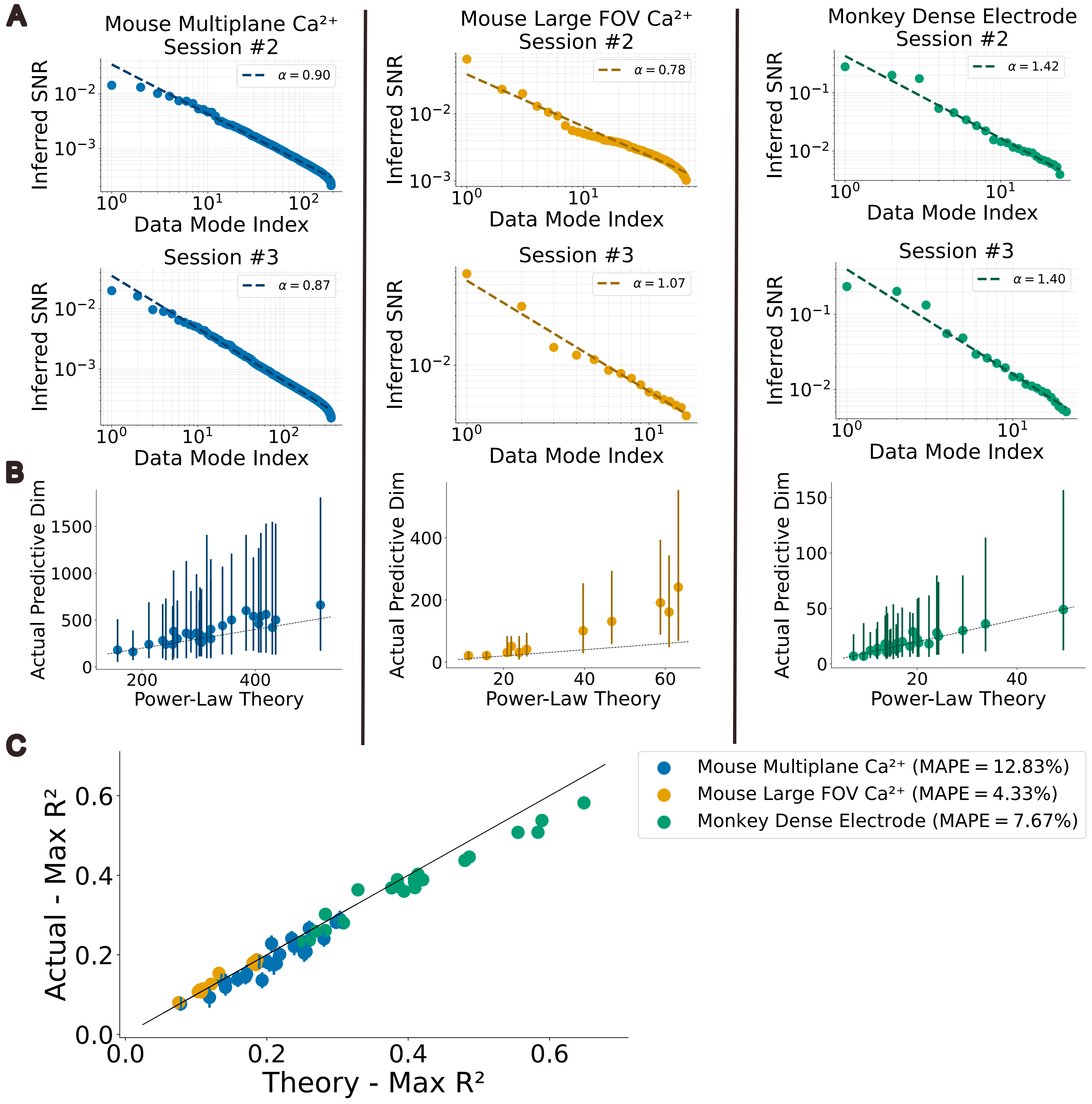

Figure S6: **SNR Power-Laws and Predictive Dimensionality A.** Log-log scatter of inferred SNRs for all potential signal modes together with linear fit for power-law exponent ($\log \widehat{\mathrm{SNR}}_k = -\alpha \log k + \beta$), for sessions #2 and #3 of each of the well-fit datasets. See Figure 5C for session #1. **B.** Optimal number of modes, $\hat{D}_{\mathrm{pred}}$, retained for masked autoencoder vs power-law prediction from theory. Prediction is obtained by inserting $\alpha$ and $\mathrm{SNR}_1 = \mathrm{e}^{\beta}$ from the above fit into Equation (11). **C.** Maximal explained variance obtained with the autoencoder vs power-law theory obtained by estimating $\mathrm{SNR}_k = \mathrm{e}^{\beta} k^{-\alpha}$ from the above fit and inserting in Equation (9). See Figure 8 for procedure and to compare theory obtained using individually inferred SNRs.

#### S2 Dataset Tables

For each of the four datasets, we report the number of neurons (or voxels) $N$, the number of trials (or images) $T$, the data dimensionality $\hat{D}$ of the data covariance, the total number of potential signal modes under the low-rank plus noise model, $\hat{D}_{\mathrm{signal}}$, the optimal predictive dimensionality $D_{\mathrm{pred}}$, and the inferred signal-to-noise ratio $\mathrm{SNR}_1$ and $\mathrm{SNR}_2$ of the top two data modes. Sessions marked "idx #$k$" are displayed in individual-session figures in the main text.

Table 1: ***Mouse V1 multiplane Ca$^{2+}$*** **(Stringer et al., 2019).** Recording sessions from multiple mice with various image stimulus sets. If not otherwise stated the stimulus set consisted of 2800 natural images, typically two repeats. In sessions labeled '4D' or '8D', the same 2800 images were projected to 4 / 8 principal dimensions of neural response. In sessions labeled 'whitened', the images were spatially whitened. In sessions labeled 'small', images were zeroed outside the receptive field location of the recorded cells. In sessions labeled '32 images', 32 natural images were selected and repeated 90–114 times. Data available at `https://github.com/MouseLand/stringer-pachitariu-et-al-2018b`.

| Session Date | Mouse | Stimuli | $N$ | $T$ | $\hat{D}$ | $\hat{D}_{\mathrm{signal}}$ | $\hat{D}_{\mathrm{pred}}$ | $\mathrm{SNR}_1$ | $\mathrm{SNR}_2$ |
|---|---|---|---|---|---|---|---|---|---|
| 2017-09-14 (idx #1) | MP032 | | 9475 | 6645 | 305 | 597 | 491 | 0.07 | 0.03 |
| 2017-09-19 (idx #2) | MP033 | 4D | 8487 | 2800 | 879 | 192 | 153 | 0.01 | 0.01 |
| 2017-09-22 (idx #3) | MP033 | 4D | 8108 | 5600 | 944 | 353 | 261 | 0.02 | 0.02 |
| 2017-08-07 | MP032 | | 9039 | 5600 | 366 | 411 | 323 | 0.06 | 0.03 |
| 2017-09-15 | MP032 | 8D | 9163 | 5600 | 278 | 510 | 423 | 0.07 | 0.04 |
| 2017-09-18 | MP032 | small | 9136 | 5600 | 334 | 454 | 371 | 0.07 | 0.04 |
| 2017-09-11 | MP032 | whitened | 7865 | 5600 | 240 | 458 | 366 | 0.06 | 0.03 |
| 2017-09-22 | MP032 | 4D | 8789 | 5600 | 367 | 464 | 381 | 0.05 | 0.04 |
| 2017-06-27 | MP031 | 32 images | 11987 | 3200 | 198 | 340 | 295 | 0.09 | 0.04 |
| 2017-06-07 | MP030 | 8D | 13106 | 5600 | 987 | 377 | 298 | 0.02 | 0.01 |
| 2017-09-20 | MP034 | 4D | 13332 | 5600 | 783 | 370 | 292 | 0.03 | 0.02 |
| 2017-07-02 | MP031 | 8D | 9145 | 5600 | 409 | 440 | 354 | 0.05 | 0.03 |
| 2017-08-10 | MP032 | 8D | 9933 | 5600 | 319 | 428 | 343 | 0.06 | 0.04 |
| 2017-08-22 | MP033 | 8D | 10130 | 5600 | 1135 | 326 | 246 | 0.01 | 0.01 |
| 2017-09-15 | MP034 | 8D | 12823 | 5600 | 656 | 493 | 409 | 0.02 | 0.02 |
| 2016-12-14 | MP027 | | 11445 | 5166 | 1067 | 308 | 235 | 0.02 | 0.01 |
| 2017-05-29 | MP030 | | 14062 | 5390 | 1320 | 350 | 277 | 0.02 | 0.01 |
| 2017-06-28 | MP031 | | 10075 | 5600 | 609 | 385 | 301 | 0.04 | 0.01 |
| 2017-08-20 | MP033 | | 10144 | 6621 | 956 | 505 | 397 | 0.03 | 0.01 |
| 2017-09-11 | MP034 | | 10102 | 5600 | 818 | 375 | 291 | 0.03 | 0.01 |
| 2017-08-23 | MP033 | small | 11627 | 5600 | 687 | 282 | 192 | 0.04 | 0.00 |
| 2017-09-17 | MP034 | small | 13622 | 5600 | 1102 | 458 | 378 | 0.02 | 0.01 |
| 2017-09-12 | MP032 | whitened | 8246 | 5600 | 307 | 491 | 402 | 0.06 | 0.04 |
| 2017-09-21 | MP033 | whitened | 8961 | 5600 | 1231 | 356 | 266 | 0.02 | 0.01 |
| 2017-09-14 | MP034 | whitened | 9478 | 3845 | 279 | 404 | 348 | 0.07 | 0.03 |
| 2016-03-23 | MP019 | 32 images | 11652 | 2832 | 533 | 216 | 184 | 0.04 | 0.02 |
| 2017-08-01 | MP032 | 32 images | 12578 | 3648 | 392 | 326 | 272 | 0.05 | 0.03 |
| 2017-08-25 | MP033 | 32 images | 11523 | 3072 | 518 | 289 | 243 | 0.04 | 0.01 |

Table 2: ***Mouse V1 Large FOV Ca$^{2+}$*** **(Rumyantsev et al., 2020).** Eleven mice, each viewing one of two pairs of oriented visual gratings ($\pm 6°$ or $\pm 30°$ angular separation). $T$ pools both stimulus orientations. Data available from the original publication.

| Mouse | Stimuli | $N$ | $T$ | $\hat{D}$ | $\hat{D}_{\text{signal}}$ | $\hat{D}_{\text{pred}}$ | $\text{SNR}_1$ | $\text{SNR}_2$ |
|---|---|---|---|---|---|---|---|---|
| Mouse 15 (idx #1) | $\pm 6°$ | 1059 | 280 | 79 | 25 | 21 | 0.10 | 0.04 |
| Mouse L347 (idx #2) | $\pm 30°$ | 1921 | 662 | 179 | 76 | 63 | 0.07 | 0.02 |
| Mouse 13 (idx #3) | $\pm 6°$ | 528 | 257 | 88 | 16 | 11 | 0.08 | 0.04 |
| Mouse 10 | $\pm 6°$ | 931 | 247 | 110 | 24 | 21 | 0.06 | 0.02 |
| Mouse 11 | $\pm 6°$ | 570 | 333 | 122 | 17 | 15 | 0.05 | 0.03 |
| Mouse 12 | $\pm 6°$ | 783 | 281 | 122 | 22 | 19 | 0.05 | 0.03 |
| Mouse 14 | $\pm 6°$ | 770 | 331 | 99 | 30 | 25 | 0.08 | 0.04 |
| Mouse L354 | $\pm 30°$ | 1141 | 484 | 217 | 52 | 43 | 0.03 | 0.02 |
| Mouse L355 | $\pm 30°$ | 2191 | 435 | 199 | 48 | 39 | 0.04 | 0.02 |
| Mouse L362 | $\pm 30°$ | 1031 | 641 | 146 | 70 | 59 | 0.07 | 0.04 |
| Mouse L363 | $\pm 30°$ | 1745 | 566 | 315 | 63 | 52 | 0.02 | 0.01 |

Table 3: ***Monkey motor and pre-motor dense electrode*** **(Sun, O'Shea et al., 2022; O'Shea, Duncker et al., 2022; Vyas et al., 2022).** Twenty-four sessions from two rhesus macaques (P and V) performing a center-out reach task. Brain region per session is indicated where recorded separately. $T$ pools all reach directions. Data available from the corresponding authors upon reasonable request; not publicly archived.

| Session ID | Monkey | Brain Region | $N$ | $T$ | $\hat{D}$ | $\hat{D}_{\text{signal}}$ | $\hat{D}_{\text{pred}}$ | $\text{SNR}_1$ | $\text{SNR}_2$ |
|---|---|---|---|---|---|---|---|---|---|
| V20180822 (idx #1) | V | M1 | 106 | 316 | 18 | 16 | 11 | 0.33 | 0.20 |
| V20180919 (idx #2) | V | PMd | 151 | 578 | 20 | 24 | 16 | 0.28 | 0.20 |
| V20180820 (idx #3) | V | PMd | 133 | 552 | 22 | 21 | 14 | 0.24 | 0.20 |
| P20180323 | P | M1 | 108 | 248 | 26 | 12 | 10 | 0.18 | 0.15 |
| P20180327 | P | M1 | 62 | 253 | 16 | 10 | 7 | 0.28 | 0.18 |
| P20180607 | P | PMd | 336 | 253 | 54 | 22 | 17 | 0.12 | 0.07 |
| P20180608 | P | PMd | 259 | 258 | 38 | 21 | 17 | 0.17 | 0.12 |
| P20180609 | P | PMd | 193 | 256 | 41 | 17 | 15 | 0.15 | 0.08 |
| P20180612 | P | PMd | 418 | 258 | 33 | 26 | 22 | 0.21 | 0.12 |
| P20180613 | P | PMd | 171 | 204 | 34 | 17 | 13 | 0.16 | 0.12 |
| P20180614 | P | PMd | 216 | 208 | 41 | 15 | 13 | 0.15 | 0.08 |
| P20180615 | P | PMd | 232 | 207 | 40 | 16 | 13 | 0.14 | 0.09 |
| P20180620 | P | M1 | 130 | 249 | 18 | 17 | 14 | 0.32 | 0.26 |
| P20180622 | P | M1 | 180 | 207 | 17 | 24 | 20 | 0.49 | 0.30 |
| P20180704 | P | M1 | 424 | 364 | 22 | 51 | 45 | 0.42 | 0.38 |
| P20180705 | P | M1 | 221 | 335 | 18 | 33 | 28 | 0.42 | 0.36 |
| P20180707 | P | M1 | 340 | 281 | 23 | 35 | 31 | 0.36 | 0.24 |
| P20180711 | P | PMd | 81 | 398 | 17 | 15 | 9 | 0.27 | 0.23 |
| V20180814 | V | PMd | 221 | 287 | 22 | 25 | 20 | 0.31 | 0.22 |
| V20180815 | V | PMd | 194 | 313 | 26 | 22 | 17 | 0.24 | 0.22 |
| V20180817 | V | PMd | 215 | 267 | 24 | 20 | 15 | 0.29 | 0.16 |
| V20180818 | V | PMd | 288 | 563 | 31 | 33 | 25 | 0.23 | 0.16 |
| V20180819 | V | PMd | 114 | 320 | 22 | 16 | 11 | 0.22 | 0.21 |
| V20180821 | V | M1 | 211 | 301 | 22 | 24 | 20 | 0.30 | 0.24 |

Table 4: ***Human fMRI* (Allen et al., 2022).** Eight subjects from the Natural Scenes Dataset (NSD), left hemisphere, voxelwise responses to 750 natural images pooled across visual ROIs (V1v, V1d, V2v, V2d, V3v, V3d, hV4). Data available at `https://naturalscenesdataset.org`. Note: this dataset constitutes a known failure case for the low-rank plus noise model; see Results and Discussion.

| Subject | $N$ | $T$ | $\hat{D}$ |
|---|---|---|---|
| 1 (idx #1) | 13440 | 750 | 35 |
| 4 (idx #2) | 8893 | 750 | 49 |
| 6 (idx #3) | 11546 | 750 | 39 |
| 2 | 11361 | 750 | 44 |
| 3 | 11162 | 750 | 33 |
| 5 | 11097 | 750 | 29 |
| 7 | 9582 | 750 | 34 |
| 8 | 9631 | 750 | 43 |

### S3 Mathematical Supplement – Intro

In this Supplemental Note we provide derivations for the theoretical results in the main manuscript. We present our general assumptions here. In Section S4 we review mathematical preliminaries. In Section S5 we present the derivation of the scaling of data dimensionality with neurons and trials (main text Section Dimensionality). In Section S6 we derive the scaling of accuracy and reliability of the sample covariance with neurons and trials (main text Section Accuracy and Reliability). In Section S8 we review existing random matrix results for the low-rank plus noise model, present our method for fitting the model to data, and derive the implications for signal geometry as a function of neurons and trials (main text Section Low-rank plus noise model). In Section S9 we derive the predictive variance of a masked linear autoencoder and the optimal predictive dimensionality, a principled threshold for dimensionality reduction obtained by optimizing the number of hidden units in the autoencoder (main text Section Held-out Prediction). Section S10 reviews and derives random matrix formulas used throughout the supplement.

Our general model for neural data is an $N \times T$ data matrix, $R_{N,T}$, consisting of $T$ random, independent samples from an $N$-dimensional unknown *neural covariance*:

$$R_{N,T} = \sqrt{C_N} X_{N,T}, \tag{S3.1}$$

with $C_N$ a positive-definite $N \times N$ covariance matrix with arbitrary spectrum, and $X_{N,T}$ a random, i.i.d. $N \times T$ matrix with zero mean, unit variance, and finite $4$th moment.

We write the neurons-to-trials aspect ratio as

$$q := \frac{N}{T}. \tag{S3.2}$$

We write the *data covariance* (sample covariance) and its eigen-decomposition as

$$\hat{C}_{N,T} := \frac{1}{T} R_{N,T} R_{N,T}^T = \sum_{k=1}^{N} \hat{d}_k \hat{\boldsymbol{u}}_k \hat{\boldsymbol{u}}_k^T. \tag{S3.3}$$

Similarly, the neural covariance is assumed to have eigen-decomposition $C_N = \sum_k d_k \boldsymbol{u}_k \boldsymbol{u}_k^T$. We refer to the pair $(d_k, \boldsymbol{u}_k)$ as the $k$th *neural mode*, and $(\hat{d}_k, \hat{\boldsymbol{u}}_k)$ as the corresponding *data mode.*

Note that we can write

$$\hat{C}_{N,T} = \sqrt{C_N} W_{N,T} \sqrt{C_N}, \tag{S3.4}$$

where $W_{N,T} = \frac{1}{T} X_{N,T} X_{N,T}^T$ is a covariance of i.i.d. noise with aspect ratio $q$.

#### S4 Mathematical Preliminaries – Moments of Sample and Population Covariances

We begin with a few standard results that we will use throughout the supplement. Define the normalized trace operator:

$$\tau[A] := \frac{1}{N}\sum_{i=1}^{N} A_{ii}, \tag{S4.1}$$

where $A$ is an $N \times N$ matrix.

First, we have

$$\tau[W_{N,T}] = \frac{1}{NT}\sum_{i,t} X_{it}^2 \to 1, \tag{S4.2}$$

where the second step follows from the law of large numbers.

Next, we will need

$$\tau[W_{N,T}^2] = \frac{1}{NT^2}\sum_{i,j,s,t} X_{it}X_{jt}X_{js}X_{is}. \tag{S4.3}$$

The four-way product will average out to zero over all combinations of indices except those in which each element is paired, i.e. either $i = j$ or $s = t$. Those terms yield, in the large $T$ limit:

$$\tau[W_{N,T}^2] = \frac{1}{NT^2}\left[\sum_{i=j,s,t} X_{it}^2X_{is}^2 + \sum_{i,j,s=t} X_{it}^2X_{jt}^2 - \sum_{i=j,s=t} X_{it}^4\right] = 1 + \frac{N}{T} + \mathcal{O}\left(\frac{1}{T}\right), \tag{S4.4}$$

where the $\mathcal{O}\left(\frac{1}{T}\right)$ correction is due to the contribution of $NT$ terms with $i = j, s = t$, which depends on the kurtosis of $X$, and which vanishes in the large $N$ limit. This term is the leading contribution to finite $N$ and $T$ errors in our formulas for dimensionality, accuracy, and reliability below.

Next, we will need several mixed trace products - for example, $\tau[C_N W_{N,T}]$. For such products we rely on *free probability* theory, an analogue of probability theory for non-commutative random variables. For random matrices, the (expected) normalized trace plays the role of the expectation operator, and statistical independence is replaced by the notion of *freeness*. While finite matrices generally cannot be exactly free, freeness holds asymptotically (in the limit of large matrix dimension) in a wide variety of matrix settings, including a) the very important case of relatively rotationally invariant matrix ensembles, where $p(A, B) = p(A, OBO^T)$ for arbitrary orthogonal matrix $O$, and b) between fixed deterministic matrices and random matrices consisting of i.i.d. mean-$0$ elements with bounded moments - eg. $C_N$ and $W_{N,T}$. Thus we will assume (asymptotic) freeness of $C_N$ and $W_{N,T}$ throughout.

In classical probability, the expectation of the product of two independent random variables is the product of their expectations. An analogous formula holds for free random variables: $\tau[AB] = \tau[A]\tau[B]$. Thus in particular,

$$\tau[C_N W_{N,T}] = \tau[C_N]\tau[W_{N,T}] = \tau[C_N]. \tag{S4.5}$$

Free probability theory also yields formulas for mixed products of higher order, which never arise in classical probability where multiplication of random variables is commutative. In particular, we will need to consider the following mixed-product trace: $\tau[ABAB]$.

In free probability, the trace of all mixed-products of *trace-zero* variables is zero. Thus in principle, one can center each variable and write

$$0 = \tau\left[(A - \tau[A])(B - \tau[B])(A - \tau[A])(B - \tau[B])\right]. \tag{S4.6}$$

Then in order to calculate $\tau[ABAB]$ one can expand the above as a non-commutative binomial expansion into 16 terms. This is straightforward but somewhat cumbersome, and becomes quickly unwieldy for higher mixed moments.

To understand how such moments are computed efficiently in free probability, it is helpful to use the following analogy to ordinary probability. In the classical setting, where multiplication of random variables is commutative, the moment $\mathbb{E}[abab] = \mathbb{E}[a^2b^2]$ for independent random variables $a$ and $b$ can be expressed in terms of a cumulant expansion, which sums over all ways of partitioning the factors $\{a, a, b, b\}$ into pure $a$- and $b$-subsets. For each such partition, each subset contributes a corresponding cumulant factor. Thus, schematically,

$$\mathbb{E}[abab] = \{a,a\}\{b,b\} + \{a\}\{a\}\{b,b\} + \{a,a\}\{b\}\{b\} + \{a\}\{a\}\{b\}\{b\} \tag{S4.7}$$

$$= \kappa_2(a)\,\kappa_2(b) + \kappa_1(a)^2\,\kappa_2(b) + \kappa_2(a)\,\kappa_1(b)^2 + \kappa_1(a)^2\,\kappa_1(b)^2\,, \tag{S4.8}$$

where $\kappa_1(x) = \mathbb{E}[x]$ and $\kappa_2(x) = \mathbb{E}[x^2] - \mathbb{E}[x]^2$ are the first two classical cumulants. Free probability provides an analogous expansion in terms of *free* cumulants, where the sum over arbitrary set partitions is replaced by one over *non-crossing* partitions of the letters of the word $ABAB$. Non-crossing means that we consider the four letters *in order* and any partition with intersecting subsets is excluded. For example, the partition $\{A, A\}\{B, B\}$ is excluded because the two subsets intersect, whereas the partition $\{A, A\}\{B\}\{B\}$ is included because singleton subsets do not intersect. Thus, we are left with

$$\tau[ABAB] = \{A\}\{A\}\{B,B\} + \{A,A\}\{B\}\{B\} + \{A\}\{A\}\{B\}\{B\} \tag{S4.9}$$

$$= \kappa_1(A)^2\,\kappa_2(B) + \kappa_2(A)\,\kappa_1(B)^2 + \kappa_1(A)^2\,\kappa_1(B)^2\,, \tag{S4.10}$$

where $\kappa_1(X) = \tau[X]$ and $\kappa_2(X) = \tau[X^2] - \tau[X]^2$ are the first two *free cumulants*.

Thus, we find

$$\tau[ABAB] = \tau[A]^2\tau[B^2] + \tau[B]^2\tau[A^2] - \tau[A]^2\tau[B]^2. \tag{S4.11}$$

Note that the first three free cumulants correspond directly to their classical counterparts but higher cumulants do not. For a thorough treatment of free cumulant expansions see for example Mingo and Speicher (2017).

In our setting, asymptotic freeness of $W_{N,T}$ and $C_N$ implies

$$\tau[\hat{C}_{N,T}^2] = \tau[C_N W_{N,T} C_N W_{N,T}] = \tau[C_N^2] + \frac{N}{T}\tau[C_N]^2 + \mathcal{O}\left(\frac{1}{T}\right). \tag{S4.12}$$

#### S5 Neural Dimensionality and Data Dimensionality

As a measure of the dimensionality, we use the "participation-ratio" dimensionality of the covariance matrix, defined as

$$D(C) := \frac{\left(\sum_k d_k\right)^2}{\sum_k d_k^2}. \tag{S5.1}$$

where $d_k$ are the eigenvalues of $C$. When all eigenvalues are identical this yields $D = N$, and when all are zero except for one this yields $D = 1$.

We can rewrite this in terms of the normalized trace ($\tau[C] = \frac{1}{N}\sum_i C_{ii} = \frac{1}{N}\sum_k d_k$) as follows:

$$D(C) = N\frac{\tau[C]^2}{\tau[C^2]}. \tag{S5.2}$$

We define the *neural dimensionality* of a network of $N$ neurons as the participation-ratio dimensionality of the ground-truth, neural covariance, $C_N$, and we write it as

$$D_N := D(C_N). \tag{S5.3}$$

For an empirical observation of $N$ neurons and $T$ trials, we define the *data dimensionality* as the participation-ratio dimensionality of the empirical, data covariance, $\hat{C}_{N,T} = \sqrt{C_N}W\sqrt{C_N}$, and write it as

$$\hat{D}_{N,T} := D(\hat{C}_{N,T}). \tag{S5.4}$$

We will study the relationship between the data dimensionality, $\hat{D}_{N,T}$, and the neural dimensionality, $D_N$, and how both depend on the numbers of neurons and trials.

#### S5.1 Dimensionality as a Function of Trials

In Section S4 we derived the relationship between the first two trace moments of the sample covariance and the population covariance:

$$\tau[\hat{C}_{N,T}] = \tau[C_N] \tag{S5.5}$$

$$\tau[\hat{C}^2_{N,T}] = \tau[C^2_N] + \frac{N}{T}\tau[C_N]^2 + \mathcal{O}\left(\frac{1}{T}\right). \tag{S5.6}$$

Substituting these values into Equation (S5.2), we find the data dimensionality as a function of trials, $T$:

$$\hat{D}_{N,T} = \frac{D_N}{1 + \frac{D_N}{T}}, \tag{S5.7}$$

up to an $\mathcal{O}\left(\frac{1}{T}\right)$ correction.

This takes an appealing form when written in terms of reciprocal dimensionalities:

$$\hat{D}^{-1}_{N,T} = D^{-1}_N + T^{-1}. \tag{S5.8}$$

From these expressions, one sees that the data dimensionality underestimates the neural dimensionality for finite $T$.

Next, we invert this relationship in order to infer the neural dimensionality from data:

$$D^{-1}_N = \hat{D}^{-1}_{N,T} - T^{-1}. \tag{S5.9}$$

Finally, we can use this to predict the data dimensionality of $T$ trials after we have observed only $P$ trials:

$$\hat{D}^{-1}_{N,T} = \hat{D}^{-1}_{N,P} + T^{-1} - P^{-1}. \tag{S5.10}$$

#### S5.2 Dimensionality as a Function of Neurons

Predicting the dimensionality for a different number of neurons is a more complicated task. In order to do so, we must specify the relationship between an $M \times M$ covariance, $C_M$, and an $N \times N$ covariance, $C_N$.

We can write the smaller matrix, $C_M$, as a subsample of the larger matrix, $C_N$:

$$C_M = SC_NS^T, \tag{S5.11}$$

where $S$ is an $M \times N$ matrix whose entries are $0$ or $1$ and that satisfies $SS^T = I_M$.

The key assumption for tractability is that the full covariance and the subsampling matrix are free (see Section S4). Intuitively, we can think of this condition as meaning that the subsampling does not align with the eigenbasis of $C_N$ in any significant way, or equivalently, that the neural modes of $C_N$ are isotropically dispersed across the neurons in the network. This is also equivalent to assuming that the spectrum of the resulting subsampled covariance, $C_M$, is statistically indistinguishable from that obtained by a random orthogonal projection. We verify that this assumption is reasonable (Figure S1), and we therefore proceed under the assumption that $S$ is a random orthonormal matrix.

We would like to predict how the neural dimensionality changes so we need the first two trace moments: $\tau[C_M]$ and $\tau[C^2_M]$.

We write

$$O = S^TS \in \mathbb{R}^{N\times N}, \tag{S5.12}$$

and we note that by the cyclic property of the trace, $\mathrm{Tr}_N[O] = \mathrm{Tr}_M[SS^T] = M$, so $\tau[O] = \frac{M}{N}$.

Again by the cyclic property, $\mathrm{Tr}_M[SC_NS^T] = \mathrm{Tr}_N[OC_N]$. Therefore we find that the normalized traces of the two covariances are equal, as we should expect:

$$\tau[C_M] = \frac{N}{M}\tau[OC_N] = \frac{N}{M}\tau[O]\tau[C_N] = \tau[C_N], \tag{S5.13}$$

For the second trace moment, we note $\mathrm{Tr}_M[C_M^2] = \mathrm{Tr}_M[SC_NS^TSC_NS] = \mathrm{Tr}_N[OC_NOC_N]$, so that after normalizing we have $\tau[C_M^2] = \frac{N}{M}\tau[OC_NOC_N]$.

And furthermore, $\mathrm{Tr}_N[O^2] = \mathrm{Tr}_N[S^TSS^TS] = \mathrm{Tr}_M[I_m] = M$, so that $\tau[O^2] = \frac{M}{N}$.

We can now use Equation (S4.11) from Section S4 for the trace of the mixed product of free matrices:

$$\tau\left(OC_NOC_N\right) = \tau[C_N]^2\left(\tau[O^2] - \tau[O]^2\right) + \tau[C_N^2]\tau[O]^2. \tag{S5.14}$$

Thus,

$$\tau\left(OC_NOC_N\right) = \frac{M}{N}\left(1 - \frac{M}{N}\right)\tau[C_N]^2 + \left(\frac{M}{N}\right)^2\tau[C_N^2], \tag{S5.15}$$

and therefore

$$\tau[C_M^2] = \frac{M}{N}\tau[C_N^2] + \left(1 - \frac{M}{N}\right)\tau[C_N]^2. \tag{S5.16}$$

Now we can calculate the neural dimensionality of the subsample:

$$D_M = M\frac{\tau[C_N]^2}{\frac{M}{N}\tau[C_N^2] + \left(1 - \frac{M}{N}\right)\tau[C_N]^2} = \frac{D_N}{1 + \left(\frac{1}{M} - \frac{1}{N}\right)D_N}. \tag{S5.17}$$

Or in terms of reciprocal dimensionality:

$$D_M^{-1} = D_N^{-1} + M^{-1} - N^{-1}. \tag{S5.18}$$

In order to predict the dimensionality of the full network, $D_N$, from the dimensionality of the subsample, $D_M$, we simply rearrange:

$$D_N^{-1} = D_M^{-1} + N^{-1} - M^{-1}. \tag{S5.19}$$

Note also that the same relationship holds for subsampling the sample covariance, i.e. between $\hat{D}_{M,P}$ and $\hat{D}_{N,P}$.

We lastly observe that by taking $N \to \infty$, we obtain

$$D_\infty^{-1} = D_M^{-1} - M^{-1}. \tag{S5.20}$$

We find that for any $D_M < M$ there is a finite $D_\infty$ limit.

Thus, for any finite-dimensional covariance $C_M$ that has data dimensionality strictly less than $M$, if we record more and more neurons, then under our assumption of isotropically distributed modes, the data dimensionality must saturate at a finite value of $D_\infty^{-1} = D_M^{-1} - M^{-1}$.

In contrast, assume that $D_N \to \infty$ as $N \to \infty$. At the extreme of $D_N \approx N$ we have $C_N \approx dI_N$ for some uniform neural mode strength $d$, and therefore any subsample or projection will yield $C_M \approx dI_M$ and $D_M \approx M$. Therefore, assuming that $D_N < N$, then rewriting Eq. (S5.18)

$$D_M = \frac{M}{1 + \frac{M}{D_N}\left(1 - \frac{D_N}{N}\right)}. \tag{S5.21}$$

Thus, any finite, $M$-dimensional projection will yield approximately

$$D_M \approx M\left(1 - \frac{M}{D_N}\left(1 - \frac{D_N}{N}\right)\right). \tag{S5.22}$$

##### S5.3 Dimensionality as a Function of Both Neurons and Trials

We would like to predict the true neural dimensionality of a larger network of $N$ neurons (i.e. the dimensionality of the ground truth $C_N$), from a recording of $M$ neurons and only $P$ trials (i.e. from the dimensionality of the smaller sample covariance, $\hat{C}_{M,P}$). To do so we return to Equation (S5.9) which infers the neural dimensionality, $D_N$, in terms of the data dimensionality of the same $N$ neurons from $P$

trials, $\hat{D}_{N,P}$: $D_N^{-1} = \hat{D}_{N,P}^{-1} - P^{-1}$. We then insert Equation (S5.19) to predict the data dimensionality of the larger set of neurons, $\hat{D}_{N,P}$, from the data dimensionality of fewer neurons, $\hat{D}_{M,P}$. Together this gives:

$$D_N^{-1} = \hat{D}_{M,P}^{-1} + N^{-1} - M^{-1} - P^{-1}, \tag{S5.23}$$

or

$$D_N = \frac{\hat{D}_{M,P}}{1 + (N^{-1} - M^{-1} - P^{-1})\,\hat{D}_{M,P}}. \tag{S5.24}$$

Finally, we would like to predict empirical data dimensionality of $T$ trials from those same $N$ neurons using the same observed data dimensionality, $\hat{D}_{M,P}$. To do so we apply Equation (S5.8):

$$\hat{D}_{N,T}^{-1} = \hat{D}_{M,P}^{-1} + N^{-1} + T^{-1} - M^{-1} - P^{-1}, \tag{S5.25}$$

or

$$\hat{D}_{N,T} = \frac{\hat{D}_{M,P}}{1 + (N^{-1} - M^{-1} + T^{-1} - P^{-1})\,\hat{D}_{M,P}}. \tag{S5.26}$$

### S6 Accuracy and Reliability of the Sample Covariance

In order to quantify both the accuracy and the reliability of the data covariance, we use the normalized squared Frobenius norm error, or Euclidean error.

The squared Frobenius norm of a matrix $A$, $\|A\|_F^2 = \sum_{i,j} A_{ij}^2$, is the sum of the squares of the elements of a matrix. For a symmetric matrix, this can be rewritten as the trace of the square of the matrix. Therefore, for any pair of $N \times N$ matrices, $A$ and $B$, we have

$$\frac{\|A\|_F^2}{\|B\|_F^2} = \frac{\tau[A^2]}{\tau[B^2]}, \tag{S6.1}$$

where again, $\tau$ is the normalized trace operator: $\tau[A] := \frac{1}{N}\sum_i A_{ii}$ for an $N \times N$ matrix, $A$.

#### S6.1 Accuracy: Similarity of Data Covariance to Neural Covariance

To quantify the accuracy of the data covariance we define the *estimation error* as the squared Frobenius norm of the difference between data covariance and true neural covariance, normalized by the squared Frobenius norm of the neural covariance:

$$\mathrm{EstErr}\left(\hat{C}_{N,T}\right) = \frac{\tau\left[\left(\hat{C}_{N,T} - C_N\right)^2\right]}{\tau[C_N^2]}. \tag{S6.2}$$

In order to calculate the accuracy, $\mathrm{EstErr}(\hat{C}_{N,T})$ we use the same relationships between trace-moments derived in Section S4 (and used in Section S5):

$$\tau[\hat{C}_{N,T}] = \tau[C_N] \tag{S6.3}$$

$$\tau[\hat{C}_{N,T}^2] = \tau[C_N^2] + \frac{N}{T}\tau[C_N]^2 + \mathcal{O}\left(\frac{1}{T}\right). \tag{S6.4}$$

We will also need

$$\tau[\hat{C}_{N,T}C_N] = \tau[C_N^2 W] = \tau[C_N^2], \tag{S6.5}$$

where for the first equality we used the circular invariance of the trace, and for the second we used asymptotic freeness of $C_N$ and $W$, along with $\tau[W] = 1$.

Looking at the raw error in the numerator of Equation (S6.2), we have

$$\tau\left[(\hat{C}_{N,T}-C_N)^2\right]=\tau\left[\hat{C}_{N,T}^2\right]+\tau\left[C_N^2\right]-2\tau\left[\hat{C}_{N,T}C_N\right]=\frac{N}{T}\tau[C_N]^2+\mathcal{O}\left(\frac{1}{T}\right). \tag{S6.6}$$

Note the relative uncertainty in $\tau\left[(\hat{C}_{N,T}-C_N)^2\right]$ is $\mathcal{O}\left(\frac{1}{N}\right)$.

We therefore have the squared error simply

$$\mathrm{EstErr}\left(\hat{C}_{N,T}\right)=\frac{N}{T}\frac{\tau[C_N]^2}{\tau[C_N^2]}. \tag{S6.7}$$

The ratio $N\frac{\tau[C_N]^2}{\tau[C_N^2]}$ is exactly the neural dimensionality, $D_N$. So we have

$$\mathrm{EstErr}\left(\hat{C}_{N,T}\right)=\frac{D_N}{T}, \tag{S6.8}$$

up to an order $\mathcal{O}\left(\frac{1}{N}\right)$ relative correction.

#### S6.2 Reliability: Similarity Between Empirical Covariances of Two Disjoint Samples

We now consider two distinct samples from the same neural covariance, $R_a=\sqrt{C}X_a$ and $R_b=\sqrt{C}X_b$. For simplicity we will focus on the case in which the two samples have the same number of trials $T_a=T_b=T$. Generalization to the case of differing numbers of trials is straightforward.

We quantify reliability via the error obtained from using one data covariance to estimate the other:

$$\mathrm{RepErr}\left(\hat{C}_{N,T}\right)=\frac{\|\hat{C}_{N,T}^{(a)}-\hat{C}_{N,T}^{(b)}\|_F^2}{\|\hat{C}_{N,T}^{(b)}\|_F^2}, \tag{S6.9}$$

or equivalently:

$$\mathrm{RepErr}\left(\hat{C}_{N,T}\right)=\frac{\tau[(\hat{C}_{N,T}^{(a)})^2]+\tau[(\hat{C}_{N,T}^{(b)})^2]-2\tau[\hat{C}_{N,T}^{(a)}\hat{C}_{N,T}^{(b)}]}{\tau[(\hat{C}_{N,T}^{(b)})^2]}. \tag{S6.10}$$

The two samples, $R_a$ and $R_b$, have data covariances, $\hat{C}_{N,T}^{(a)}=\sqrt{C}W_a\sqrt{C}$ and $\hat{C}_{N,T}^{(b)}=\sqrt{C}W_b\sqrt{C}$, where both $W_a$ and $W_b$ are sample covariances of i.i.d. noise, $W_k=\frac{1}{T_k}X_kX_k^T$ for both $k\in\{a,b\}$.

As above, we have the first two moments of $\hat{C}_{N,T}^{(a)}$, i.e.

$$\tau[\hat{C}_{N,T}^{(a)}]=\tau[C_N] \tag{S6.11}$$

$$\tau[(\hat{C}_{N,T}^{(a)})^2]=\tau[C_N^2]+\frac{N}{T_a}\tau[C_N]^2, \tag{S6.12}$$

and similarly for $\hat{C}_{N,T}^{(b)}$.

In addition, we need to find $\tau[\hat{C}_{N,T}^{(a)}\hat{C}_{N,T}^{(b)}]=\tau[C_NW_aC_NW_b]$, where we used the circular invariance of the trace. We now make use of the fact that the $W_k$ are each free random matrices with $\tau[W_k]=1$, both with regard to each other and with regard to $C_N$. Therefore

$$\begin{aligned}
\tau[\hat{C}_{N,T}^{(a)}\hat{C}_{N,T}^{(b)}]&=\tau[C_NW_aC_NW_b] && \text{(S6.13)}\\
&=\tau[C_NW_aC_N]\tau[W_b] && \text{(S6.14)}\\
&=\tau[C_N^2W_a] && \text{(S6.15)}\\
&=\tau[C_N^2]. && \text{(S6.16)}
\end{aligned}$$

As mentioned above, for simplicity we focus on the case $T_a=T_b=T$. Plugging into Equation (S6.10), we find the replication error in terms of the true neural dimensionality, $D_N$:

$$\mathrm{RepErr}\left(\hat{C}_{N,T}\right)=2\frac{\frac{N}{T}\tau[C_N]^2}{\tau[C_N^2]+\frac{N}{T}\tau[C_N]^2}=2\frac{D_N}{T+D_N}. \tag{S6.17}$$

This can be rewritten as $\frac{2}{T}\frac{1}{D_N^{-1}+T^{-1}}$, and we recognize the denominator as exactly the inverse of the data dimensionality of each of the empirical samples, $\hat{D}_{N,T}$ (Eq. (S5.8)). Therefore in terms of the data dimensionality we have simply

$$\mathrm{RepErr}\left(\hat{C}_{N,T}\right) = 2\frac{\hat{D}_{N,T}}{T}. \tag{S6.18}$$

Finally, we can use the data dimensionality from $M$ neurons and $P$ trials to extrapolate and predict the replication error between two datasets of $N$ neurons and $T$ trials, by simply inserting Equation (S5.25) for $\hat{D}_{N,T}$:

$$\mathrm{RepErr}\left(\hat{C}_{N,T}\right) = \frac{2}{T}\left(\hat{D}_{M,P}^{-1} + N^{-1} - M^{-1} + T^{-1} - P^{-1}\right)^{-1}. \tag{S6.19}$$

### S7 Incorporating Trial-to-Trial Variability in Overall Power

Throughout this study, we normalized individual trials as part of the data preprocessing stage. In this section we examine the implications of *not* performing this normalization, with the aim of both justifying the normalization and providing predictions of accuracy and reliability in its absence.

We consider a model with a trial-by-trial power factor, $\alpha_t > 0$, which sets the power (mean-squared) of activity over neurons on trial $t$:

$$\boldsymbol{r}_t = \sqrt{C_N}\boldsymbol{x}_t\sqrt{\alpha_t}. \tag{S7.1}$$

The overall scale of the $\alpha_t$ can be chosen arbitrarily by simply absorbing it into $C_N$, so we choose to set the average equal to 1, $\langle\alpha\rangle = 1$

In matrix form, we have:

$$R_{N,T} = \sqrt{C_N}X_{N,T}\sqrt{A_T}, \tag{S7.2}$$

where $A$ is a diagonal matrix.

Then we can write $\hat{C}_{N,T} = \sqrt{C_N}\tilde{W}_{N,T}\sqrt{C_N}$, where

$$\tilde{W}_{N,T} = \frac{1}{T}X_{N,T}A_T X_{N,T}^T. \tag{S7.3}$$

Recall that by construction $\tau[A_T] = 1$.

We will need the first two trace moments of $\tilde{W}_{N,T}$. Suppressing the dependence on $N$ and $T$ momentarily, we have $\tilde{W}_{ij} = \frac{1}{T}\sum_{k=1}^{T}\alpha_k X_{ik}X_{jk}$. The scale factors are assumed to be independent of $X_{ik}$ so that $\tau[\tilde{W}_{N,T}] = \left\langle X_{ik}^2\right\rangle\langle\alpha_k\rangle = 1$.

For the second trace moment:

$$\tau[\tilde{W}_{N,T}^2] = \frac{1}{NT^2}\sum_{k,l=1}^{T}\sum_{i,j=1}^{N} X_{ik}\alpha_k X_{jk}X_{jl}\alpha_l X_{il}. \tag{S7.4}$$

The terms that contribute are those that pair the Gaussian variables, i.e. those with either $i = j$ or $k = l$. There are $NT^2$ terms with $i = j$, which are of the form $X_{ik}^2\alpha_k X_{il}^2\alpha_l$. Each of these terms has expected value $\left\langle X_{ik}^2\right\rangle^2\langle\alpha_k\rangle^2 = 1$. There are $N^2T$ terms with $k = l$, each of the form $X_{ik}^2X_{jk}^2\alpha_k^2$, which have expected value $\left\langle\alpha^2\right\rangle$. Thus, we have

$$\tau[\tilde{W}_{N,T}^2] = 1 + \left\langle\alpha^2\right\rangle\frac{N}{T}, \tag{S7.5}$$

which is consistent with $\tau[W_{N,T}^2] = 1 + \frac{N}{T}$ for the case of identical trials.

We have $\tau[\hat{C}_{N,T}^2] = \tau[C_N\tilde{W}_{N,T}C_N\tilde{W}_{N,T}]$, and using Equation (S4.11), we can write

$$\tau[\hat{C}_{N,T}^2] = \tau[{C_N}^2] + \tau[C_N]^2\left(\tau[\tilde{W}_{N,T}^2] - 1\right), \tag{S7.6}$$

where we have inserted $\tau[\tilde{W}_{N,T}]^2 = 1$.

We therefore have

$$\tau[\hat{C}^2_{N,T}] = \tau[C^2_N] + \langle\alpha^2\rangle \frac{N}{T}\tau[C_N]^2. \tag{S7.7}$$

Recall that in the setting of identical trials ($\alpha_k = 1$ for all $k$), we had $\tau[\hat{C}^2_{N,T}] = \tau[C^2_N] + \frac{N}{T}\tau[C_N]^2$. Thus, the dispersion of the scale factor effectively reduces the number of trials, i.e. we replace $\frac{N}{T}$ with $\langle\alpha^2\rangle \frac{N}{T}$ in our expressions.

In particular, the data dimensionality given $T$ trials will be:

$$\hat{D}_{N,T} = \frac{D_N}{1 + \frac{\langle\alpha^2\rangle}{T} D_N}, \tag{S7.8}$$

or

$$\hat{D}^{-1}_{N,T} = D^{-1}_N + \frac{\langle\alpha^2\rangle}{T}. \tag{S7.9}$$

The equation for $D_N$ as a function of $N$ (Eq. (S5.18)) is unchanged, so given an observation of $M$ neurons and $P$ trials, we can infer the asymptotic dimensionality, $D_\infty$, as

$$D_\infty = \frac{\hat{D}_{M,P}}{1 - \left(\frac{1}{M} + \frac{\langle\alpha^2\rangle}{P}\right)\hat{D}_{M,P}}. \tag{S7.10}$$

Given $D_\infty$, we can predict the dimensionality for $N > M$ neurons and $T > P$ trials:

$$\hat{D}_{N,T} = \frac{D_\infty}{1 + \left(\frac{1}{N} + \frac{\langle\alpha^2\rangle}{T}\right) D_\infty}. \tag{S7.11}$$

We can predict $\hat{D}_{N,T}$ directly from $\hat{D}_{M,P}$:

$$\hat{D}_{N,T} = \frac{\hat{D}_{M,P}}{1 + [N^{-1} - M^{-1} + \langle\alpha^2\rangle (T^{-1} - P^{-1})]\,\hat{D}_{M,P}}, \tag{S7.12}$$

or equivalently:

$$\hat{D}^{-1}_{N,T} = \hat{D}^{-1}_{M,P} + \langle\alpha^2\rangle \left(\frac{1}{T} - \frac{1}{P}\right) + \frac{1}{N} - \frac{1}{M}. \tag{S7.13}$$

Similarly, the predicted estimation error of the sample covariance will be

$$\mathrm{EstErr}\left(\hat{C}_{N,T}\right) = \frac{\langle\alpha^2\rangle}{T} D_N, \tag{S7.14}$$

and the replication error between two sample covariances from $T$ is:

$$\mathrm{RepErr}\left(\hat{C}_{N,T}\right) = 2\frac{D_N}{D_N + \frac{T}{\langle\alpha^2\rangle}} = 2\frac{\langle\alpha^2\rangle}{T}\hat{D}_{N,T}. \tag{S7.15}$$

Finally, we can predict the replication error between two datasets of $N$ neurons and $T$ trials, from the data dimensionality of a subsample of $M$ neurons and $P$ trials:

$$\mathrm{RepErr}\left(\hat{C}_{N,T}\right) = 2\frac{\langle\alpha^2\rangle}{T}\left(\hat{D}^{-1}_{M,P} + N^{-1} - M^{-1} + \langle\alpha^2\rangle\left(T^{-1} - P^{-1}\right)\right)^{-1}. \tag{S7.16}$$

As can be seen from Equations (S7.14) and (S7.15), both the estimation error and replication error increase with growing variability in the trial-to-trial scale factor. Thus we can obtain a more accurate and more reliable sample covariance by first normalizing the data to remove the trial-to-trial variability in the overall scale.

In practice, therefore, we add a preprocessing step to normalize each trial's activity: Given the data matrix, $\tilde{R}_{it}$, we write:

$$R_{it} = \frac{1}{\sqrt{\hat{\alpha}_t/\langle\hat{\alpha}_t\rangle}}\tilde{R}_{it}, \tag{S7.17}$$

where $\hat{\alpha}_t$ is the mean-square activity of trial $t$:

$$\hat{\alpha}_t = \frac{1}{N}\sum_{i=1}^{N}\tilde{R}_{it}^2, \tag{S7.18}$$

and $\langle\hat{\alpha}_t\rangle$ is the average mean-square activity over all trials. Then we model the resulting data matrix as $R = \sqrt{C}X$.

### S8 Low-Rank Plus Noise Model for Covariance Structure

In the previous sections we made no assumptions about the structure of the unknown neural covariance matrix. We now study the case of a low-rank "spiked" model for the covariance.

In the low-rank plus noise model we assume that the data is generated by $K$ latent variables, each with signal strength $s_k^2$ driving the $N$ neurons via a set of orthonormal signal mode vectors $\boldsymbol{u}_k$, together with additive independent identically distributed noise of variance $\sigma^2$. Writing the signal strengths in non-increasing order along a diagonal matrix $S$, and the signal mode vectors as columns of the matrix $U$, we represent the population activity on trial $t$ as

$$\boldsymbol{r}_t = US\boldsymbol{y}_t + \sigma\boldsymbol{\eta}_t, \tag{S8.1}$$

where $\boldsymbol{y}_t \in \mathbb{R}^K$ is the $K$-dimensional signal or latent state, which for simplicity we assume to be standard normal, and $\boldsymbol{\eta}_t \in \mathbb{R}^N$ is a vector of i.i.d. noise.

Observe that the covariance of $US\boldsymbol{y}_t$ is $US^2U^T$, while the covariance of $\boldsymbol{\eta}_t$ is the identity, $I$, by definition. Thus, the neural covariance is the sum of a low-rank component plus the scaled identity matrix:

$$C_N = US^2U^T + \sigma^2 I. \tag{S8.2}$$

The neural mode strengths, i.e. eigenvalues of the true covariance matrix, are then given by $d_k = s_k^2 + \sigma^2$ for $k \leq K$ and $d_k = \sigma^2$ for $k > K$.

We note that the elements of the covariance matrix have units of the square of neural activity, and that the signal component of the covariance is $\sum_{k=1}^{K} s_k^2 u_{k_i} u_{k_j}$. Assuming the components of the eigenvectors $u_k$ are spread isotropically across the neurons, the product of components $u_{k_i}u_{k_j}$ has order of magnitude $\frac{1}{N}$ due to the unit-norm normalization of each eigenvector $\boldsymbol{u}_k$. However, in an experimental setting, the actual scale of the single neuron variances and neuron pair covariances are expected to be $O(1)$ quantities that do not change with the number of recorded neurons $N$. This requires that the eigenvalues $s_k^2$ of $C_N$ are $O(N)$ to compensate for the $O(\frac{1}{N})$ contribution from eigenvector components. We therefore define $K$ signal-to-noise ratios that are independent of $N$ and $T$, each associated with its corresponding rank-$1$ mode:

$$\mathrm{SNR}_k := \frac{s_k^2}{N\sigma^2}. \tag{S8.3}$$

We can think of $\frac{s_k^2}{\sigma^2}$ as the $O(N)$ signal-to-noise ratio of mode $k$, while $\mathrm{SNR}_k$ is the corresponding $O(1)$ signal-to-noise ratio of mode $k$, *per neuron*. Crucially, we assume that each mode $k$'s *per-neuron* signal-to-noise ratio $\mathrm{SNR}_k$ does not change as a function of the number of recorded neurons $N$ and trials $T$.

Just as above, the data covariance is given by

$$\hat{C}_{N,T} = \sqrt{C_N}W_{N,T}\sqrt{C_N}, \tag{S8.4}$$

where $W_{N,T} = \frac{1}{T}X_{N,T}X_{N,T}^T$, with $X_{N,T}$ being an $N \times T$ matrix of i.i.d. zero mean unit variance elements with finite fourth moments.

We will continue to write the eigenvalue decomposition of the sample covariance as

$$\hat{C}_{N,T} = \hat{U}\hat{D}\hat{U}^T = \sum_{k=1}^{N} \hat{d}_k \hat{\boldsymbol{u}}_k \hat{\boldsymbol{u}}_k^T. \tag{S8.5}$$

#### S8.1 Review of Low-Rank Theory

We briefly review the theory predicting the data mode strengths, i.e. sample eigenvalues, and the relationship between the data mode vectors, sample eigenvectors, and ground truth eigenvectors. These are existing results in the literature (eg Shabalin and Nobel (2013); Loubaton and Vallet (2011); Benaych-Georges and Nadakuditi (2012)), but for completeness we present derivations below in Sections S10.1 and S10.2.

In the random matrix literature, the eigenvalue spectra of the matrices at hand are constructed to have a well-defined $\mathcal{O}(1)$ limit as the matrix size is taken to infinity with a fixed aspect ratio $q = \frac{N}{T}$.

In the absence of low-rank structure, i.e. with true covariance $C_N = \sigma^2 I$, the eigenvalue spectrum of $\hat{C}_{N,T}$ is given by the well-known Marchenko–Pastur (MP) distribution with bounds $\hat{d}_\pm = \sigma^2(1 \pm \sqrt{q})^2$ (ignoring the $N-T$ zero eigenvalues whenever $N > T$; Marčenko and Pastur (1967)).

From the perspective of random matrix theory, the low-rank plus noise model fixes a finite number $K$ modes with "spiked" eigenvalues, i.e. each increased by an amount $s_k^2$ above the homogeneous noise level $\sigma^2$. We report existing results on the statistics of the sample eigenvalues, $\hat{d}_k$, and eigenvectors, $\hat{\boldsymbol{u}}_k$, as a function of $q$.

First, the sample eigenvalues contain a "bulk distribution" of $N-K$ eigenvalues that are indistinguishable from the MP distribution.

Each of the $K$ signal eigenvalues $s_k^2$ is visible as a corresponding outlier eigenvalue $\hat{d}_k$ in the data only if the signal is greater than a critical threshold. Otherwise, the signal mode is absorbed into the upper edge of the bulk:

$$\hat{d}_k = \begin{cases} s_k^2\left(1 + \frac{\sigma^2}{s_k^2}\right)\left(1 + q\frac{\sigma^2}{s_k^2}\right) & \text{if } s_k^2 \geq \sqrt{q}\,\sigma^2 \\ \sigma^2\left(1+\sqrt{q}\right)^2 & \text{if } s_k^2 < \sqrt{q}\,\sigma^2 \end{cases}. \tag{S8.6}$$

The corresponding data eigenvectors are each only partially aligned with their corresponding ground truth eigenvectors, again subject to the same signal strength threshold:

$$|\hat{\boldsymbol{u}}_k \cdot \boldsymbol{u}_j|^2 = \begin{cases} \delta_{jk}\frac{s_k^2 - \frac{q}{s_k^2}}{s_k^2 + q} & \text{if } s_k^2 \geq \sqrt{q}\,\sigma^2 \\ 0 & \text{if } s_k^2 < \sqrt{q}\,\sigma^2 \end{cases}. \tag{S8.7}$$

For simplicity we have assumed that the true eigenvalues are non-degenerate, i.e. $s_k \neq s_j$ for $k \neq j$. If there are degenerate eigenvalues then the alignment formula in Equation (S8.7) predicts the square norm of the sample eigenvector's projection into the corresponding eigenvector subspace. See Section S10 for derivations of (S8.6) and (S8.7).

#### S8.2 Fitting the Model

As detailed in the Methods Model-fit, we set the inferred noise variance, $\hat{\sigma}^2$, in order to fit the median *non-zero* sample eigenvalue in the data, $\hat{d}_{\text{median}}$, with an MP distribution of the same variance:

$$\hat{\sigma}^2 = \frac{\hat{d}_{\text{median}}}{\mu_{\text{median}}}. \tag{S8.8}$$

where $\mu_{median}$ is the median of the MP distribution and is found numerically by solving

$$\frac{1}{2} = \frac{1}{2\pi \min(q,1)} \int_{\delta_-}^{\mu_{median}} dx\, \frac{\sqrt{(\delta_+ - x)(x - \delta_-)}}{x}, \tag{S8.9}$$

with $\delta_\pm = \left(1 \pm \sqrt{q}\right)^2$.

Given our noise estimate, we can infer the signal-to-noise ratios of the potential signal modes. Every sample eigenvalue above the inferred noise ceiling, $\hat{d}_+ = \hat{\sigma}^2 \left(1 + \sqrt{q}\right)^2$, is a potential signal mode.

Inserting the signal mode strength in terms of SNR $s_k^2 = N\hat{\sigma}^2\,\mathrm{SNR}_k$, into our forward model for the data mode strengths (Eq. (S8.6)), yields

$$\hat{d}_k = \hat{\sigma}^2 \left( N\,\mathrm{SNR}_k + \frac{1}{T\,\mathrm{SNR}_k} + 1 + q \right). \tag{S8.10}$$

Interestingly, $\hat{\sigma}^2\,(1+q)$ is exactly the midpoint of the noise sea: $\hat{d}_{\mathrm{mid}} = \frac{1}{2}\left(\hat{d}_- + \hat{d}_+\right)$. We therefore define a kind of empirical SNR by subtracting $\hat{d}_{\mathrm{mid}}$ from $\hat{d}_k$, and dividing by the total noise variance $N\hat{\sigma}^2$:

$$\widetilde{\mathrm{SNR}}_k \equiv \frac{\hat{d}_k - \hat{d}_{\mathrm{mid}}}{N\hat{\sigma}^2}. \tag{S8.11}$$

From Equation (S8.10) this yields

$$\widetilde{\mathrm{SNR}}_k = \mathrm{SNR}_k + \frac{1}{NT\,\mathrm{SNR}_k}, \tag{S8.12}$$

which gives a quadratic equation for the underlying SNR:

$$\mathrm{SNR}_k^2 - \widetilde{\mathrm{SNR}}_k \mathrm{SNR}_k + \frac{1}{NT} = 0, \tag{S8.13}$$

and finally, inverting this formula yields the inferred SNR as the positive quadratic root,

$$\widehat{\mathrm{SNR}}_k = \frac{\widetilde{\mathrm{SNR}}_k + \sqrt{\widetilde{\mathrm{SNR}}_k^2 - \frac{4}{NT}}}{2}. \tag{S8.14}$$

#### S8.3 The Impact of Neuron and Trial Counts on Signal Geometry

We derive predictions for the impact of the number of neurons and trials on the inferred signal geometry.

By assumption, the key parameters of our model, the noise variance ($\sigma^2$) and each mode's per-neuron signal-to-noise ratio ($\mathrm{SNR}_k$) are independent of the numbers of neurons and trials, and therefore the predictions are straightforward. Recall that the eigenvalues of the low-rank component of the covariance corresponding to each signal mode are given by $s_k^2 = N\sigma^2\,\mathrm{SNR}_k$.

Thus, we first conclude, from the critical eigenvalue threshold, $s^2_{\mathrm{crit}} = \sqrt{q}\,\sigma^2$, that the critical SNR is:

$$\mathrm{SNR}_{\mathrm{crit}} = \frac{1}{\sqrt{NT}}. \tag{S8.15}$$

Next, writing the signal–data mode relation (S8.6) in terms of SNRs, we have

$$\hat{d}_k(N,T) = \begin{cases} N\,\mathrm{SNR}_k\,\sigma^2 \left(1 + \frac{1}{N\mathrm{SNR}_k}\right)\left(1 + \frac{1}{T\mathrm{SNR}_k}\right) & \text{if } \mathrm{SNR}_k \geq \frac{1}{\sqrt{NT}} \\ \sigma^2 \left(1 + \sqrt{\frac{N}{T}}\right)^2 & \text{if } \mathrm{SNR}_k < \frac{1}{\sqrt{NT}} \end{cases}. \tag{S8.16}$$

This relation gives a method for predicting data modes for $N$ neurons and $T$ trials from data with $M$ neurons and $P$ trials: infer SNRs from the $M \times P$ subsampled data, and for each mode above threshold in the subsample (i.e. $\mathrm{SNR}_k > \frac{1}{\sqrt{MP}}$), plug the inferred SNR into (S8.16) to obtain $\hat{d}_k(N,T)$.

##### S8.3.1 Signal Mode Strength Inflation

In order to examine the signal geometry and the way that it changes as a function of neurons and trials, we compute the ratio between consecutive eigenvalues, which is equivalent to the ratio of variances in PCA.

Using Equation (S8.16), we have:

$$\frac{\hat{d}_k}{\hat{d}_{k+1}} = \frac{\mathrm{SNR}_k + \frac{1}{NT\mathrm{SNR}_k} + \frac{1}{N} + \frac{1}{T}}{\mathrm{SNR}_{k+1} + \frac{1}{NT\mathrm{SNR}_{k+1}} + \frac{1}{N} + \frac{1}{T}}, \tag{S8.17}$$

which is hyperbolic and symmetric in $N$ and $T$, and approaches the ground truth ratio $\frac{\mathrm{SNR}_k}{\mathrm{SNR}_{k+1}}$ in the limit of large $N$ and $T$.

We find the iso-ratio curves as a function of the ratio $R_k = \frac{\hat{d}_k}{\hat{d}_{k+1}}$:

$$T(N, R_k) = \frac{N(R_k - 1) + \frac{R_k}{\mathrm{SNR}_{k+1}} - \frac{1}{\mathrm{SNR}_k}}{N\left(\mathrm{SNR}_k - R_k \mathrm{SNR}_{k+1}\right) + 1 - R_k}. \tag{S8.18}$$

##### S8.3.2 Signal Mode Vector Misalignment

To predict the impact of neuron and trial counts on signal mode vectors, we insert the single-neuron, single-mode SNRs ($s_k^2 = N\sigma^2\,\mathrm{SNR}_k$), into Equation (S8.7) for the data mode alignments, and rearrange, yielding

$$\mathcal{A}_k = \frac{T\,\mathrm{SNR}_k}{T\,\mathrm{SNR}_k + 1}\left(1 - \frac{1}{NT\,\mathrm{SNR}_k^2}\right), \tag{S8.19}$$

for $\mathrm{SNR}_k > \frac{1}{\sqrt{NT}}$, and $0$ otherwise.

For a fixed number of neurons, $N$, this formula predicts perfect alignment in the limit of infinite trials, $\mathcal{A}_k \to 1$. For fixed number of trials, $T$, however, the alignment saturates at $\mathcal{A}_k \to \frac{T\,\mathrm{SNR}_k}{T\,\mathrm{SNR}_k + 1}$, despite infinite neurons. This result has a standard form of a ratio of signal variance accumulated over $T$ samples to the total variance, and makes intuitive sense because the number of degrees of freedom in the true eigenvector is growing with this limit. Nevertheless, for finite $N$ and $T$, this formula predicts that the alignment increases monotonically with the number of neurons observed.

##### S8.3.3 Data Mode Reliability

We define the data mode reliability, $\mathcal{R}_k$, as the squared overlap between the $k$th data mode in one trial split and the $k$th data mode in a second trial split:

$$\mathcal{R}_k \equiv \left|\hat{\boldsymbol{u}}_k^{(1)} \cdot \hat{\boldsymbol{u}}_k^{(2)}\right|^2. \tag{S8.20}$$

Given that the alignment of the $k$th mode from the $n$th data split with the true $k$th signal mode is $\mathcal{A}_k^{(n)}$, we now show that

$$\mathcal{R}_k = \mathcal{A}_k^{(1)}\mathcal{A}_k^{(2)}. \tag{S8.21}$$

We can decompose the data mode vector, $\hat{\boldsymbol{u}}_k^{(n)}$, as the projection onto the signal-mode vector, $\boldsymbol{u}_k$, and an orthogonal unit vector, $\boldsymbol{v}_k^{(n)}$, i.e. $\boldsymbol{u}_k \cdot \boldsymbol{v}_k^{(n)} = 0$:

$$\hat{\boldsymbol{u}}_k^{(n)} = \sqrt{\mathcal{A}_k^{(n)}}\boldsymbol{u}_k + \sqrt{1 - \mathcal{A}_k^{(n)}}\boldsymbol{v}_k^{(n)}. \tag{S8.22}$$

Now we consider the $k$th mode on two distinct trial splits. We write

$$\hat{\boldsymbol{u}}_k^{(1)} \cdot \hat{\boldsymbol{u}}_k^{(2)} = \sqrt{\mathcal{A}_k^{(1)}\mathcal{A}_k^{(2)}} + \sqrt{\left(1 - \mathcal{A}_k^{(1)}\right)\left(1 - \mathcal{A}_k^{(2)}\right)}\boldsymbol{v}_k^{(1)} \cdot \boldsymbol{v}_k^{(2)}. \tag{S8.23}$$

By assumption, trials are independent and therefore $\boldsymbol{v}_k^{(1)}$ and $\boldsymbol{v}_k^{(2)}$ are independently sampled unit vectors from the subspace orthogonal to $\boldsymbol{u}_k$. Thus, $|\boldsymbol{v}_k^{(1)} \cdot \boldsymbol{v}_k^{(2)}| = \mathcal{O}\left(\frac{1}{N}\right)$ with high probability, and we have:

$$\left|\hat{\boldsymbol{u}}_k^{(1)} \cdot \hat{\boldsymbol{u}}_k^{(2)}\right|^2 = \mathcal{A}_k^{(1)} \mathcal{A}_k^{(2)}. \tag{S8.24}$$

almost surely in the large $N$, $T$ limit.

###### S8.3.4 Discussion of Subspace Overlap

Recall that the low-rank model data can be considered as sampling i.i.d. columns from a normal distribution with mean $0$ and neural covariance

$$C = US^2U^T + \sigma^2 I. \tag{S8.25}$$

And the $N \times T$ data matrix can be written

$$R = \sqrt{C}X, \tag{S8.26}$$

where $X \in \mathbb{R}^{N \times T}$ is a standard normal matrix.

Now, let $\boldsymbol{u}_k$ be a signal mode vector of $C$, and let $w$ be an eigenvalue (soft) windowing function to be applied to the data's eigenvalues. The total squared overlap of the ground truth signal vector onto the data's windowed spaces is

$$\mathcal{A}_k^s(w) := \boldsymbol{u}_k^T w \left(\frac{1}{T}\sqrt{C}XX^T\sqrt{C}\right) \boldsymbol{u}_k. \tag{S8.27}$$

Note that in the limit $N, T \to \infty$, all cross-overlaps tend toward $0$ and so $\mathcal{A}_k^s(w)$ reduces to the overlap with the $k^{th}$ outlier space with a weight of $w(\hat{d}_k)$. Thus, when $w(\hat{d}_k) = 1$, we have $\mathcal{A}_k^s(w) \to \mathcal{A}_k$.

The Gaussian Poincare inequality can be used to show

$$\operatorname{std}\left(\mathcal{A}_k^s(w)\right) \leq \frac{2\kappa\left(s_k^2 + \sigma^2\right)}{\sqrt{T}} \|w'\|_\infty, \tag{S8.28}$$

for some small constant $\kappa$, where $\|w'\|_\infty$ is the maximum absolute value attained by the windowing function's derivative.

The previous inequality generally illustrates how the problem of signal-data overlap "leaking" into nearby data spaces can be mitigated by accumulating overlap over a wider range of data modes: Suppose $\boldsymbol{u}_k$ has significant cross-overlap within a tight cluster of data modes. Singling out individual data modes in a cluster requires a sharp windowing function $w$, meaning $\|w'\|_\infty$ is large and the variance of $\mathcal{A}_k^s(w)$ can be large - as is often observed in practice. On the other hand, including all of the cluster's data modes in the windowing function $w$ allows for gentler drop-off of $w$ after the edges of the cluster, decreasing $\|w'\|_\infty$, and decreasing the bound for the variance of $\mathcal{A}_k^s(w)$. Thus, roughly speaking, the total overlap onto a tight, well-isolated cluster of data modes is nearly deterministic.

#### S9 Low-Rank Theory for Masked Linear Autoencoder

To assess the predictive utility of the low-rank structure, and to derive a principled rule for optimal dimensionality reduction, we implement a masked-neuron linear autoencoder.

We measure the sample covariance from training trials and choose a number of data modes, $\hat{K}$, to use for prediction. These will serve as hidden units in the masked autoencoder.

On each test trial, we mask a small number of neurons (in numerical tests we restrict to one held-out neuron per test trial) and compute the $\hat{K}$ projections onto each of the $\hat{K}$ modes using only the non-held-out neurons. Then we attempt to reconstruct the activity of the held-out neurons by weighting each mode by the neurons' corresponding eigenvector loadings, i.e. feature vectors. This approach can be thought of as a form of PCA cross-validation, and provides a principled method for dimensionality reduction by optimizing $\hat{K}$ to maximize variance explained on held-out neuron single-trial activity.

We assume that we have observed $T$ training trials

$$R = USY + Z, \tag{S9.1}$$

where $R$ and $Z$ are $N$ by $T$ matrices, and $Y$ is a $K$ by $T$ matrix. We compute the sample covariance, $\hat{C} = \frac{1}{T}RR^T$, and we would like to know to what extent a fixed number $\hat{K}$ of its eigenvectors can predict held-out neuron activity.

We are presented with a test trial, yielding an $N$-dimensional neural activity vector $\boldsymbol{r}$ given by

$$\boldsymbol{r} = US\boldsymbol{y} + \boldsymbol{z}. \tag{S9.2}$$

In what follows, a necessary assumption is that the number of held-out neurons comprises a vanishing fraction of the total number of neurons, i.e. as $N$ and $T$ go to infinity, the number of held-out neurons is fixed. For notational simplicity, however, we restrict our derivation to a single held-out neuron.

We split this activity into that of a single held-out neuron, and the rest of the $N-1$ neurons. We denote the held-out neuron activity by

$$r_0 = \boldsymbol{u}_0^T S\boldsymbol{y} + z_0, \tag{S9.3}$$

where $\boldsymbol{u}_0^T$ is a single row of $U$, i.e. a $1 \times K$ row vector, so that $r_0$ is a scalar. Note that we can write this as $r_0 = \sum_{k=1}^{K} u_{0_k} s_k y_k + z_0$. We denote the rest of the test trial neural activity of the other $N-1$ neurons by

$$\boldsymbol{r}_{-0} = U_{-0}S\boldsymbol{y} + \boldsymbol{z}_{-0}. \tag{S9.4}$$

By a slight abuse of notation we denote the matrix consisting of the first $\hat{K}$ sample eigenvectors as $\hat{U}$ (rather than the full matrix of sample eigenvectors), and we denote by $\hat{\boldsymbol{u}}_0^T$ the row of this matrix corresponding to the held-out neuron (i.e. the first $\hat{K}$ sample eigenvector components evaluated on the held-out neuron).

Our prediction for the activity of the held-out neuron is simply

$$\hat{r}_0 = \hat{\boldsymbol{u}}_0^T \hat{U}_{-0}^T \boldsymbol{r}_{-0}. \tag{S9.5}$$

Inserting Equation (S9.4), we have:

$$\hat{r}_0 = \hat{\boldsymbol{u}}_0^T \hat{U}_{-0}^T U_{-0} S\boldsymbol{y} + \eta_0, \tag{S9.6}$$

where

$$\eta_0 = \hat{\boldsymbol{u}}_0^T \hat{U}_{-0}^T \boldsymbol{z}_{-0}. \tag{S9.7}$$

This noise term sums $N\hat{K}$ terms each with variance $\frac{1}{N^2}$, therefore $\eta_0 \sim \mathcal{N}\left(0, \frac{\hat{K}}{N}\right)$.

Observe that $\hat{U}_{-0}^T U_{-0}$ is a $\hat{K} \times K$ matrix of approximate alignments between sample covariance eigenvectors and ground truth population eigenvectors. Because $1 \ll N$, we approximate $\hat{U}_{-0}^T U_{-0} \approx \hat{U}^T U$. This is simply the matrix of (non-squared) dot products between ground truth eigenvectors and sample eigenvectors. In the limit of large $N$ and $T$, the off-diagonal elements of this matrix are approximately zero, while the diagonal elements are $(\hat{\boldsymbol{u}}_k \cdot \boldsymbol{u}_k)$ (in Section S10.2 we find $(\hat{\boldsymbol{u}}_k \cdot \boldsymbol{u}_k)^2 = \mathcal{A}_k$). Therefore we can write

$$\hat{r}_0 \approx \sum_{\min\{K,\hat{K}\}} \hat{u}_{0_k} \left(\hat{\boldsymbol{u}}_k \cdot \boldsymbol{u}_k\right) s_k y_k + \eta_0. \tag{S9.8}$$

We now compute the variance explained, $R^2$, by our prediction. We begin with the un-normalized error, which we will average over test trials as well as neurons, assuming isotropic statistics of the $U$:

$$\left\langle (r_0 - \hat{r}_0)^2 \right\rangle = \left\langle r_0^2 \right\rangle + \left\langle \hat{r}_0^2 \right\rangle - 2 \left\langle r\hat{r}_0 \right\rangle. \tag{S9.9}$$

Recall that, by construction, the diagonal elements of $S$ are each $N$ times the per-neuron variance of the corresponding signal mode: $s_k^2 = N\mathrm{SNR}_k$ (where we've assumed $\sigma^2 = 1$ for simplicity). Additionally, the eigenvector components have squared norm $\frac{1}{N}$ on average. Thus, $\left\langle (u_{0_k} s_k y_k)^2 \right\rangle = \mathrm{SNR}_k$, and using $r_0 = \sum_{k=1}^{K} u_{0_k} s_k y_k + z_0$ (Eq. (S9.3)) we find $\left\langle r_0^2 \right\rangle = \sum_{k=1}^{K} \mathrm{SNR}_k + 1$.

Similarly, using $\hat{r}_0 \approx \sum_{k=1}^{\min\{K,\hat{K}\}} \hat{u}_{0_k} \left(\hat{\boldsymbol{u}}_k \cdot \boldsymbol{u}_k\right) s_k y_k + \eta_0$, since the latents are independent, the mean square is the sum of the squares of each term. Using $\left(\hat{\boldsymbol{u}}_k \cdot \boldsymbol{u}_k\right)^2 = \mathcal{A}_k$, we find

$$\left\langle \hat{r}_0^2 \right\rangle = \sum_{k=1}^{\hat{K}} \mathcal{A}_k \mathrm{SNR}_k + \frac{\hat{K}}{N}, \tag{S9.10}$$

where $\frac{\hat{K}}{N}$ accounts for the variance of $\eta_0$.

We also find

$$\left\langle r_0 \hat{r}_0 \right\rangle = \sum_{k=1}^{\min\{K,\hat{K}\}} \left\langle \hat{u}_{0_k} u_{0_k} \right\rangle \left\langle \hat{\boldsymbol{u}}_k \cdot \boldsymbol{u}_k \right\rangle s_k^2 = \sum_{k=1}^{\hat{K}} \mathcal{A}_k \mathrm{SNR}_k. \tag{S9.11}$$

Note that we have safely ignored the $\min\{K, \hat{K}\}$ in the previous two equations because $\mathcal{A}_k = 0$ for $k > K$.

Combining these results we have

$$\left\langle \left(r_0 - \hat{r}_0\right)^2 \right\rangle = \sum_{k=1}^{K} \mathrm{SNR}_k - \sum_{k=1}^{\hat{K}} \mathrm{SNR}_k \mathcal{A}_k + 1 + \frac{\hat{K}}{N}. \tag{S9.12}$$

This yields for the $R^2 \equiv 1 - \frac{\left\langle \left(r_0 - \hat{r}_0\right)^2 \right\rangle}{\left\langle r_0^2 \right\rangle}$:

$$R^2 = \frac{\sum_{k=1}^{\hat{K}} \mathcal{A}_k \mathrm{SNR}_k - \frac{\hat{K}}{N}}{\sum_{k=1}^{K} \mathrm{SNR}_k + 1}, \tag{S9.13}$$

where $K$ is the ground truth signal dimension, and the alignments are (rearranged here from Equation (S8.7) for convenience):

$$\mathcal{A}_k = \frac{1}{N\,\mathrm{SNR}_k} \frac{NT\,\mathrm{SNR}_k^2 - 1}{T\,\mathrm{SNR}_k + 1}. \tag{S9.14}$$

##### S9.1 Predictive Dimensionality – Optimal Number of Hidden Units

Equation (S9.13) predicts the variance explained by the masked autoencoder on held-out trials as a function of the number of hidden units, i.e. the number of data modes, used for prediction. In order to obtain the optimal predictive dimensionality, we observe that $R^2_{\hat{K}} - R^2_{\hat{K}-1} \propto \mathcal{A}_k \mathrm{SNR}_k - \frac{1}{N}$. Writing the alignment as a function of SNR for fixed $N$ and $T$, $\mathcal{A}(x) = \frac{1}{Nx} \frac{NT\,x^2 - 1}{Tx + 1}$ and we can find the optimal SNR threshold, $\mathrm{SNR}_{\mathrm{th}}$ by setting

$$\mathcal{A}\left(\mathrm{SNR}_{\mathrm{th}}\right) \mathrm{SNR}_{\mathrm{th}} = \frac{1}{N}, \tag{S9.15}$$

which yields a quadratic equation for the optimal threshold value for SNR:

$$NT\mathrm{SNR}_{\mathrm{th}}^2 - T\mathrm{SNR}_{\mathrm{th}} - 2 = 0. \tag{S9.16}$$

This yields an optimal threshold:

$$\mathrm{SNR}_{\mathrm{th}} = \frac{1 + \sqrt{1 + 8\frac{N}{T}}}{2N}. \tag{S9.17}$$

We can plug $s_{\mathrm{th}}^2 = N\,\mathrm{SNR}_{\mathrm{th}}\,\hat{\sigma}^2$ into Equation (S8.6) for the empirical sample eigenvalues in order to find the closed-form sample eigenvalue cut-off:

$$\hat{d}_{\mathrm{th}} = \hat{\sigma}^2 \left(1 + \frac{2}{1 + \sqrt{1 + 8q}}\right) \left(q + \frac{1 + \sqrt{1 + 8q}}{2}\right), \tag{S9.18}$$

with $q \equiv \frac{N}{T}$.

Interestingly, we can rescale the numbers of neurons/trials by the SNR:

$$n \equiv N \cdot \mathrm{SNR} \tag{S9.19}$$
$$t \equiv T \cdot \mathrm{SNR}, \tag{S9.20}$$

and then the equation for critical curves in $n - t$ space is:

$$(n-1)t = 2. \tag{S9.21}$$

So the critical SNR is hyperbolic in $N$ and $T$, up to a shift in $N$.

#### S9.2 Masked Autoencoder with Non-negligible Cross-Overlaps

In real data we observe some non-trivial cross-alignment between distinct latent modes. We therefore recalculate the held-out neuron prediction variance explained without assuming purely diagonal alignment, i.e. we allow for

$$\mathcal{A}_{jk} = \left(\hat{\boldsymbol{u}}_j^T \boldsymbol{u}_k\right)^2 \neq 0. \tag{S9.22}$$

Suppose that

$$\boldsymbol{r} = US\boldsymbol{y} + \boldsymbol{z}, \tag{S9.23}$$

where $U$ is $N \times K$ with $N \gg K$, $S$ is diagonal $K \times K$ with $s_k = \sqrt{N\mathrm{SNR}_k}$ along the diagonal.

As defined at the beginning of this section, let $\hat{U}$ be the $N \times \hat{K}$ matrix consisting of the first $\hat{K}$ empirical modes, and $\hat{\boldsymbol{u}}_0^T$ the held-out row of $\hat{U}$. The prediction for masked neuron $r_0 = \boldsymbol{u}_0^T S\boldsymbol{y} + \boldsymbol{z}_0$ is

$$\hat{r}_0 = \hat{\boldsymbol{u}}_0 \hat{U}^T \boldsymbol{r} = \hat{\boldsymbol{u}}_0 \hat{U}^T US\boldsymbol{y} + \eta_0, \tag{S9.24}$$

with $\eta_0 = \hat{\boldsymbol{u}}_0^T \hat{U}^T \boldsymbol{z} \sim \mathcal{N}\left(0, \frac{K}{N}\right)$.

The signal part of the prediction is

$$\hat{r}_{\mathrm{signal}} \equiv \hat{\boldsymbol{u}}_0 \hat{U}^T USy = \sum_{j=1}^{\hat{K}} \sum_{k=1}^{K} \hat{u}_{0_j} \left(\hat{\boldsymbol{u}}_j^T \boldsymbol{u}_k\right) s_k y_k. \tag{S9.25}$$

Averaging over held-out neurons and noise:

$$\left\langle \hat{r}_{\mathrm{signal}}^2 \right\rangle = \sum_{j,j'=1}^{\hat{K}} \sum_{k,k'=1}^{K} \left\langle \hat{u}_{0_j} \hat{u}_{0_{j'}} \left(\hat{\boldsymbol{u}}_j^T \boldsymbol{u}_k\right) \left(\hat{\boldsymbol{u}}_{j'}^T \boldsymbol{u}_{k'}\right) s_k s_{k'} y_k y_{k'} \right\rangle. \tag{S9.26}$$

By independence of the columns of $U$, a term can only contribute if $k = k'$. Similarly, due to the factor of $\hat{u}_{0_j} \hat{u}_{0_{j'}}$, a term will be negligible unless $j = j'$. Since $\left(\hat{\boldsymbol{u}}_j^T \boldsymbol{u}_k\right)^2 = \mathcal{A}_{jk}$, $s_k^2 = N\mathrm{SNR}_k$, $\left\langle \hat{u}_{0_j}^2 \right\rangle = \frac{1}{N}$ and $\left\langle y_k^2 \right\rangle = 1$, we obtain

$$\left\langle \hat{r}_{\mathrm{signal}}^2 \right\rangle = \sum_{k=1}^{K} \sum_{j=1}^{\hat{K}} \mathcal{A}_{jk} \mathrm{SNR}_k. \tag{S9.27}$$

As for the correlation with the signal, $r_{\mathrm{signal}} = \boldsymbol{u}_0^T S\boldsymbol{y}$:

$$\left\langle \hat{r}_{\mathrm{signal}} r_{\mathrm{signal}} \right\rangle = \sum_{j=1}^{\hat{K}} \sum_{k,k'=1}^{K} \left\langle \hat{u}_{0_j} u_{0_{k'}} \left(\hat{\boldsymbol{u}}_j^T \boldsymbol{u}_k\right) s_k s_{k'} y_k y_{k'} \right\rangle. \tag{S9.28}$$

As before, only terms with $k = k'$ contribute, and we have $\left\langle \hat{u}_{0_j} u_{0_k} \left(\hat{\boldsymbol{u}}_j^T \boldsymbol{u}_k\right) \right\rangle = \mathcal{A}_{jk}$, yielding

$$\left\langle \hat{r}_{\mathrm{signal}} r_{\mathrm{signal}} \right\rangle = \sum_{j=1}^{\hat{K}} \sum_{k=1}^{K} \mathcal{A}_{jk} \mathrm{SNR}_k. \tag{S9.29}$$

The error is

$$\left\langle (\hat{r}_0 - r_0)^2 \right\rangle = \left\langle r_{\text{signal}}^2 \right\rangle + \left\langle \hat{r}_{\text{signal}}^2 \right\rangle - 2 \left\langle r_{\text{signal}} \hat{r}_{\text{signal}} \right\rangle + 1 + \frac{\hat{K}}{N}, \tag{S9.30}$$

where $1 + \frac{\hat{K}}{N}$ comes from the noise terms, $z_0$ and $\eta_0$.

We have $\left\langle r_{\text{signal}}^2 \right\rangle = \sum_{k=1}^{K} \text{SNR}_k$, so we have for the error

$$\left\langle (\hat{r}_0 - r_0)^2 \right\rangle = \sum_{k=1}^{K} \text{SNR}_k \left( 1 - \sum_{j=1}^{\hat{K}} \mathcal{A}_{jk} \right) + 1 + \frac{\hat{K}}{N}. \tag{S9.31}$$

The variance explained is then

$$R^2 = \frac{\sum_{k=1}^{K} \text{SNR}_k \left( \sum_{j=1}^{\hat{K}} \mathcal{A}_{jk} \right) - \frac{\hat{K}}{N}}{\sum_{k=1}^{K} \text{SNR}_k + 1}. \tag{S9.32}$$

Interestingly, we find that the sum of alignments from all empirical eigenvectors into each true signal eigenvector enters the calculation:

$$\sum_{j=1}^{\hat{K}} \mathcal{A}_{jk} = \sum_{j=1}^{\hat{K}} \left( \hat{\boldsymbol{u}}_j^T \boldsymbol{u}_k \right)^2. \tag{S9.33}$$

This is a measure of subspace alignment, $\left\| \boldsymbol{u}_k^T \hat{U} \right\|^2$, i.e. the norm squared of the projection of the $k$th signal eigenvector into the entire $\hat{K}$-dim subspace of the sample eigenvectors.

The appearance of the full subspace alignment partially explains why the cross-alignment-free expression for $R^2$ in Equation (S9.13) predicts $R^2$ reasonably well, even in circumstances where individual overlaps $\mathcal{A}_{jk}$ deviate from the theory (which predicts $\mathcal{A}_{jk} = \delta_{jk} \mathcal{A}_k$): By summing over a range of data subspaces, the subspace overlap recovers signal overlap that has rotated into nearby subspaces. Then, roughly,

$$\left\| \boldsymbol{u}_k^T \hat{U} \right\|^2 = \sum_{j=1}^{\hat{K}} \mathcal{A}_{jk} \approx \mathcal{A}_k, \tag{S9.34}$$

whenever $k \leq \hat{K}$, and Equation (S9.32) approximately reduces to Equation (S9.13). See Section S8.3.4 for more discussion of the subspace overlap.

Thus, while quantifying individual cross-alignments for finite-$N, T$ is significantly more involved Bun et al. (2018); Landau et al. (2023), the cross-alignment-free expression (S9.13) is found to yield a reasonable match to experiments for modest $N, T$.

#### S9.3 Scaling laws for masked linear autoencoders

We found in Section Low-Rank Fit that the three datasets that are well fit by the low-rank plus noise model (Mouse MP, Mouse FOV, and Monkey) all exhibit inferred SNRs that obey an approximate power-law:

$$\text{SNR}_k = \frac{\text{SNR}_1}{k^\alpha}. \tag{S9.35}$$

Given these SNRs, the equation for threshold ($\mathcal{A}\text{SNR} = \frac{1}{N}$) becomes

$$2k^{2\alpha} + T\,\text{SNR}_1 k^\alpha - NT\,\text{SNR}_1^2 = 1. \tag{S9.36}$$

We find the positive quadratic root for the threshold index $k_{\text{th}}^\alpha$

$$k_{\text{th}}^\alpha = \frac{T\,\text{SNR}_1 \left( -1 + \sqrt{1 + 8\frac{N}{T}} \right)}{4}, \tag{S9.37}$$

and then take the $\frac{1}{\alpha}$th power to find the predictive dimensionality as displayed in Equation (11).

To find simplified scaling laws we expand first in the $T \gg N$ limit. In this limit we have $\sqrt{1+8\frac{N}{T}} \approx 1+4\frac{N}{T}$ which yields $k_{\rm th}^{\alpha} \approx N\,\mathrm{SNR}_1$, i.e.

$$\hat{D}_{\rm pred} \approx (N\,\mathrm{SNR}_1)^{\frac{1}{\alpha}}\,. \tag{S9.38}$$

This is consistent with the observation that in the large T limit, $\mathcal{A}=1$ for all signal modes and the resulting threshold for predictivity is $\mathrm{SNR}_{\rm th} = \frac{1}{N}$, and it is the consequence of irreducible test noise: in order to contribute positively to prediction, each mode must overcome the independent neural noise.

In the other extreme, $N \gg T$ , we have simply $\sqrt{1+8\frac{N}{T}} \approx \sqrt{8\frac{N}{T}}$, and this becomes the leading order term. The result is to leading order $k^{\alpha} \approx \sqrt{\frac{NT}{2}}\mathrm{SNR}_1$, i.e.

$$\hat{D}_{\rm pred} \approx \left(\tfrac{NT}{2}\,\mathrm{SNR}_1^2\right)^{\frac{1}{2\alpha}}\,. \tag{S9.39}$$

### S10 Review of Random Matrix Theory Derivations

Before reviewing the derivation of key low-rank random matrix theory results, we present a few useful transforms and identities.

First, the *matrix resolvent* of a square matrix $A$ is a matrix-valued function of a complex argument defined as

$$G_A(z) \equiv (z-A)^{-1}\,. \tag{S10.1}$$

The normalized trace of the resolvent is a complex-valued function of a complex argument, called the *Stieltjes transform*:

$$g_A(z) \equiv \tau\left[G_A(z)\right] = \tau\left[(z-A)^{-1}\right]. \tag{S10.2}$$

Writing $A$ in its eigenbasis one sees this is equal to

$$g_A(z) = \frac{1}{N}\sum_{k=1}^{N}\frac{1}{z-\lambda_k}. \tag{S10.3}$$

where $\lambda_k$ are the eigenvalues of $A$.

In the large $N$ limit, one can recover the eigenvalue density from the Stieltjes transform:

$$\rho_W(x) = \frac{1}{\pi}\lim_{\eta\to 0^+}\mathrm{Im}\left[g_W(x-i\eta)\right]. \tag{S10.4}$$

From the eigenvalue density, the Stieltjes transform is obtained via $g_A(z) = \int dx \frac{\rho_A(x)}{z-x}$.

A closely related transform is the *t-transform*:

$$t_A(z) \equiv \tau\left[\frac{A}{z-A}\right] = z g_A(z) - 1. \tag{S10.5}$$

$t_A(z)$ can be interpreted as a generating series for the trace-moments of $A$ which, under the assumption that the moments grow at most exponentially, converges for all sufficiently large $|z|$:

$$t_A(z) = \sum_{n=1}^{\infty}\frac{\tau\left[A^n\right]}{z^n}. \tag{S10.6}$$

Finally, we'll make use of the S-transform which is defined via the functional inverse, $t^{-1}(x)$ of the t-transform (i.e. $t(t^{-1}(x)) = x$ where such a function exists):

$$S_A(x) = \frac{1+x}{x t_A^{-1}(x)}. \tag{S10.7}$$

In this paper we apply these transforms to noise matrices of the form $W = \frac{1}{T}XX^T$ where $X$ has i.i.d. entries with finite fourth moments. For such a matrix, the asymptotic eigenvalue density is given by the Marchenko-Pastur (MP) distribution:

$$\rho_W(x) = \frac{\sqrt{\left(\hat{d}_+ - x\right)\left(x - \hat{d}_-\right)}}{2\pi q x}, \qquad x \in [\hat{d}_-, \hat{d}_+], \tag{S10.8}$$

where $\hat{d}_\pm$ are the lower and upper bounds of the eigenvalue distribution, given by $\hat{d}_\pm = \left(1 \pm \sqrt{q}\right)^2$

The MP distribution has Stieltjes transform

$$g_W(z) = \frac{z + q - 1 - \sqrt{z - \hat{d}_-}\sqrt{z - \hat{d}_+}}{2qz}, \tag{S10.9}$$

and t-transform

$$t_W(z) = \frac{z - q - 1 - \sqrt{z - \hat{d}_-}\sqrt{z - \hat{d}_+}}{2q}. \tag{S10.10}$$

##### S10.1 Review Derivation: Outlier Eigenvalues

We review the derivation for the empirical outlier eigenvalue of $\hat{C} = \sqrt{C}W\sqrt{C}$, given that $C$ is a low-rank signal covariance plus identity. For simplicity we assume a rank-one signal so that $C = s^2\boldsymbol{u}\boldsymbol{u}^T + I$.

An eigenvalue, $\hat{d}$, of $\hat{C}$ satisfies $\det\left(\hat{d}I - \hat{C}\right) = 0$. Inside the determinant we can multiply from the left by $\sqrt{C}$ and from the right by $\sqrt{C}^{-1}$, and the condition becomes

$$0 = \det\left(\hat{d}I - CW\right) \tag{S10.11}$$

$$= \det\left(\hat{d}I - W - s^2\boldsymbol{u}\boldsymbol{u}^T W\right). \tag{S10.12}$$

Now we write $M = \hat{d}I - W$ and use the matrix determinant lemma to arrive at

$$0 = \det(M) \cdot \left(1 - s^2\boldsymbol{u}^T W M^{-1}\boldsymbol{u}\right). \tag{S10.13}$$

Now we recognize that computing the quadratic form of a generic matrix $A$ with a random, normalized vector $\boldsymbol{u}$ yields the normalized trace: $\boldsymbol{u}^T A\boldsymbol{u} \sim \tau[A]$. Therefore we have

$$\boldsymbol{u}^T W M^{-1}\boldsymbol{u} = \boldsymbol{u}^T \frac{W}{\hat{d}I - W}\boldsymbol{u} \to \tau\left[\frac{W}{\hat{d}I - W}\right] = t_W(\hat{d}). \tag{S10.14}$$

This is exactly the scalar t-transform of the matrix $W$.

Thus we finally have that $\hat{d}$ is an outlier eigenvalue of $\hat{C}$ if

$$t_W(\hat{d}) = \frac{1}{s^2}. \tag{S10.15}$$

The inverse of the t-transform only exists above the ceiling of the noise bulk, i.e. for $z > \hat{d}_+$. Specifically, $t_W(z)$ is strictly decreasing with $t_W(z) \to 0^+$ as $z \to \infty$ (the derivative with respect to $z$ is $-\tau\left[W(z - W)^{-2}\right]$ which is negative for all $z > \hat{d}_+$). Thus a solution $\hat{d}$ only exists if $\frac{1}{s^2} < t_W(\hat{d}_+)$, i.e. $s^2 > \frac{1}{t_W(\hat{d}_+)}$

To find the range of $s^2$ for which an outlier exists, we must therefore find $t_W(\hat{d}_+) = \hat{d}_+ g_W(\hat{d}_+) - 1$. From Equation (S10.10), we find that for our case of i.i.d. samples with finite fourth moments, we have $t_W(\hat{d}_+) = \frac{1}{\sqrt{q}}$. Thus, an outlier eigenvalue can only exist if $s^2 > \sqrt{q}$.

Proceeding to find the value of the outlier, $\hat{d}$, the inverse of the t-transform can be written as $t_W^{-1}(z) = \frac{z+1}{zS_W(z)}$, where $S_W$ is the S-transform. In our case, the S-transform is given by $S_W(z) = (1 + qz)^{-1}$. This yields $t_W^{-1}(z) = (1 + \frac{1}{z})(1 + qz) = 1 + q + qz + \frac{1}{z}$

We therefore have, finally, the sample-covariance outlier eigenvalue as a function of the true covariance signal eigenvalue.

$$\hat{d} = t_W^{-1}\left(\frac{1}{s^2}\right) = s^2 + 1 + q + \frac{q}{s^2}, \tag{S10.16}$$

which decomposes into signal $s^2$, noise $1+q$, and a signal-dependent inflation term $\frac{q}{s^2}$.

Note that for $s^2 = \sqrt{q}$, we have $\hat{d} = (1+\sqrt{q})^2$, which is the edge of the noise spectrum.

**Single-Neuron SNR Scaling** To get the scaling with single-neuron SNR we write

$$C = s^2 \boldsymbol{u}\boldsymbol{u}^T + \sigma^2 I, \tag{S10.17}$$

and recall that $s^2 = N\,\mathrm{SNR}\,\sigma^2$.

Following through the same calculation as above, we find that $\hat{d}$ is an outlier if it satisfies $t_W\left(\frac{\hat{d}}{\sigma^2}\right) = \frac{\sigma^2}{s^2}$, which yields

$$\hat{d} = \sigma^2\left(N\,\mathrm{SNR} + 1 + \frac{N}{T} + \frac{1}{T\,\mathrm{SNR}}\right), \tag{S10.18}$$

for $\mathrm{SNR} > \frac{1}{\sqrt{NT}}$, and otherwise $\hat{d} = \sigma^2\left(1+\sqrt{\frac{N}{T}}\right)^2$.

#### S10.2 Review Derivation: Outlier Eigenvector Alignment

We provide an elementary derivation for the alignment between eigenvectors of the true and sample covariances. The result was established rigorously by Paul (2007) and with more generality by Benaych-Georges and Nadakuditi (2012) via different methods.

Consider again the rank-1 model, $C = s^2\boldsymbol{u}\boldsymbol{u}^T + I$, and the resulting sample covariance, $\hat{C} = \sqrt{C}W\sqrt{C}$, where $W = \frac{1}{T}XX^T$ with $X_{ij}$ standard normal, $N \times T$.

Write the sample covariance in its eigenbasis:

$$\hat{C} = \sum_{k=1}^{N} \hat{d}_k \hat{\boldsymbol{u}}_k \hat{\boldsymbol{u}}_k^T. \tag{S10.19}$$

Then the resolvent matrix of $\hat{C}$ can be written as

$$\hat{G}(z) := \left(z - \hat{C}\right)^{-1} = \sum_{k=1}^{N} \frac{\hat{\boldsymbol{u}}_k \hat{\boldsymbol{u}}_k^T}{z - \hat{d}_k}. \tag{S10.20}$$

If the signal, $s^2$, is sufficiently large such that the leading eigenvalue, $\hat{d}_1$, is a well-separated outlier, then the complex function $\boldsymbol{u}^T\hat{G}(z)\boldsymbol{u}$ has a simple pole at $z = \hat{d}_1$ where the residue is exactly the alignment, $|\boldsymbol{u}^T\hat{\boldsymbol{u}}_1|^2$.

Thus, in the case of an outlier, we can write the alignment between the true covariance eigenvector $\boldsymbol{u}$ and any sample covariance eigenvector, $\hat{\boldsymbol{u}}_k$, as:

$$|\boldsymbol{u}^T\hat{\boldsymbol{u}}_k|^2 = \lim_{z\to\hat{d}_k}\left(z - \hat{d}_k\right)\boldsymbol{u}^T\hat{G}(z)\boldsymbol{u}. \tag{S10.21}$$

The main step to derive the expression for the eigenvector alignment is then to find the resolvent, $\hat{G}(z)$. To do so we will use the Woodbury matrix identity: $(A+VMV^T)^{-1} = A^{-1} - A^{-1}V\left(M^{-1} + V^TA^{-1}V\right)^{-1}V^TA^{-1}$.

First we write

$$\sqrt{C} = \sqrt{I + s^2\boldsymbol{u}\boldsymbol{u}^T} = I + \left(\sqrt{s^2+1} - 1\right)\boldsymbol{u}\boldsymbol{u}^T, \tag{S10.22}$$

and then we can rewrite

$$\hat{C} = \sqrt{C}W\sqrt{C} = W + \alpha VMV^T, \tag{S10.23}$$

where we have defined the $N \times 2$ matrix

$$V := \begin{bmatrix} \boldsymbol{u} & W\boldsymbol{u} \end{bmatrix}, \tag{S10.24}$$

the $2 \times 2$ matrix

$$M := \begin{pmatrix} \alpha & 1 \\ 1 & 0 \end{pmatrix}, \tag{S10.25}$$

and finally the scalar

$$\alpha := \sqrt{s^2 + 1} - 1. \tag{S10.26}$$

Now we can apply the Woodbury matrix identity to find

$$\hat{G}(z) = G(z) + \alpha G(z) V \underbrace{\left(M^{-1} - \alpha V^T G(z) V\right)}_{Q(z)}{}^{-1} V^T G(z), \tag{S10.27}$$

where $G(z)$ is the resolvent of $W$:

$$G(z) := (z - W)^{-1}, \tag{S10.28}$$

and we've introduced

$$Q(z) \equiv M^{-1} - \alpha V^T G(z) V. \tag{S10.29}$$

We next seek to find the inverse, $Q(z)^{-1}$.

First, we have $M^{-1} = \begin{pmatrix} 0 & 1 \\ 1 & -\alpha \end{pmatrix}$.

We observe that $\boldsymbol{u}^T G(z) \boldsymbol{u} = g(z)$, while $\boldsymbol{u}^T G(z) W \boldsymbol{u} = t(z)$, and $\boldsymbol{u}^T W G(z) W \boldsymbol{u} = z t(z) - 1$, where $g(z)$ and $t(z)$ are the Stieltjes and t-transforms of $W$, respectively. Thus,

$$Q(z) = \begin{pmatrix} -\alpha g(z) & 1 - \alpha t(z) \\ 1 - \alpha t(z) & -\alpha z t(z) \end{pmatrix}. \tag{S10.30}$$

To find $Q(z)^{-1}$ we first find

$$\det\left[Q(z)\right] = \alpha^2 t(z) \left[z g(z) - t(z)\right] + 2\alpha t(z) - 1 \tag{S10.31}$$

$$= \alpha(\alpha + 2) t(z) - 1, \tag{S10.32}$$

where the last equality follows from the identity $t(z) = z g(z) - 1$.

We can now proceed to find $\boldsymbol{u}^T \hat{G}(z) \boldsymbol{u}$ from Equation (S10.27). We first simplify the formula by observing that $\boldsymbol{u}^T G(z) V = \begin{bmatrix} g(z) & t(z) \end{bmatrix}$.

For notational simplicity, we temporarily suppress the dependence on $z$. We obtain

$$\boldsymbol{u}^T \hat{G} \boldsymbol{u} = g + \frac{\alpha}{\alpha\left(\alpha + 2\right) t - 1} \begin{bmatrix} g & t \end{bmatrix} \begin{pmatrix} -\alpha t z & \alpha t - 1 \\ \alpha t - 1 & -\alpha g \end{pmatrix} \begin{bmatrix} g \\ t \end{bmatrix}. \tag{S10.33}$$

To simplify we next examine

$$\begin{bmatrix} g & t \end{bmatrix} \begin{pmatrix} -\alpha t z & \alpha t - 1 \\ \alpha t - 1 & -\alpha g \end{pmatrix} \begin{bmatrix} g \\ t \end{bmatrix} = -\alpha t z g^2 + 2(\alpha t - 1) g t - \alpha g t^2 \tag{S10.34}$$

$$= -\alpha t z g^2 + \alpha g t^2 - 2 g t \tag{S10.35}$$

$$= \alpha g t \underbrace{\left(-z g + t\right)}_{-1} - 2 g t \tag{S10.36}$$

$$= -(\alpha + 2) g t. \tag{S10.37}$$

Thus we have

$$\boldsymbol{u}^T \hat{G}(z) \boldsymbol{u} = g(z) \left(1 + \frac{\alpha\left(\alpha + 2\right) t(z)}{1 - \alpha\left(\alpha + 2\right) t(z)}\right). \tag{S10.38}$$

We recognize that $\alpha(\alpha+2) = \left(\sqrt{s^2+1}-1\right)\left(\sqrt{s^2+1}+1\right) = s^2$, and arrive at

$$\boldsymbol{u}^T\hat{G}(z)\boldsymbol{u} = \frac{g(z)}{1-s^2t(z)}. \tag{S10.39}$$

Finally, we need to compute this expression for the limit as $z$ approaches the outlier eigenvalue, $\hat{d}_1$. We recall that the outlier eigenvalue satisfies $t(\hat{d}_1) = \frac{1}{s^2}$ as derived in the previous section (Section S10.1), so that taking the limit of $z \to \hat{d}_1$ initially yields $\frac{0}{0}$. By applying L'Hopital's rule we finally obtain

$$|\boldsymbol{u}^T\hat{\boldsymbol{u}}_1|^2 = -\frac{g(\hat{d}_1)t(\hat{d}_1)}{t'(\hat{d}_1)}. \tag{S10.40}$$

Up to this point, we have not used the distribution of the noise covariance $W = \frac{1}{T}XX^T$, and so the expression in Equation (S10.40) holds for more or less arbitrary noise distribution, subject to mild regularity conditions.

We now proceed to the case of finite fourth-moment i.i.d. entries studied throughout the paper. We use that $\hat{d}_1 = (1+\frac{1}{s^2})(s^2+q)$, as derived in Section S10.1, together with $t(\hat{d}_1) = \frac{1}{s^2}$ and $g(z) = \frac{t(z)+1}{z}$ to find that $g(\hat{d}_1) = \frac{1}{s^2+q}$.

To find $t'(z)$ we use the relationship to the S-transform defined above via the functional inverse of $t(z)$: $t^{-1}(x) = \frac{x+1}{xS(x)}$. Assigning $x = t(z)$, rearranging, and taking the derivative with respect to $z$ we arrive at

$$t'(z) = \frac{t(z)S\left(t(z)\right)}{1 - zS\left(t(z)\right) - zt(z)S'\left(t(z)\right)}. \tag{S10.41}$$

As mentioned above, the S-transform for i.i.d. Gaussian samples is $S(t) = (1+qt)^{-1}$, which yields

$$S'(t) = -\frac{q}{(1+qt)^2}. \tag{S10.42}$$

Finally, we would like to evaluate for $z = \hat{d}_1$, so we again assign $t(z) = \frac{1}{s^2}$. We have $S\left(t(z)\right) = \frac{s^2}{s^2+q}$, and $S'\left(t(z)\right) = -\frac{qs^4}{(s^2+q)^2}$

By evaluating Equation (S10.40), we have (suppressing $z$-dependence in $g$ and $t$),

$$
\begin{aligned}
|\boldsymbol{u}^T\hat{\boldsymbol{u}}_1|^2 &= g\frac{ztS'(t) + zS(t) - 1}{S(t)} && \text{(S10.43)}\\
&= \frac{1}{s^2+q}\frac{s^2+q}{s^2}\left(-\frac{\hat{d}_1 s^2 q}{(s^2+q)^2} + \frac{\hat{d}_1 s^2}{s^2+q} - 1\right) && \text{(S10.44)}\\
&= \frac{1}{s^2}\left(\frac{\hat{d}_1 s^4}{(s^2+q)^2} - 1\right) && \text{(S10.45)}\\
&= \frac{s^2+1}{s^2+q} - \frac{1}{s^2}, && \text{(S10.46)}
\end{aligned}
$$

where in the last line we applied $\hat{d}_1 = (1+\frac{1}{s^2})(s^2+q)$.

This at last yields

$$|\boldsymbol{u}^T\hat{\boldsymbol{u}}_1|^2 = \frac{s^4-q}{s^4+s^2q}, \tag{S10.47}$$

which is the result established in the random matrix theory literature.

Recall that this expression holds as long as $\hat{d}_1$ is an outlier, which we showed in Section S10.1 requires $s^2 > \sqrt{q}$. At $s^2 = \sqrt{q}$, Equation (S10.47) yields $|\boldsymbol{u}^T\hat{\boldsymbol{u}}_1|^2 = 0$.

**Single-Neuron SNR Scaling** To get the scaling with single-neuron SNR, we write

$$C = \sigma^2 \left( N\,\mathrm{SNR}\boldsymbol{u}\boldsymbol{u}^T + I \right). \tag{S10.48}$$

Since the eigenvector alignment is not influenced by the overall scale factor $\sigma^2$, we can insert $s^2 = N\,\mathrm{SNR}$ into Equation (S10.47) to obtain:

$$|\boldsymbol{u}^T \hat{\boldsymbol{u}}_1|^2 = \frac{\mathrm{SNR}^2 - \frac{1}{NT}}{\mathrm{SNR}^2 + \frac{\mathrm{SNR}}{T}}, \tag{S10.49}$$

for $\mathrm{SNR} > \frac{1}{\sqrt{NT}}$ and $0$ otherwise.

We separate the numerator to find:

$$|\boldsymbol{u}^T \hat{\boldsymbol{u}}_1|^2 = \frac{T\,\mathrm{SNR}}{T\,\mathrm{SNR} + 1} \left( 1 - \frac{1}{NT\,\mathrm{SNR}^2} \right). \tag{S10.50}$$

Thus, the squared alignment increases as a function of $N$, and saturates for fixed $T$ at $\frac{T\,\mathrm{SNR}}{T\,\mathrm{SNR}+1}$.
